# Fetal signatures in the 3D genome of iPSC-derived neurons and their implications for disease modeling

**DOI:** 10.1101/2025.08.03.667702

**Authors:** D.R. Zagirova, A.D. Kononkova, K.V. Morozov, M.N. Molodova, N.S. Vaulin, A.V. Dudkovskaia, P.I. Dozorova, O.I. Efimova, A.V. Tvorogova, K.A. Ulianov, P.E. Khaitovich, S.V. Razin, M.A. Lagarkova, S.V. Ulianov, E.E. Khrameeva

## Abstract

Induced pluripotent stem cells (iPSCs) have revolutionized neuroscience, providing an approach to generate patient-specific neurons for modeling of neurological diseases. However, it remains unclear how closely iPSC-derived neurons replicate the chromatin architecture of authentic brain neurons. Here, we uniformly process datasets for 228 human and 89 mouse Hi-C and Snm3C-seq samples of different cell subtypes merged into 96 high-coverage contact maps used to examine chromatin features ranging from chromatin compartments and topologically associating domains (TADs) to chromatin loops, Polycomb-mediated contacts, and frequently interacting regions (FIREs). We find that iPSC-derived neurons largely retain chromatin state of undifferentiated cells and resemble fetal rather than mature neurons. iPSC-derived neurons exhibit unusually strong compartmentalization, an enrichment of developmental genes at TAD borders, and a marked reduction of long-range repressive Polycomb-mediated contacts that typically silence early fetal programs. Although immature, iPSC-derived neurons offer advantages for modeling interactions between disease-associated SNPs and target genes, as many psychiatric disorders have neurodevelopmental origins. Integrating iPSC-derived and postmortem neuronal datasets therefore provides complementary insights into the chromatin landscape underlying disease-associated interactions. Our study offers a valuable Hi-C resource for the community and provides a detailed comparison of chromatin architecture throughout neuronal maturation, underscoring its importance for validating neuronal models and providing a robust framework for future studies.

## INTRODUCTION

The introduction of induced pluripotent stem cells (iPSCs) has transformed neuroscience research, providing a versatile platform for generating patient-specific neurons and modeling human neurological diseases. iPSC-derived neurons are produced by reprogramming fibroblasts into iPSCs and subsequently differentiating them into neurons (Takahashi et al. 2007). This approach enables studies of neurons that are genetically identical to the donor, offering an unprecedented *in vitro* system to investigate disease mechanisms and drug responses under controlled conditions (Jiao et al. 2013; Li et al. 2018; Mendez et al. 2023).

Generally, the *in vitro* differentiation of iPSCs into neurons resembles the process that occurs *in vivo* (Gevorgyan et al. 2020). The resulting iPSC-derived neurons exhibit many properties of mature neurons, such as proper morphology, synapse formation, and action potential generation, although manifestation of these properties depends strongly on the specific differentiation protocols employed (Jezierski et al. 2022; Koehler et al. 2011; Prè et al. 2014). Emerging evidence suggests that iPSC-derived neurons remain developmentally immature compared to neurons from the adult brain. While iPSC-derived neurons can recapitulate certain disease-related transcriptional signatures observed in patient postmortem neurons (Hoffman et al. 2017; Sandor et al. 2017), transcriptomic comparisons of more generic models indicate that iPSC-derived neurons might represent an earlier, fetal stage of development rather than fully mature neurons of an adult brain (Burke et al. 2020; Hjelm et al. 2013; Hoffman et al. 2017; Paşca et al. 2015; Stein et al. 2014). Single-cell RNA sequencing data further confirm this fetal-like identity, showing that iPSC-derived neurons closely resemble human fetal neurons (Handel et al. 2016). These differences underscore the importance of considering the developmental stage and context when utilizing iPSC-derived neurons, and highlight potential limitations of these cells in modeling adult-onset neurological and neuropsychiatric diseases.

While transcriptomic assays have been invaluable for characterizing iPSC-derived neurons, they alone are not sufficient to fully assess the quality of these cell models. Previous studies emphasize the need to take into account additional modalities, such as cell environment (e.g., spatial transcriptomics) and chromatin openness (e.g., scATAC-seq) (Childs et al. 2022). Despite these insights, to date there are no studies comparing iPSC-derived and adult postmortem neurons at the level of chromatin architecture or other epigenomic features. Existing Hi-C studies have primarily examined developmental transitions within a single experimental system or profiled chromatin architecture across stages of brain development (Ballarino et al. 2022a; Li 2016; Lu et al. 2020; Heffel et al. 2024; Rahman et al. 2023). As chromatin architecture plays a crucial role in gene regulation and cellular function (Bonev et al. 2017; Rowley and Corces 2018), it is essential to examine similarities and differences in the 3D genome organization for evaluating the validity of iPSC-derived neurons as models of mature neurons.

Here, we aim to bridge this critical gap by systematically comparing precursor cells, iPSC-derived neurons, and fetal and adult postmortem neurons across multiple layers of chromatin architecture. To enable this analysis, we assembled the largest repository of uniformly processed neuronal Hi-C datasets, encompassing more than 300 human and mouse samples (including our six newly generated Hi-C maps) spanning adult, fetal, and iPSC-derived cells. We then characterize chromatin architecture across multiple scales – including A/B compartments, TADs, and chromatin loops – and integrate public ChIP-seq data to compare epigenomic landscapes between iPSC-derived and postmortem neurons. We further assess whether adult neurons and iPSC-derived neurons share certain chromatin contacts – specifically those overlapping GWAS SNPs – to evaluate the utility of iPSC-derived neurons for annotating disease-associated GWAS loci linked to diseases of the nervous system (Won et al. 2016).

## RESULTS

### Creating a harmonized Hi-C data repository for human and mouse neuronal cells

Our data collection strategy involved a systematic search of the Gene Expression Omnibus (GEO) database for human and mouse Hi-C neuronal datasets (Figure 1A). In addition to adult brain neurons (designated “postmortem” in human datasets and “adult” - in mouse datasets) and iPSC-derived neurons, we included selected fetal brain samples and various progenitor stages – embryonic stem cells (ESCs), iPSCs, neural progenitor cells (NPCs), and neuroepithelial stem cells (NESs) differentiated from iPSC. We reprocessed all raw FASTQ files *de novo* using a unified bioinformatics pipeline (see Methods), thereby eliminating processing-related batch effects and ensuring strict cross-study comparability. Additionally, we expanded upon our previous work (Pletenev et al. 2024) by incorporating previously unpublished data, including two neuronal and two non-neuronal Hi-C maps from human brain samples. We also applied an identical experimental protocol to produce two Hi-C maps from iPSC-derived neurons, ensuring comparability between postmortem and *in vitro* differentiated neurons.

**Figure 1.**
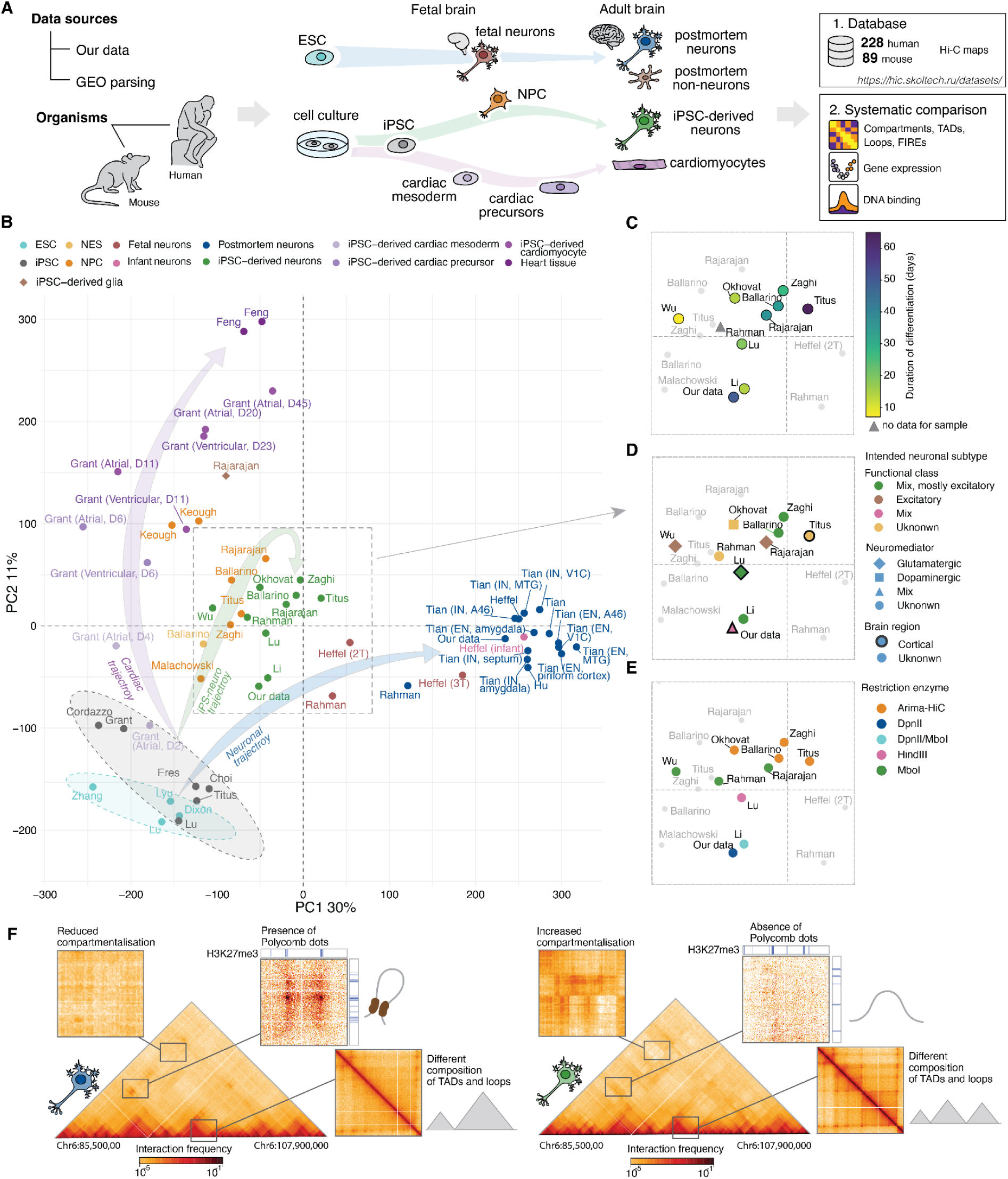
Overview of Hi-C data collection and analysis. A. Schematic representation of the data collection procedure for the downstream analysis and database construction. B. PCA of human Hi-C maps based on genome-wide insulation scores. Ellipses denote the 95% confidence intervals for each group, computed directly from sample distributions. Arrows indicate three major differentiation trajectories, manually added to illustrate developmental progression from pluripotent stem cells: сardiac (violet), iPS-neuro (green) and neuronal (blue). Brain region abbreviations: MTG, middle temporal gyrus (temporal cortex); V1C, primary visual cortex (occipital cortex); A46, prefrontal cortex (Brodmann area 46). EN – excitatory neurons, IN – inhibitory neurons. Detailed regional annotation of postmortem samples is provided in Supplemental Fig. S2. (C–E) Enlarged view of the PCA from panel B highlighting iPSC-derived neurons. Samples are color-coded by duration of the differentiation protocol (C), intended neuronal subtype (functional class, neuromediator, and brain region; D), and restriction enzyme used in the Hi-C protocol (E). F. Examples of Hi-C maps for merged postmortem (left) and iPSC-derived (right) human neurons, illustrating characteristic differences in chromatin organization.

This approach yielded a total of 228 human and 89 mouse Hi-C and Snm3C-seq maps. To our knowledge, this is the largest database of uniformly processed neuronal Hi-C data to date. The complete database, including raw and pre-processed files with diagonal values removed and contact numbers normalized, is publicly accessible at https://www.skoltech.ru/3d-genome-datasets. Applying our inclusion criteria (see Methods) resulted in a final cohort of 62 human and 34 mouse merged Hi-C maps for downstream analyses (Table 1; full metadata for all collected samples are provided in Supplemental Table S1).

**Table 1.** An overview of the collected and newly produced human and mouse Hi-C datasets used in the analysis.

| Organism | Group | Number of Hi-C maps | Name |
| --- | --- | --- | --- |
| Human | Postmortem neurons | 15 | Our data, Heffel et al. adult (2024), Hu et al. (2021), Rahman et al. (2023), Tian et al. (2023) |
| Human | iPSC-derived neurons | 10 | Our data, Ballarino et al. (2022), Li et al. (2019), Lu et al. (2020), Rahman et al. (2023), Rajarajan et al. (2018), Titus et al. (2023), Wu et al. (2021b), Zaghi et al. (2023), Okhovat et al. (2023) |
| Human | Fetal neurons | 4 | Heffel et al. 2T (2024), Heffel et al. 3T (2024), Heffel et al. infant (2024), Rahman et al. (2023) |
| Human | iPSC | 6 | Lu et al. (2020), Titus et al. (2023), Choi et al. (2020), Grant et al. (2025), Cordazzo Vargas and Shioda (2024), Eres et al. (2019) |
| Human | iPSC-derived NPC | 7 | Ballarino et al. (2022), Rajarajan et al. (2018), Titus et al. (2023), Zaghi et al. (2023), Choi et al. (2020), Malachowski et al. (2023), Keough (2023) |
| Human | Postmortem non-neurons | 8 | Our data, Hu et al. (2021), Rahman et al. (2023), Tian et al. (2023) |
| Human | iPSC-derived glia | 1 | Rajarajan et al. (2018) |
| **Human** | **iPSC-derived**<br>**cardiac**<br>**mesoderm** | 2 | Grant et al. (2025) |
| **Human** | **iPSC-derived**<br>**cardiac**<br>**precursor** | 2 | Grant et al. (2025) |
| **Human** | **iPSC-derived**<br>**cardiomyocyte** | 5 | Grant et al. (2025) |
| **Human** | **Heart tissue** | 2 | Feng et al. (2024) |
| **Mouse** | **ESC** | 6 | Bonev et al. (2017), Bauer (Bauer et al. 2021),<br>Fraser et al. (2015) |
| **Mouse** | **ESC-derived**<br>**NPC** | 6 | Bonev et al. (2017), Bauer (Bauer et al. 2021),<br>Fraser et al. (2015) |
| **Mouse** | **ESC-derived**<br>**neurons** | 7 | Bonev et al. (2017), Fraser et al. (2015) |
| **Mouse** | **Fetal NPC** | 4 | Bonev et al. (2017) |
| **Mouse** | **Fetal neurons** | 5 | Bonev et al. (2017), Kiefer et al. (2024) |
| **Mouse** | **Adult**<br>**neurons** | 6 | Kriukov et al. (2024), Chandrasekaran et al. (2021),<br>Rocks et al. (2022), Jiang et al. (2017) |

To assess global similarities across cell types, we performed principal component analysis (PCA) based on genome-wide chromatin insulation scores (Figure 1B, Supplemental Fig. S1,2). To determine whether technical or protocol-related variables could confound clustering patterns, we annotated each iPSC-derived neuronal sample by differentiation duration (Figure 1C), intended neuronal subtype (Figure 1D, Supplemental Fig. S3,4), Hi-C protocol, and restriction enzyme (Figure 1E, Supplemental Fig. S5). None of these factors produced a clear stratification among iPSC-derived neurons. Therefore, all iPSC-derived neuronal samples were treated as a single group in downstream analyses.

For human Hi-C data, samples were categorized into three distinct differentiation trajectories: *cardiac* (from iPSCs to iPSC-derived cardiac mesoderm, iPSC-derived cardiac progenitors, iPSC-derived cardiomyocytes, heart tissue), *iPS-neuro* (from iPSCs to NES, NPC, and iPSC-derived neurons), and *neuronal* (from ESCs to fetal neurons, and postmortem neurons) (Figure 1B). These trajectories were defined to illustrate the progression of differentiation stages within each lineage.

Within the neuronal trajectory, fetal samples form a continuous bridge, spanning from ESC to adult neurons. Early fetal stages (“2T”, 18 gestational weeks) align closer to precursors, while later stages (“Infant”) approach adult neurons, illustrating progressive chromatin remodeling during physiological maturation. In contrast, the *in vitro* iPS-neuro trajectory appears truncated. iPSC-derived neurons cluster near precursor cells (iPSC, NPCs, NES), with minimal separation from NPCs. This occurs despite the well-documented acquisition of mature neuronal morphology and electrophysiological activity in the same cultures (Bardy et al. 2016; Page et al. 2022), indicating that current protocols fail to recapitulate the late-stage chromatin restructuring that characterizes fully mature neurons from the adult brain, even when transcriptional and functional neuronal features have emerged. The clear separation between iPSC-derived and postmortem neuron clusters remains consistent in higher dimensions as well (Supplemental Fig. S6).

To evaluate whether this deficiency is cell-type-specific, we incorporated Hi-C maps representing cardiomyocyte differentiation into the PCA plot. A similar pattern emerged: cells are distributed sequentially from day-2 cardiac mesoderm to day-45 iPSC-derived cardiomyocytes, yet the terminal cardiomyocyte population remains clearly segregated from the adult heart tissue. As expected, adult heart tissue clusters closer to non-neuronal cells (Supplemental Fig. S1).

Parallel analysis of mouse Hi-C maps reveals a pronounced separation between adult neurons and other cells, with fetal neurons clustering closer to ESC-derived neurons and NPCs (Supplemental Fig. S7, Supplemental Fig. S8, Supplemental Fig. S9). In contrast to the human dataset, murine fetal samples do not link precursors to adult cells, potentially due to their earlier developmental stages (mouse: embryonic day E12–E16, equivalent to mid-gestation; human: gestational week GW18–GW39, corresponding to mid-to-late gestation).

Gene expression PCA in human samples recapitulates the clustering patterns observed at the chromatin level (Supplemental Fig. S10). Non-neuronal lineages form a distinct group, clearly separated from all neuronal populations. Postmortem neurons cluster apart from other groups, although several cerebellar samples partially overlap with fetal neurons, suggesting regional heterogeneity in transcriptional maturation. iPSC-derived neurons remain segregated from postmortem clusters, indicating that the divergence between *in vitro* models and mature neurons observed in chromatin architecture is also reflected at the transcriptomic level (Supplemental Fig. S11).

Notably, gene expression PCA in mice shows sharper cluster separation than chromatin-based PCA, with adult neurons clearly distinguished from all fetal and *in vitro*-derived groups (Supplemental Fig. S12). This observation suggests that changes in gene expression and 3D chromatin organization may occur at different stages or rates during maturation. Overall, across species (human, mouse) and cell lineages (neuronal, cardiomyocyte), our analyses reveal that *in vitro* differentiation generates cells that retain precursor-like chromatin architecture.

### General chromatin properties of postmortem and iPSC-derived human neurons

To gain insights into the global organization of chromatin architecture, we plotted the contact probability between genomic loci (*Pc)* as a function of their separating genomic distance (*s), Pc*(*s*), averaged across cell types for samples of neuronal and iPS-neuro trajectories (Supplemental Fig. S13A,D). Compared to other cell types and neurons specifically, non-neuronal samples show a decrease in local (<3 Mbp) and an increase in distal (>3 Mbp) chromatin interactions, consistent with our previous findings (Pletenev et al. 2024). The *Pc(s)* curves for postmortem and fetal neurons are highly similar, suggesting that higher-order chromatin structure is generally preserved during neuronal development and maturation. In contrast, *Pc(s)* curves for iPSC-derived neurons are steeper in regions corresponding to short-range interactions, mirroring the patterns observed in precursor cells and indicative of a more compact chromatin state. At greater genomic distances, however, *Pc(s)* curves for iPSC-derived neurons converge toward that of both postmortem and fetal neuronal samples. Although we initially examined *Pc(s)* curves separately across iPSC-derived subgroups defined by duration of differentiation protocol, neuronal functional class, intended neuronal subtype, Hi-C protocol, and restriction enzyme (Supplemental Fig. S14), the scaling profiles were consistent across these categories; therefore, all iPSC-derived samples were further analyzed as a single group. In cardiac samples, iPSC-derived cells (mesoderm, precursors, and cardiomyocytes) exhibit similar *Pc(s)* patterns, whereas adult heart tissue shows a distinct profile across both short-and long-range interactions.

To further characterize the general properties of chromatin folding, we examined the scaling slope of *Pc(s)*, with particular attention to two characteristic points: (i) the location of the first local maximum (interpreted as the average chromatin loop length), and (ii) the location of the minimum slope (marking the transition point where local features such as TADs and loops diminish and large-scale features like compartments predominate) (Supplemental Fig. S13A-D) (Polovnikov et al. 2023). We observed that these characteristic points progressively shifted to longer genomic distances during the course of differentiation from iPSCs to iPSC-derived neurons and from iPSCs to cardiomyocytes (Supplemental Fig. S13B,C). In postmortem neurons, the first maximum occurs at a significantly shorter distance than in iPSC-derived neurons (1.6×10^5^ and 2.1×10^5^ kb respectively, Mann–Whitney *U* test, *p* = 0.0001; Supplemental Fig. S13B) and the first minimum – at a longer distance (3.9×10^6^ and 3.6×10^6^ kb respectively, Mann–Whitney *U* test, *p* = 0.00001; Supplemental Fig. S13B), implying a depletion of local contacts and an enrichment of long-range interactions in terminally differentiated neurons.

Together, these differences in contact probability scaling reflect a broad reorganization of chromatin architecture that distinguishes iPSC-derived from postmortem neurons across multiple levels of genome organization (Figure 1F), which are examined in detail in the following sections.

### Chromatin compartments in iPSC-derived neurons exhibit an intermediate state between fetal neurons and NPCs

To compare large-scale chromatin organization changes along neuronal and iPS-neuro trajectories, we analyzed chromatin compartments across cell types. PCA of compartment vectors shows that iPSC-derived neurons have an intermediate chromatin state between NPCs and fetal neurons (Figure 2A). Notably, parental iPSCs form a distinct cluster, while iPSC-derived neurons cluster closer to fetal and adult neurons than to their progenitors, indicating progressive chromatin remodeling during differentiation.

**Figure 2.**
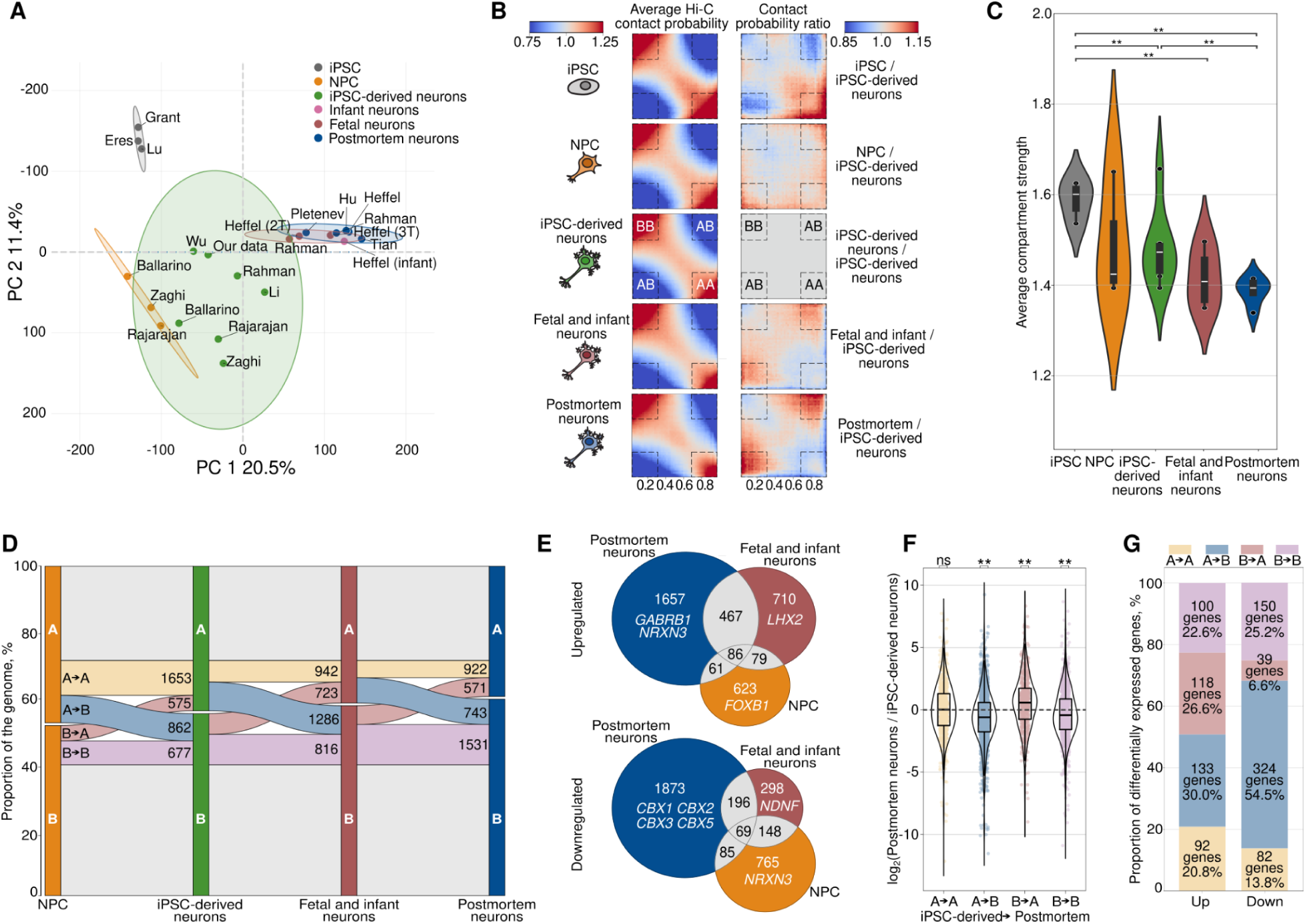
Analysis of compartments and gene expression. **A.** PCA of compartment vectors across analyzed cell groups. Transparent ellipses of matching color represent the 95% confidence intervals. **B.** Saddle plots showing average Hi-C contact probabilities within (AA, BB) and between (AB) compartments (left) and saddle plot ratios relative to iPSC-derived neurons (right). **C.** Average compartment strength. Asterisks indicate Mann-Whitney *U* test *p*-values: ** - *p* < 0.001. **D.** Compartment transitions along the trajectory NPCs → iPSC-derived → fetal/infant → adult postmortem neurons. Colors indicate transition categories: A-to-A (yellow), A-to-B (blue), B-to-A (red), B-to-B (violet), and genomic bins unchanged across all cell groups (grey). **E.** Venn diagram showing overlap among upregulated (top) and downregulated (bottom) genes in NPCs, fetal/infant neurons, and adult postmortem neurons relative to iPSC-derived neurons. **F.** Gene expression ratio log_2_-fold change (postmortem/iPSC-derived) for each type of compartment transitions (iPSC-derived → postmortem neurons: A-to-A, A-to-B, B-to-A, B-to-B). Yellow and violet-genes that retain their compartment, blue and red - genes that change their compartment. Asterisks indicate Wilcoxon test *p*-values: ** - *p* ≤ 10^-10^. **G.** Numbers of up-and downregulated genes defined in panel E within each type of compartment transitions defined in panel F.

Saddle plot ratios comparing each cell group against iPSC-derived neurons show increased inter-compartment interactions along the iPS-neuro *in vitro* differentiation trajectory (iPSCs → NPCs → iPSC-derived neurons), followed by a progressive increase for *in vivo* differentiated fetal and postmortem neurons (Figure 2B). As an increase in contacts between A and B compartments manifests as weaker compartmentalization, these observations suggest a gradually less pronounced compartmentalization pattern along the iPSC → iPSC-derived neurons transition, followed by fetal, infant and postmortem neurons, and our compartment strength analysis fully confirms this progressive decrease (Benjamini-Hochberg adjusted Mann–Whitney *U* test, *p* < 0.001; Figure 2C). Specifically, cis-interactions in an active compartment (AA) gradually decrease, in parallel with an upward trend of AB interactions (Benjamini-Hochberg adjusted Mann–Whitney *U* test, *p* < 0.05; Supplemental Fig. S15A). These observations are consistent with reports of reduced compartmentalization during neuronal development, particularly from mid-to late gestation, and with the reported inverse correlation between compartment strength and distal-to-local contact ratio (Heffel et al. 2024) (Supplemental Fig. S15B).

To investigate the relationship between compartmentalization and cell differentiation state, we analyzed compartment transition dynamics by measuring both the proportion of genomic bins and the number of genes switching between A and B compartments. The proportion of genomic bins in the A compartment, as well as the number of genes, gradually decreases along the trajectory NPCs → iPSC-derived → fetal/infant → adult postmortem neurons, while the B compartment shows an opposite trend (Figure 2D). Consistently, A-to-B transitions outnumber B-to-A transitions in all comparisons along this trajectory (by 49.9% in NPCs → iPSC-derived, by 77.8% in iPSC-derived → fetal/infant, by 30.1% in fetal/infant → adult postmortem; Figure 2D), illustrating a progressive expansion of compartment B as a feature of mature neurons.

To evaluate whether these differences in compartments correspond to transcriptional changes, we compared gene expression across the four neuronal cell groups (Supplemental Fig. S16). Differential expression analysis reveals 2,272 upregulated and 2,223 downregulated genes in postmortem compared to iPSC-derived neurons (Benjamini-Hochberg adjusted Wald test, *p* ≤ 0.01; Supplemental Fig. S16A; Supplemental Table S2). Similar comparisons between fetal/infant and iPSC-derived neurons (1,342 up, and 711 down; Benjamini-Hochberg adjusted Wald test, *p* ≤ 0.01; Supplemental Fig. S16B; Supplemental Table S2), NPCs and iPSC-derived neurons (849 up, 1,067 down; Benjamini-Hochberg adjusted Wald test *p* ≤ 0.01; Supplemental Fig. S16C, Supplemental Table S2) reveal numerous shared and unique differentially expressed genes (DEGs), including key neuronal regulators such as *LHX2, EOMES, NRXN3, EBF1, FOXB1* (Figure 2E). Notably, out of 920 unique human neurodevelopmental genes (defined in the Gene Ontology term GO:0048666 – neuron development), 71.7% (n=660) show comparable expression in iPSC-derived and postmortem neurons (i.e., demonstrate no statistically significant differences). These genes include important transcription factors *FOXG1, RUNX3, NEUROG3, DLX5* that are normally silenced in mature neurons. Concurrently, the altered compartment strength in iPSC-derived neurons aligns with upregulation of genes encoding core heterochromatin binding proteins (*CBX1, CBX3, CBX5*). Beyond its role in B compartment formation (Zenk et al. 2021), the CBX3 protein also promotes axonal and dendritic development (Huang et al. 2017). Likewise, *HDAC2*, which drives chromatin condensation in cultured mouse fetal erythroblasts (Ji et al. 2010), is more highly expressed in iPSC-derived than in postmortem neurons (Supplemental Table S2). Together, these findings reveal that DEGs between iPSC-derived and postmortem neurons include genes important for both neuronal development and chromatin structuring.

Gene Ontology enrichment analysis of DEGs, tested against the background of all expressed genes, reveals consistent functional categories across all comparisons (adult postmortem vs iPSC-derived, fetal/infant vs iPSC-derived, NPCs vs iPSC-derived; Supplemental Fig. S17). Terms related to membrane potential regulation and ion transport appear in several comparisons, becoming more and more enriched along the trajectory NPCs → iPSC-derived → postmortem neurons. At the same time, enrichment of epithelium morphogenesis and embryonic organ development terms may be a consequence of residual expression of neurodevelopmental genes, suggesting that iPSC-derived neurons exhibit an expression profile intermediate between NPCs and postmortem neurons.

To further explore the relationship between compartmentalization and transcription, we calculated the corresponding expression log_2_-fold changes for each compartment transition category (A-to-A, A-to-B, B-to-A, B-to-B) within iPSC-derived → postmortem neuron comparison. Gene expression changes positively correlate with compartment dynamics (Figure 2E,F). Notably, all transition groups except A-to-A show highly significant gene expression changes. These results are consistent with observed compartment strength alterations (Figure 2C; Supplemental Fig. S15A), as well as numbers of up-and downregulated genes within each type of compartment transitions (Figure 2G).

Collectively, these findings indicate that neuronal maturation involves extensive chromatin reorganization, marked by a progressive expansion of transcriptionally inactive B compartment. The described differences in compartmentalization and gene expression suggest that iPSC-derived neurons differ from postmortem neurons and have an intermediate chromatin organization between fetal/infant neurons and NPCs.

### Differential TAD border strength and structural alterations in iPSC-derived neurons indicate developmental characteristics

At a finer scale, chromatin organization is characterized by TADs, which coordinate key regulatory interactions (Dixon et al. 2012; Rowley and Corces 2018). PCA of TAD border strength highlights distinct cell groupings: postmortem and fetal neurons cluster together, iPSC-derived neurons largely overlap with NPCs, and undifferentiated iPSCs form a separate cluster, distinct from neuronal populations (Figure 3A). Direct comparison of average TAD border profiles indicates that iPSC-derived neurons are more similar to NPCs than to fetal or port-mortem neurons (Figure 3B). At TAD borders, we also examined “stripe” features, corresponding to cohesin-mediated loop extrusion tracks that appear to be, on average, more pronounced in iPSC-derived neurons (Supplemental Fig. S18A,B). To simplify downstream analysis, we concentrated on the terminal states of neuronal and iPS-neuro trajectories (Figure 1B): specifically, postmortem neurons and iPSC-derived neurons.

**Figure 3.**
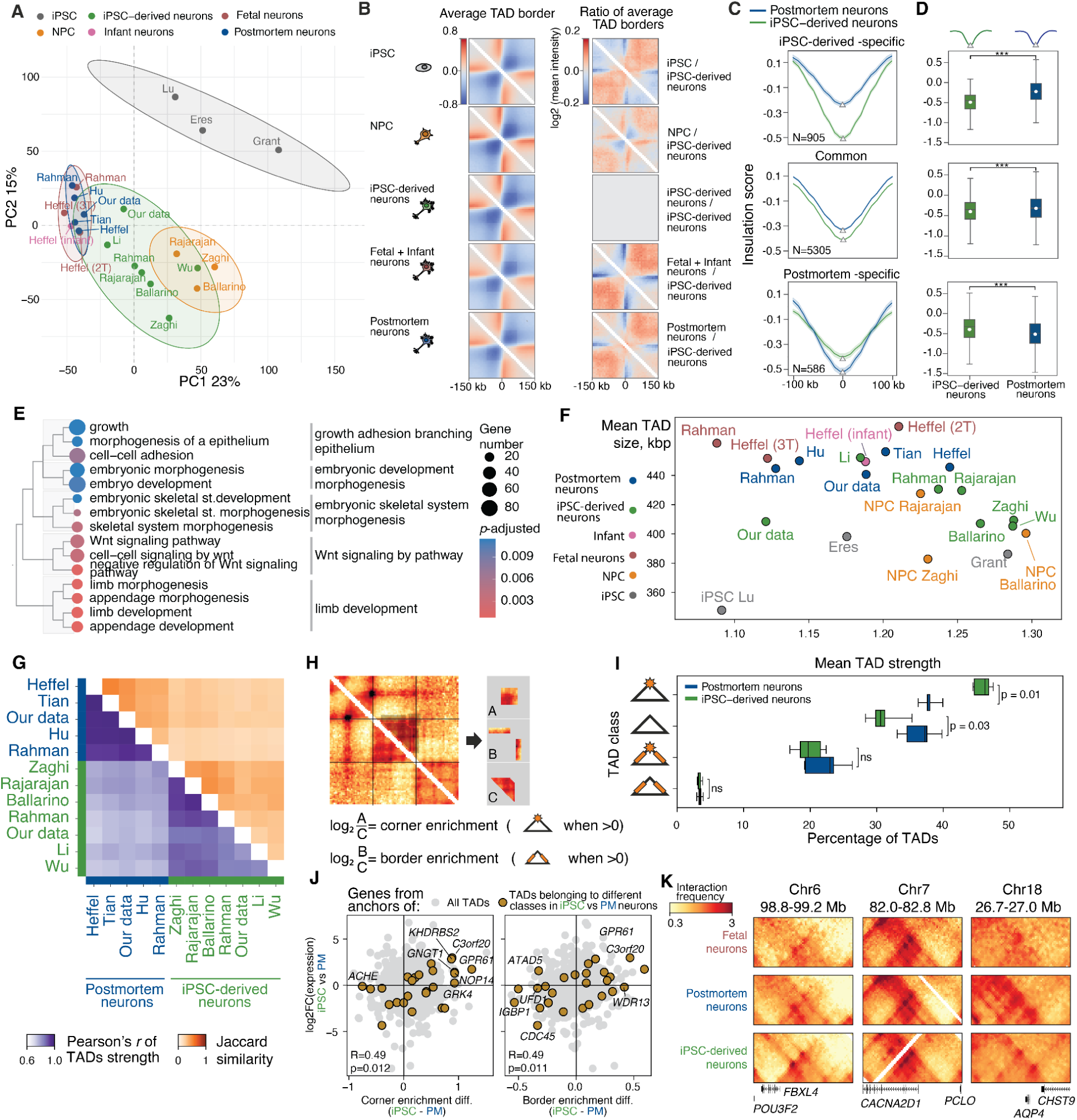
Analysis of TADs. A. PCA of TAD border strength across analyzed groups. B. Average TAD borders for selected cell types (left panel), and the ratio of these average TAD borders relative to iPSC-derived neurons (right panel). C. Insulation score profiles at common TAD borders (middle panel), and at differential borders: iPSC-derived neuron-specific (top panel) and postmortem neuron-specific (bottom panel). D. Box plots showing the distribution of insulation scores across borders. For each border, the insulation score was obtained from the genomic bin with the minimum value within the border region. Asterisks indicate Mann-Whitney *U* test *p*-values: *** - *p* < 0.001. Е. GO terms enrichment for genes located at TAD borders specific to iPSC-derived neurons. F. Relationship between mean TAD strength and TAD size across samples. G. Jaccard similarity of TADs with common borders (upper triangle) and Pearson’s correlation of TAD strength (lower triangle), with an adjacent barplot showing the total number of TADs per map. H. Schematic representation of a procedure to extract TAD features. I. Percentage of TADs exhibiting corner and/or border enrichment in postmortem compared to iPSC-derived neurons. J: Scatterplot of log_2_FC of gene expression versus difference in corner and border enrichment scores between iPSC-derived and postmortem neurons (R - Pearson’s correlation coefficient, p - *p*-value for testing non-correlation). K: Representative Hi-C maps (observed/expected) from postmortem and iPSC-derived neurons (all samples within the group merged together) illustrating structural differences.

Differential analysis of TAD borders revealed two distinct groups: borders specific to iPSC-derived neurons and borders specific to postmortem neurons (Supplemental Fig. S 19). Although insulation around post mortem-specific borders was significantly decreased (Mann–Whitney *U* test, p = 3.67×10-27; Figure 3D), the aggregate amplitude of change remained modest (Figure 3C,D). By contrast, iPSC-derived-specific borders demonstrated a very pronounced difference in insulation strength. Further, we found that these iPSC-neuron-specific borders exhibited similarly high insulation strength in both NPCs and iPSC-derived neurons (Supplemental Fig. S20), suggesting that these borders may represent residual NPC features that persist in iPSC-derived neurons.

Genes located at iPSC-derived neuron-specific borders were significantly enriched for developmental processes (Figure 3E). In contrast, postmortem-specific borders showed no statistically significant GO enrichment, although they did include relevant genes such as *GABRB2* (implicated in synapse formation), underscoring potential biological relevance of defined postmortem-specific borders (Supplemental Fig. S21A,B). The relatively lower number of postmortem-specific TAD borders and absence of GO category enrichment are likely due to the high variability of TAD border strength observed in iPSC-derived neurons (Welch’s *t*-test, p = 3.23×10-17; Supplemental Fig. S22A). For example, some borders that are consistently strong in all postmortem samples show variable or absent strength in iPSC-derived neurons, which limits our statistical power to define them as postmortem-specific. This is illustrated by *CSMD2*, a key neuronal gene, which exhibits strong TAD borders in postmortem samples but variable or even absent borders in iPSC-derived neurons (Supplemental Fig. S22B-D). These findings suggest that one key difference between iPSC-derived and postmortem neurons may lie in the presence of additional borders, representing residual NPC features that persist in iPSC-derived neurons and contain development-related genes; moreover, these borders may not be lost during *in vitro* maturation.

Using unified preprocessing and TAD calling methods, we observed that TADs in iPSC-derived neurons are generally smaller and exhibit higher domain scores (TAD strength) than those in postmortem neurons (Figure 3F). Postmortem samples show greater similarity to each other and to infant samples than to any iPSC-derived samples (Figure 3G; Supplemental Fig. S23A). In contrast, iPSC-derived neurons share a larger fraction of TADs with NPCs, consistent with their developmental origin. We further classified TADs as corner-or border-enriched (Figure 3H) and observed significant differences in the proportions of these classes between iPSC-derived and postmortem neurons (Mann–Whitney *U* test; Figure 3I). Differences in the enrichment values correlate with gene expression changes for genes located at TAD boundaries that consistently belong to different classes in all iPSC-derived and postmortem samples (Figure 3J). For example, the neuronal genes *POU3F2* and *FBXL4* show reduced expression in iPSC-derived neurons and reside within a TAD displaying stronger corner enrichment in iPSC-derived samples (Figure 3K; Supplemental Fig. S23B).

This indicates that iPSC-derived neurons have not only additional TAD borders, but also an altered inner structure of TADs. Specifically, their TADs are denser and exhibit more enriched corner intensity compared to postmortem neurons. These features of iPSC-derived TADs make them more similar to TADs from stem-like cells.

### Developmental gene-associated chromatin loops distinguish iPSC-derived from postmortem neurons

We further analyzed the fine-scale chromatin organization, focusing on chromatin loops. PCA of loop intensities reveals a clustering pattern consistent with other chromatin features: postmortem neurons are closely aligned with fetal neurons, while iPSC-derived neurons occupy an intermediate position between these two groups and NPCs (Figure 4A). In line with results for TAD borders, comparison of average loop profiles shows that iPSC-derived neurons are most similar to NPCs compared to fetal and postmortem neurons or iPSC (Figure 4B). Direct comparison between iPSC-derived and postmortem neurons identified both common and cell type-specific chromatin loops (Supplemental Fig. S24A-H). Notably, common loops exhibit slightly higher intensities in iPSC-derived neurons (Supplemental Fig. S24D,E). Correlation analysis and assessment of the fraction of shared loops further demonstrate clear segregation between postmortem and iPSC-derived neurons, with stronger intra-group heterogeneity among iPSC-derived samples (Figure 4C), which resembles results for TAD borders. Although the Zaghi dataset does not cluster with other iPSC-derived samples based on loop intensity correlations, it exhibits highly similar loop positions and higher-order chromatin organization, supporting its inclusion in the analysis.

**Figure 4.**
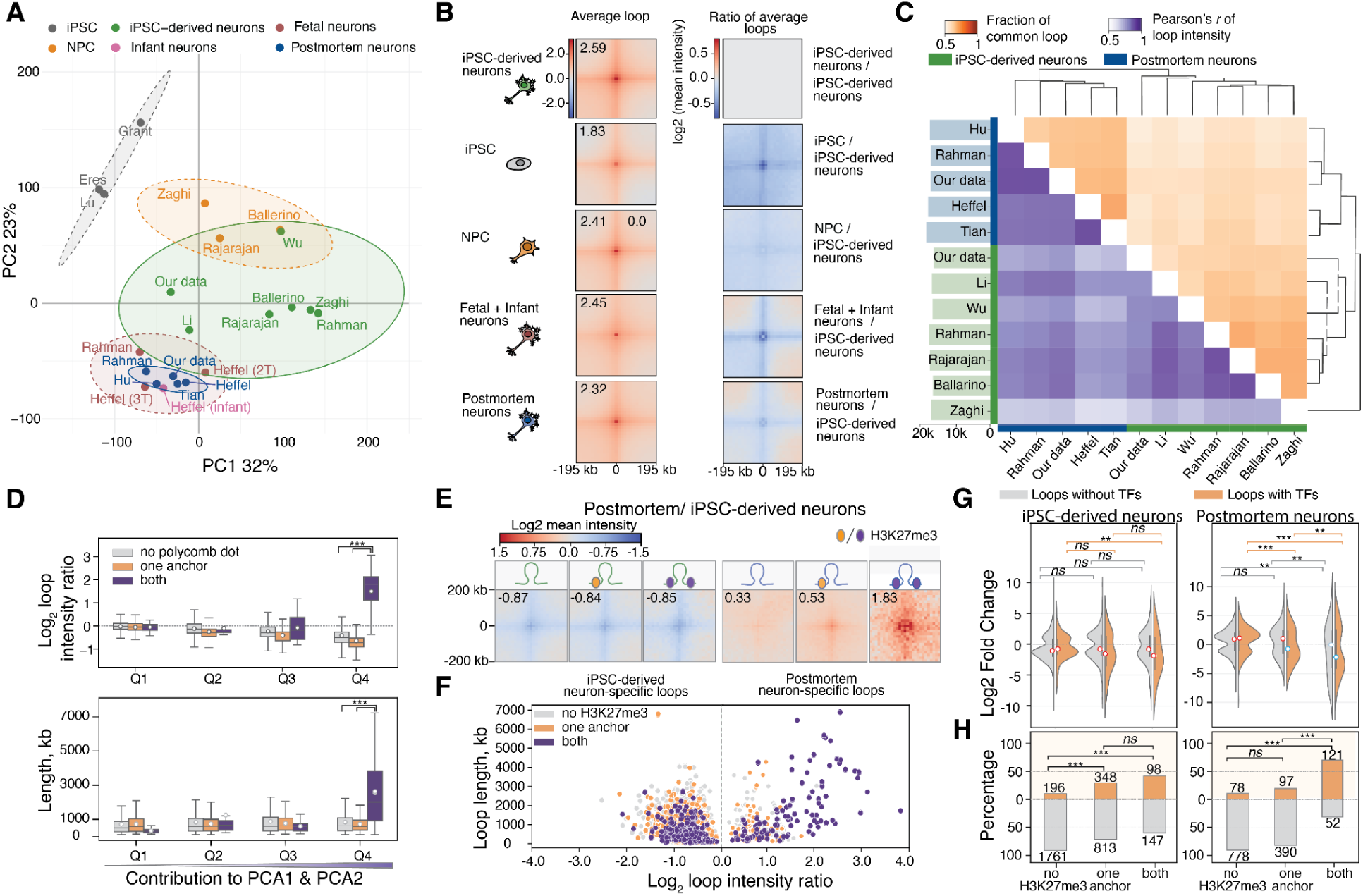
Analysis of chromatin loops. A. PCA of loop intensities across analyzed groups. B. Average chromatin loop intensities for each group. C. Clustermap showing the fraction of common loops (upper triangle) and Pearson’s correlation of loop intensities (lower triangle), with a bar plot indicating the number of unique loops per sample. D. Box plots displaying loop categories (Q1-Q4) partitioned based on their contribution to PC1 and PC2 separation and on the presence or absence of PcG-contacts. The top plot shows loop intensity ratios, and the bottom plot displays loop lengths in kilobases. Asterisks indicate Welch’s *t*-test *p*-values: *** - *p* < 0.001. E. Average loop intensities for loop categories partitioned based on the H3K27me3 presence in one or both anchors. F. A scatter plot of loop length versus the log_2_ loop intensity ratio (iPSC-derived/postmortem neurons). G. Log_2_ gene expression fold change (postmortem/iPSC-derived) across chromatin loops with or without H3K27me3 in one or both anchors, categorized by the presence of transcription factor (TF) genes in iPSC-derived (left) and postmortem (right) neurons. Positive values indicate higher expression in postmortem neurons; negative values indicate higher expression in iPSC-derived neurons. Asterisks indicate Welch’s *t*-test *p*-values: ns - not significant, ** - p < 0.01, *** - p < 0.001. H. Proportion of loops with and without TFs. Numbers near bars indicate total loop counts. Asterisks denote Fisher’s exact test *p*-values: ns - not significant, ** - p < 0.01, *** - p < 0.001.

Next, we partitioned chromatin loops into four categories (Q1–Q4) based on their contributions to PC1 and PC2 in the PCA for postmortem and iPSC-derived neuron samples (Supplemental Fig. S25A), where Q4 loops had the strongest influence on sample separation. Since most chromatin loops are formed via loop extrusion mediated by cohesin and CTCF (Banigan and Mirny 2020; Fudenberg et al. 2016), we annotated loop anchors based on the presence of CTCF peaks. Loop intensity and length were similar across the loop groups (Supplemental Fig. S25B,C), suggesting that fundamental properties of CTCF-mediated loops remain stable among cell types.

In our recent work, we identified long-range Polycomb-mediated interactions (“PcG-contacts”) as a neuronal hallmark (Pletenev et al. 2024). We thus asked if Polycomb could also influence the distinct short-range loop patterns observed between iPSC-derived and postmortem neurons. Indeed, we detected a subset of loops enriched at PcG-contact anchors that exhibit significantly higher intensity in postmortem neurons and strongly contribute to the separation between cell types (Upper plot: “both” vs “one anchor” Welch’s *t*-test, *p* = 2.99×10^-210^, “both” vs “no PcG-contact” Welch’s *t*-test, *p* = 4.71×10^-206^, Lower plot: “both” vs “one anchor” Welch’s *t*-test, *p* = 3.70×10^-115^, “both” vs “no PcG-contact” Welch’s *t*-test, *p* = 1.97×10^-113^; Figure 4D). These results suggest that considering chromatin marks beyond CTCF, particularly those linked to Polycomb, can reveal additional architectural differences in neuronal chromatin organization.

We identified differential chromatin loops between postmortem and iPSC-derived neurons, including 1,488 loops stronger in postmortem neurons and 3,696 loops stronger in iPSC-derived neurons. Given the established association between Polycomb complexes and the H3K27me3 mark (Cao et al. 2002; Czermin et al. 2002), we intersected these loops with H3K27me3 peaks for each cell type. iPSC-derived neurons showed no pronounced correlation between loop intensity or length and H3K27me3 occupancy in anchors (Figure 4E,F, Supplemental Fig. S26A-C). Conversely, H3K27me3-marked loops in postmortem neurons show increased length and strength (“both” vs “one anchor” Welch’s *t*-test, *p* = 1.29×10^-204^, “both” vs “no H3K27me3” Welch’s *t*-test, *p* = 3.55×10^-206^; Supplemental Fig. S26B; “both” vs “one anchor” Welch’s *t*-test, *p* = 5.11×10^-213^, “both” vs “no H3K27me3” Welch’s *t*-test *p* = 3.52×10^-215^; Supplemental Fig. S26D). Building on our previous observation that PcG-contacts repress transcription factor (TF) genes (Pletenev et al. 2024), we examined gene expression of TF and non-TF genes in differential chromatin loops. Postmortem-specific loops with H3K27me3 in anchors were preferentially associated with downregulated TFs in postmortem neurons (Figure 4G,H, Supplemental Fig. S27), and these repressed TFs were significantly enriched for developmental GO terms (Supplemental Fig. S28A). By contrast, in iPSC-derived neurons, genes associated with differential loops were, on average, upregulated across all loop categories and enriched for developmental GO terms (Figure 4G,H, Supplemental Fig. S28B), suggesting that activating interactions broadly promote developmental gene expression in these cells.

These findings suggest an additional repression mechanism in postmortem neurons at shorter genomic distances, complementing long-range PcG-contacts, which appears particularly important for silencing developmental TF programs in mature neurons.

### Long-range Polycomb-mediated interactions are not prominent in iPSC-derived neurons

Next, we focused directly on the long-range PcG-contacts. Visual inspection of Hi-C maps reveals that long-range PcG-contacts are much less prominent in iPSC-derived neurons (Figure 1E, Supplemental Fig. S29A). PCA of PcG-contact intensities further supports this observation, showing a clear separation between iPSC-derived and postmortem (infant and adult) samples (Figure 5A). Moreover, iPSC-derived samples position along PC1, which primarily describes the median level of PcG-contact frequency, closer to fetal than to postmortem neurons and almost overlap with NPCs. Consistently, average Hi-C maps display gradually increasing enrichment of PcG-contacts relative to local background from iPSC-derived to fetal to postmortem neurons, without a prominent difference between NPCs and iPSC-derived neurons (Figure 5B). This attenuation of PcG-contacts in iPSC-derived neurons is observed genome-wide and across a wide range of genomic distances (Figure 5C). Overall, our observations suggest that at this level of chromatin organization, iPSC-derived neurons exhibit even more immature state than in loops, TADs, or compartment analyses. A weak but noticeable enrichment is also observed in iPSCs, which cluster closer to postmortem neurons along PC2, perhaps reflecting specific PcG interactions controlling genes that remain poised in iPSCs but are fully repressed in mature postmortem neurons.

**Figure 5.**
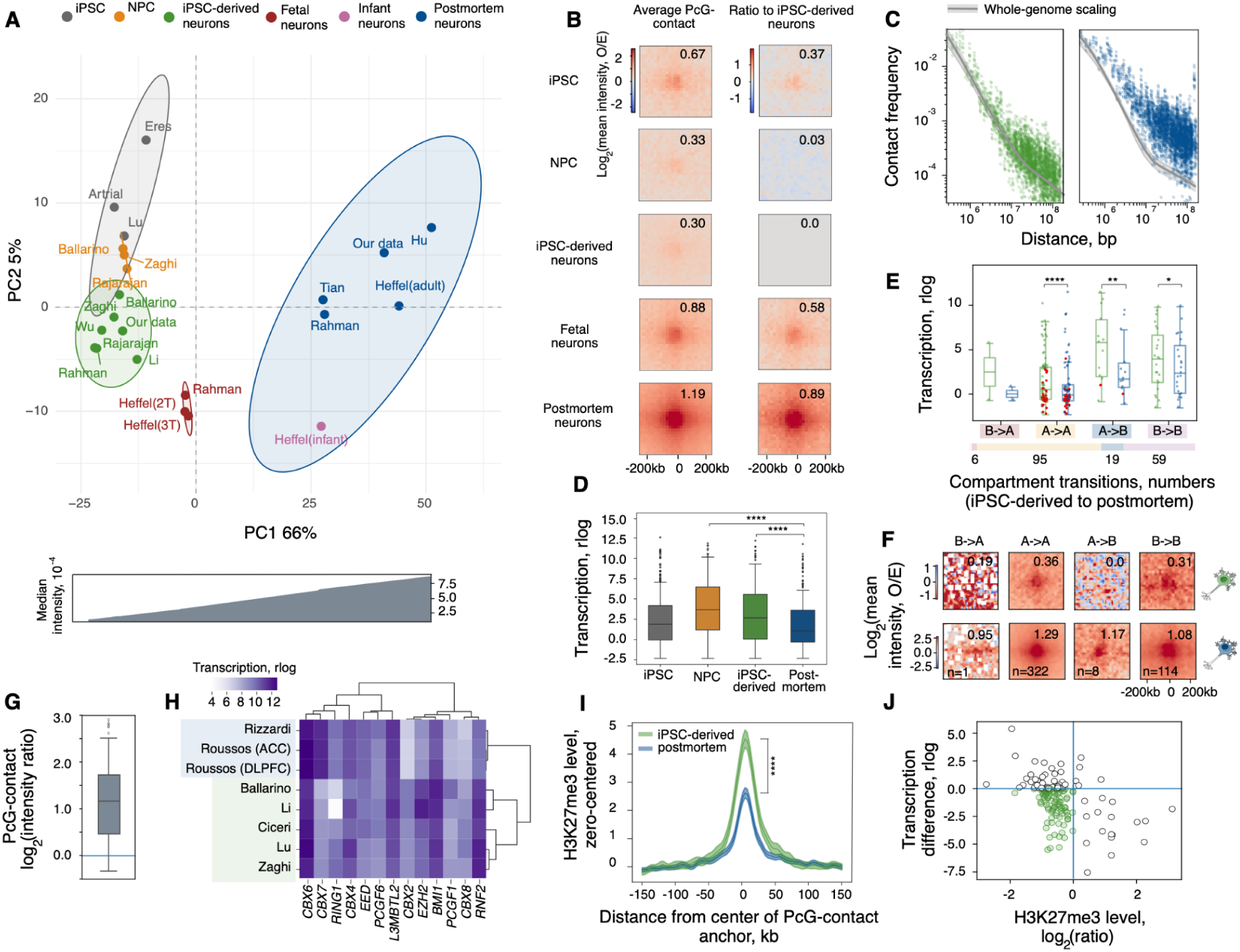
Analysis of long-range PcG-contacts. A. PCA of long-range PcG-contact intensities for analyzed groups (top) and median PcG-contact intensity across samples, ordered by PC1 values (bottom). B. Average PcG-contact intensities for each cell group. C. Contact scaling of PcG-contacts in iPSC-derived and postmortem (infant and adult) samples. Dots represent PcG-contact intensities across all samples. The gray line and shaded area represent the average genome-wide contact scaling and its standard deviation. D. Expression of protein-coding genes located at PcG-contact anchors. E. Expression of protein-coding genes located at PcG-contact anchors, classified by compartmental transitions. *HOX* genes are highlighted in red color. The stacked bar plot at the bottom shows the proportion of anchor regions in each compartment transition category (from iPSC-derived to postmortem neurons). F. Average PcG-contact intensities divided into groups by compartment transition categories in both anchors (from iPSC-derived to postmortem neurons), where numbers at the bottom indicate the PcG-contact counts. G. Intensity ratios of PcG-contacts (postmortem/iPSC-derived) excluding contacts with high gene expression (TPM < 1 at both anchors). H. Heatmap displaying rlog-normalized expression of PRC1/2 genes in iPSC-derived (highlighted in green) and postmortem neurons (highlighted in blue). I. Average normalized H3K27me3 ChIP-seq signal profiles centered on PcG-contact anchor midpoints. J. Expression differences compared to H3K27me3 differences (postmortem/iPSC-derived) in a union of iPSC-derived and postmortem ChIP-seq peaks at PcG-contact anchors. Green points mark loci with higher expression and higher H3K27me3 in iPSC-derived neurons. In all panels, asterisks indicate the significance level: *** - p < 0.001, **** - p < 0.0001.

To explore the relationship between PcG-contact prominence and gene expression, we analyzed RNA-seq datasets available for iPSCs, NPCs, iPSC-derived, and postmortem neurons. For postmortem neurons, we focused on brain regions with prominent PcG-contacts (dorsolateral prefrontal cortex and anterior cingulate cortex, Supplemental Fig. S29B). Genes located at anchors of PcG-contacts exhibit significantly lower expression than other genes (Wilcoxon test, *p* = 8.2×10^-27^, Supplemental Fig. S29C). However, their expression is significantly higher in NPCs and iPSC-derived neurons than in postmortem neurons (Wilcoxon test, *p* = 3.2×10^-20^ and *p* = 4.1×10^-12^, respectively; Figure 5D), consistent with the repressive role of Polycomb complexes. Moreover, PcG-contact anchors containing genes upregulated in iPSC-derived neurons, as opposed to anchors with non-differentially expressed genes, exhibit a significantly lower iPSC-derived/postmortem contact frequency ratio (Mann-Whitney *U* test, *p* = 2×10^-8^; Supplemental Fig. S29D). This finding supports a negative association between PcG-contact enrichment and transcription.

To dissect how PcG-contact differences intersect with changes in other chromatin features, we examined compartmental transitions at PcG-contact anchors. Although nearly 85% of anchors occupy the same compartment in iPSC-derived and postmortem neurons, up to 10% of anchors shift from the A compartment in iPSC-derived to the B compartment in postmortem neurons (Figure 5E, bottom). This A-to-B transition is accompanied by significant gene upregulation in iPSC-derived neurons (Wilcoxon test, *p* = 0.0002; Figure 5E, top, Supplemental Fig. S29E) and the greatest decrease in PcG-contact strength (Figure 5F). Genes at these PcG transitioning anchors function in early head and brain development: craniofacial skeleton formation (*IRX5, ALX4, SIX2*) (Bonnard et al. 2012; Kayserili et al. 2009; Garcez et al. 2013), inner ear development (*HOXA1*) (Makki and Capecchi 2010) and eye development (*PAX6*) (Baker et al. 2018). Notably, most anchors of PcG-contacts (55%) reside in A compartment in both iPSC-derived and postmortem neurons, harboring 92% of *HOX* genes, which are almost fully repressed in postmortem neurons but retain low expression in iPSC-derived neurons (Figure 5E, top).

Despite the observed correlation between expression and PcG-contact strength, this relationship does not fully account for the weakened contacts in iPSC-derived neurons. Specifically, analysis of PcG-contacts with low expression (TPM<1) at both anchors in both types of neurons reveals that these contacts are markedly less pronounced in iPSC-derived neurons than in postmortem neurons (Wilcoxon test, *p* = 1.2×10^-17^; Figure 5G), indicating that additional factors could influence PcG-contact formation, and these factors might be different between iPSC-derived and postmortem neurons. Consistently, iPSC-derived and postmortem neurons diverge in expression patterns of genes encoding Polycomb subunits, including *CBX7* and *CBX2* (Figure 5H). Transcription of *CBX7* gene, a PRC1 component and known negative regulator of axon growth and regeneration (Duan et al. 2018), weakly expressed in mature neurons, is decreased in iPSC-derived neurons, whereas *CBX2* and *CBX8* genes, whose orthologs are repressed by *Cbx7* in mouse ESCs (O’Loghlen et al. 2012), show higher expression in iPSC-derived neurons (Supplemental Table S2). In agreement with the observed tendency of iPSC-derived neurons to possess characteristics of immature neurons, these shifts in PRC1 composition at the level of transcripts mirror patterns of core PRC1 genes from the *Cbx* family seen during mouse cortical development (Duan et al. 2018; Gu et al. 2018). Studies in mouse ESCs suggest that Cbx proteins may affect chromatin condensate formation and phase separation (Guan et al. 2024; Niekamp et al. 2024), processes that may also contribute to the PcG-contact weakness in iPSC-derived neurons.

Given the direct involvement of H3K27me3 in PcG-mediated repression (Kraft et al. 2022), we further investigated the impact of this histone mark using ChIP-seq data for iPSC-derived and postmortem neurons. Both cell types show strong H3K27me3 enrichment at PcG-contact anchors; however, this enrichment is significantly higher in iPSC-derived neurons (Wilcoxon test, *p* = 3.4×10^-16^; Figure 5I; Supplemental Fig. S29F). This observation cannot be explained by a global increase in H3K27me3, as the mean fold-enrichment of ChIP-seq peaks within PcG-contact anchors, normalized to the mean enrichment across the rest of the genome, is significantly higher in iPSC-derived samples (Mann-Whitney U-test, *p* = 0.033; Supplemental Fig. S29G), emphasizing specificity of H3K27me3 to PcG-contact anchors. Considering the paradoxical observation of both high H3K27me3 (Figure 5I) and active transcription (Figure 5D) in iPSC-derived neurons, we performed a joint analysis: indeed, many anchor genes exhibited a simultaneous increase in both H3K27me3 and expression levels in iPSC-derived neurons (Figure 5J), highlighting a divergence from canonical repressive mechanisms in cultured neurons. Consistently, H3K27me3 has been shown to differentially target CBX7 and CBX2 (Zhen et al. 2016), underscoring that PcG complex composition rather than H3K27me3 abundance alone might control PcG-contact formation.

### Postmortem and iPSC-derived neurons reveal distinct FIREs landscape

We next analyzed frequently interacting regions (FIREs) – hotspots of local interactions, associated with regulation and cell-type specificity (Crowley et al. 2021). For each sample, we identified positions of FIREs, calculated FIREs scores (see Methods), and compiled the union of all FIREs positions across samples for further analysis. The projection of FIREs scores onto the space of principal components demonstrates separation of samples into two main groups corresponding to *in vitro*-derived cells and human brain tissue (Figure 6A, Supplemental Fig. S30A), without any noticeable effect of used restriction enzymes (Supplemental Fig. S30B). Consistent with other chromatin features, iPSC-derived neurons occupy an intermediate position between NPCs and fetal neurons. To further assess their similarity to other cell types, we quantified the relative enrichment of FIREs identified in iPSC-derived neurons across cell groups (Figure 6B). For each sample from iPSC, NPC, fetal and postmortem groups, we calculated the median signal intensity within FIREs defined in one of iPSC-derived samples and normalized it by the mean intensity of the other, not shared with iPSC-derived, FIREs. This approach enables estimation of the prominence of iPSC-derived FIREs in samples originating either from the same study or from independent datasets and accounts for global FIRE intensity. The strongest enrichment of iPSC-derived FIREs was observed in NPCs, highlighting the immature characteristics of iPSC-derived neurons (Mann–Whitney *U* test, p = 0.0007, 2.37×10⁻⁵, and 1.34×10⁻⁶ for comparisons with iPSCs, fetal, and postmortem neurons, respectively; Figure 6B).

**Figure 6.**
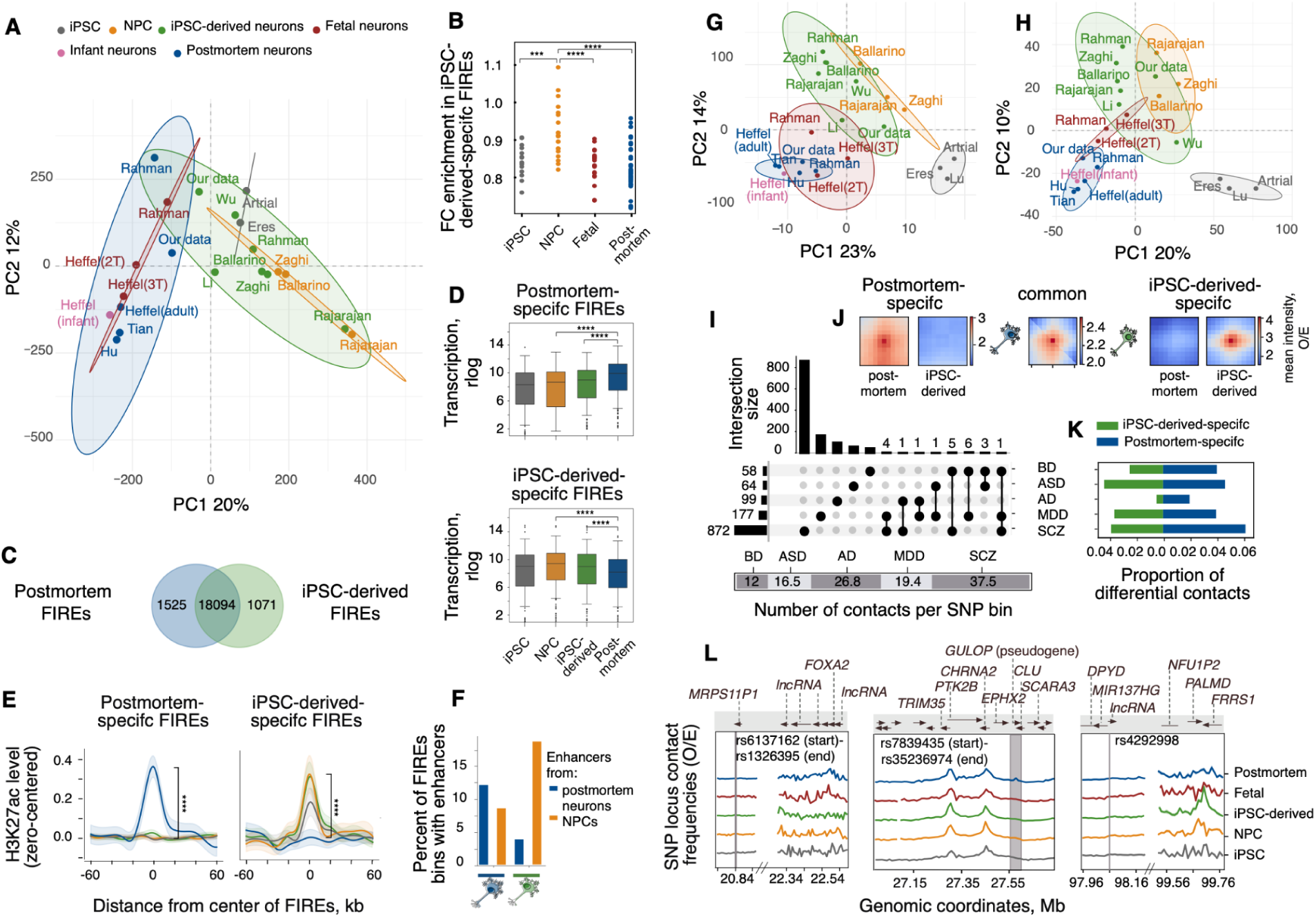
Analysis of FIREs and contacts involving disease-associated SNPs. A. PCA of FIREs scores for analyzed groups. B. Relative enrichment of FIREs from iPSC-derived neurons in other cell groups. C. Venn diagram showing overlap of FIREs, based on the differential analysis of FIREs scores. D. Expression levels of genes with promoters in postmortem-specific (top) and iPSC-derived-specific (bottom) FIREs in iPSCs, NPCs, iPSC-derived and postmortem (infant and adult) neurons. E. Average normalized H3K27ac ChIP-seq profile centered on postmortem-specific (left) and iPSC-derived-specific (right) FIREs. F. Percentage of differential FIREs bins containing enhancers defined in postmortem neurons and NPCs. Left two bars: fraction of postmortem-specific FIREs harboring postmortem vs NPCs enhancers. Right two bars: fraction of iPSC-derived-specific FIREs with postmortem vs NPCs enhancers. G-H. PCA of intensities of significant contacts involving SNPs associated with SCZ (G) or AD (H). I. UpSet plot showing numbers of shared and unique 20-kb genomic bins containing GWAS SNPs associated with BD, ASD, AD, MDD, or SCZ (top). Numbers of significant contacts per SNP-containing bin (bottom) J. Average contact intensities for common (center) and differential (left, right) disease-associated contacts, illustrated for SCZ-associated SNPs. K. Proportion of differential contacts among all significant contacts. Blue: higher intensity in postmortem neurons; green: higher intensity in iPSC-derived neurons. L. Example of average observed/expected contact frequency profiles for schizophrenia-associated loci showing cell-type-specific (left, right) or shared (middle) chromatin interactions between iPSC-derived and post-mortem neurons. In all panels, asterisks indicate the significance level: *** - p < 0.001, **** - p < 0.0001.

To further explore FIREs properties, we focused on iPSC-derived and postmortem (infant and adult) samples and conducted a differential analysis of their FIREs scores. While the majority of FIREs show similar patterns of *cis*-interactions in both groups, nearly 5% of FIREs are significantly upregulated in iPSC-derived neurons and ∼7% in postmortem neurons (*limma* differential analysis, BH–corrected *p* < 0.05, Figure 6C, Supplemental Fig. S30C). We next examined expression of genes associated with these differential FIREs for all cell types (except fetal, for which data were not available). Consistent with the transcription-activating role of FIREs, genes with promoters harbored by iPSC-derived-specific FIREs demonstrate significantly higher expression in iPSC-derived neurons (and similarly for postmortem-specific FIREs in postmortem neurons) (Figure 6D). Moreover, mirroring the PCA segregation of *in vitro* cells as a separate cluster, genes in iPSC-derived-specific FIREs are also highly expressed in NPCs and even in iPSCs (Figure 6D, bottom). GO enrichment analysis further highlights functional dissimilarity of genes within differential FIREs. In particular, iPSC-derived-specific FIREs are enriched for GO terms associated with stem cell population maintenance and other non-neuronal processes (Supplemental Fig. S30D). In contrast, postmortem-specific FIREs are enriched for neuronal GO categories such as postsynaptic specialization and apical dendrite (Supplemental Fig. S30E).

Because chromatin loops are also involved in regulation of gene expression, we explored the genomic co-localization of differential FIREs and differential loops FIREs. Nearly 23% of postmortem-specific FIREs overlap postmortem-specific loops and ∼15% of iPSC-derived-specific FIREs overlap iPSC-derived-specific loops (Supplemental Fig. S30F). This association between differential FIREs and differential loops is more pronounced in postmortem than in iPSC-derived neurons (Pearson’s chi-squared test, *p* = 4.6×10^-12^and 0.0038, respectively), suggesting that the weak coordination between loops and FIREs, which are both responsible for transcription regulation, may contribute to the immaturity of *in vitro* differentiated cells.

As FIREs are shown to intersect with regions exhibiting *cis*-regulatory epigenomic signatures (Crowley et al. 2021), we integrated genomic positions of differential FIREs with public data on active histone marks attributed to enhancers. Specifically, we gathered H3K27ac ChIP-seq data for all cell types (except fetal, for which data were not available) and H3K4me3 peaks (to mask active promoters and isolate enhancers). The masked H3K27ac profiles centered on differential FIREs reveal a strong significant enrichment matching the origin of cells (Wilcoxon test, *p* = 5.1×10^-50^ for postmortem-specific FIREs and *p* = 1.6×10^-52^ for iPSC-derived-specific FIREs; Figure 6E; Supplemental Fig. S30G).

Additionally, we leveraged the union of enhancer annotations from Dong et al. (2022) and GeneHancer (Fishilevich et al. 2017) for postmortem neurons and NPCs, respectively, and compared their genomic positions with positions of differential FIREs. Postmortem-specific FIREs demonstrate high association with enhancers defined in postmortem neurons (Figure 6F, Pearson’s chi-squared test, *p* = 6.8×10^-27^). A similar tendency is observed for iPSC-derived-specific FIREs and NPCs enhancers (Figure 6F, Pearson’s chi-squared test, *p* = 7.9×10^-27^), in agreement with the trends seen in the H3K27ac masked profiles (Figure 6E).

Collectively, these results reveal a prominent divergence between iPSC-derived and postmortem neurons in fine-grained chromatin interactions like FIREs. In line with the prominent regulatory function of FIREs, our observations are supported at the levels of gene expression, epigenetic profiles, and functional annotation.

### Applicability of *in vitro* differentiated neuronal cells for GWAS annotation

Because Hi-C maps are increasingly used to link unattributed GWAS SNPs and genes, we evaluated the ability of different Hi-C datasets to capture contacts involving disease-associated SNPs. To illustrate this ability, we focused on credible SNPs associated with four psychiatric disorders – schizophrenia (SCZ) (Pardiñas et al. 2018), bipolar disorder (BD) (O’Connell et al. 2025), major depressive disorder (MDD) (Adams et al. 2025), and autism spectrum disorder (ASD) (Grove et al. 2019) – as they were shown to be primarily active in neurons (Liu et al. 2025; Hu et al. 2021).

To extend beyond disorders with a presumed neurodevelopmental origin, we also included Alzheimer’s disease (AD) (Wightman et al. 2021). In addition to the established role of microglia, AD pathogenesis can originate from and be exacerbated by processes in other cell types, including neurons (Saleem et al. 2026; Rosenthal et al. 2022; Henstridge et al. 2019). Specifically, mitochondrial dysfunction, aberrant calcium homeostasis, and altered cholesterol metabolism occurring in neurons have been linked to AD neurodegeneration (Dong et al. 2025; Ge et al. 2022; Wolozin 2004). Many AD-associated SNPs lie in genes not directly involved in amyloid-β or tau pathology but rather in genes involved in other cellular processes; for example, *CSTF1* (RNA processing and polyadenylation) and *EXOSC4* (RNA exosome component) (Yeganeh Markid et al. 2024; Fasken et al. 2024). Further evidence for neuronal involvement comes from high-risk genes with high neuronal expression, such as *BIN1*, *ABCA7*, and *FERMT2* (Saha et al. 2024; von Maydell et al. 2025; Eysert et al. 2021).

To ensure that the study of AD-associated SNPs is relevant to neurons, we examined expression levels of genes carrying SNPs and genes interacting with SNP loci in neurons. While microglial expression dominates for the first group of genes, they are also expressed at moderate levels in other cell types, including neurons (Supplemental Fig. S31A,B). For genes interacting with AD-associated SNPs via neuronal Hi-C maps, comparable expression is observed in microglia and neurons (Supplemental Fig. S31C,D). We also examined genomic positions of AD-associated SNPs relative to candidate cis-regulatory elements (cCREs) annotated in specific cell types (Liu et al. 2025). Although prior studies reported strong LDSC enrichment in neuron-specific open chromatin for psychiatric disorders and in glial/microglial open or acetylated chromatin for AD (Liu et al. 2025; Hu et al. 2021), direct intersection of SNP coordinates with cCREs reveals that the largest proportions of AD-associated SNPs map to excitatory neurons from layer 3–4 (Exc L3–4) and activated microglia, compared to other cell types (Supplemental Fig. S31E,F). Collectively, these results indicate that, while neurons may not represent the primary causal cell type in AD pathogenesis, they warrant consideration with respect to chromatin interactions involving AD-associated SNPs as a potential layer of the regulatory landscape.

To identify SNP-associated interactions, a straightforward approach was applied: for each sample, we detected significant contacts (see Methods) involving a disease-associated SNP within a 5-Mb window, then pooled these significant contacts across samples and aggregated their interaction frequencies within +- 1 bin around the contacts. PCA of the intensities of these SNP-associated contacts reveals that iPSC-derived samples are closer to both NPCs and fetal samples than to postmortem neurons, consistent with our observations for larger-scale chromatin features (Figure 6G,H, Supplemental Fig. S32A-C). This pattern, preserved across all the examined disorders, was not driven by overlap of SNPs across GWAS datasets, as shared 20-kb bins containing SNPs from different disorders were rare (Figure 6I, top). The number of significant contacts per tested bin also varied across diseases, with higher values for SCZ and AD, which may indicate greater regulatory complexity underlying these conditions (Figure 6I, bottom). In addition, compared with other diseases, SNP-associated contacts detected for AD showed greater dissimilarity between postmortem neurons and less mature cell types, as indicated by their increased distance from these cell types (Figure 6H).

Analysis of aggregated interaction intensities identified SNP-associated contacts specific to either iPSC-derived or postmortem neurons (Figure 6J). The proportion of differential contacts varied across disorders (Figure 6K). Notably, AD displayed a distinct pattern compared with psychiatric and neurodevelopmental disorders (SCZ, BD, MDD, ASD), showing the fewest uncommon contacts, most of which were stronger in postmortem neurons. The scarcity of iPSC-derived–specific contacts is consistent with the late onset of AD and minimal neurodevelopmental contribution to its pathogenesis.

Differential contacts may reflect stage-specific regulatory mechanisms across neuronal maturation. For example, among schizophrenia-associated SNP contacts specific to postmortem neurons, we identified an interaction with the transcription factor *FOXA2* (Figure 6L, left). In addition to its role in neuronal development, FOXA2 is essential for the maintenance of dopaminergic neurons in the adult brain (Domanskyi et al. 2014). Another contact – with higher intensity in iPSC-derived neurons – corresponds to an interaction with the transcription end site of *FRRS1* (Figure 6L, right). Although FRRS1 is critical for neuronal viability (Han et al. 2017), it is also implicated in the biogenesis of the AMPA receptor complex (Stewart et al. 2019). We also revisited two representative examples from a fetal brain Hi-C study (Won et al. 2016) involving schizophrenia-associated SNPs, plotting the profile of contact frequencies around GWAS loci interacting with the *CHRNA2/PTK2* gene region (Figure 6L, middle) and with *SOX2* (Supplemental Fig. S32D). Both profile plots reveal an increase in contact frequency between the GWAS locus and the gene in almost every group. Notably, contacts involving the *CHRNA2* gene, which is poorly expressed in iPSCs-derived neurons (Chatzidaki et al. 2015), show similar profiles across all cell types, indicating that mutually exclusive transcription activation or repression via SNP–gene contacts is not the only possible scenario by which disease risk variants can act.

Together, these findings highlight the value of integrating Hi-C datasets from iPSC-derived and postmortem neurons to study chromatin interactions at GWAS loci associated with psychiatric and neurodevelopmental disorders.

## DISCUSSION

iPSC-derived neurons are widely used to model mature neurons in studies of brain-related diseases, as they provide an accessible and renewable source of neural cells compared to limited postmortem tissue. However, direct comparisons between iPSC-derived and adult brain neurons remain sparse, despite being crucial for establishing their validity as a model system. Existing research mainly focuses on transcriptomic profiling, suggesting that iPSC-derived neurons often retain developmental characteristics typical of earlier stages of neuronal development (Handel et al. 2016; Hoffman et al. 2017). Whether this developmental immaturity extends to higher-order chromatin organization has remained unclear. To address this gap, we performed a systematic cross-comparison of precursor cells (ESCs, iPSCs, NPCs, NESs), iPSC-derived neurons, and fetal and adult postmortem neurons, at all levels of chromatin architecture — from widely studied features such as compartments, TADs, and loops to more specialized structures including FIREs and PcG-mediated contacts.

Our multi-scale analysis of 62 human and 34 mouse Hi-C maps indicates that iPSC-derived neurons mostly exhibit an immature chromatin state similar to earlier developmental stages. At the compartment level, they occupy an intermediate state between precursor cells and postmortem neurons, displaying stronger compartmentalization than both fetal and postmortem neurons. This pronounced compartmentalization aligns with previous findings reporting stronger compartment structures in non-neuronal cells compared to neurons, highlighting weak compartmentalization as a distinctive feature of neuronal cells (Pletenev et al. 2024). Immaturity of iPSC-derived neurons is further reflected in TAD organization. We detected additional borders inherited from precursor cells enriched with developmentally regulated genes. Structural properties of TADs in iPSC-derived neurons are in line with these findings. Consistent with previous observations in mouse neural differentiation (Bonev et al. 2017), we noted an increase in TAD size in adult neurons compared to other cell populations, suggesting that TAD enlargement may be a characteristic feature of neuronal maturation. At the level of chromatin loops, iPSC-derived neurons lack the long-range, high-intensity loops enriched for H3K27me3 that are found to suppress developmental genes in postmortem neurons.

Beyond conventional chromatin features, our analysis of PcG-contacts reveals another distinctive property of iPSC-derived neurons: despite higher levels of H3K27me3 at PcG-contact anchors, their interactions are substantially weaker than in postmortem neurons. The concurrent increase in both H3K27me3 and expression levels at certain loci – seemingly paradoxical given the repressive role of this mark – has also been observed during *in vitro* hepatocyte differentiation, suggesting that H3K27me3 may be partially uncoupled from transcriptional repression in cultured cells (Vanhove et al. 2016). Together with observations at the loop level, these findings suggest an incomplete establishment of mature repressive mechanisms through H3K27me3-mediated contacts in the *in vitro* environment, potentially contributing to the immature state of these neurons.

Additionally, in the context of chromatin features associated with enhancer activity, iPSC-specific FIREs show enrichment patterns more similar to neural stem progenitor cells than to mature neurons. Notably, we observed reduced prominence of FIREs associated with critical neuronal functions, including synapse organization. This reduced FIREs intensity in neuron-specific regulatory regions may provide an additional explanation for the limited electrophysiological maturity of iPSC-derived neurons reported in the transcriptomic analysis (Handel et al. 2016).

Our analysis of SNP-associated contacts highlights differences between neurodegenerative disorders, exemplified by Alzheimer’s disease, and psychiatric conditions. For many psychiatric disorders, disease-relevant aspects of 3D genome organization appear to arise early during neuronal development (Owen et al. 2011; Khodosevich and Sellgren 2023; Rawdin et al. 2014; Kloiber et al. 2020) and may not be fully preserved in mature neurons. This highlights the value of including *in vitro* differentiated neurons alongside postmortem ones, as they can provide complementary insights into disease-associated regulatory interactions. In contrast, postmortem neurons likely represent a more appropriate model for Alzheimer’s disease, consistent with its late-onset pathology.

In summary, our cross-species analysis shows that although iPSC-derived neurons acquire key aspects of neuronal identity and functional properties (Bardy et al. 2016; Page et al. 2022), they retain immature characteristics across multiple levels of chromatin architecture, indicating that current differentiation protocols do not fully recapitulate mature neuronal chromatin organization.

Nevertheless, several limitations should be considered when interpreting these findings. First, our dataset compiles iPSC-derived neuronal samples differentiated under diverse protocols and toward varied neuronal subtypes, which may introduce heterogeneity that is not fully captured in group-level comparisons. Although we did not observe clear stratification by protocol duration, neuronal subtype, or other properties, incomplete annotation of some public datasets may limit the detection of subtype-specific chromatin features. Second, the approach applied to GWAS SNP annotation is exploratory. Large bin sizes and strategy of nearby bin aggregation limits resolution of this analysis, as bins can harbor several genes and independent SNPs. In addition, restricting the analysis to interactions beyond 40 kb may exclude relevant contacts. Together, these limitations may obscure fine-scale differences between iPSC-derived and postmortem neuronal maps.

Overall, our findings highlight the need to carefully balance the limitations and translational advantages of iPSC-derived neuronal cultures when designing experimental studies, and underscore the importance of systematic *in vivo* model evaluation. Building on previous model benchmarking efforts that emphasized the importance of multi-omics integration (Childs et al. 2022), our work establishes chromatin architecture as an additional crucial parameter for model validation. By providing the largest uniformly processed collection of neuronal Hi-C maps to date, our study offers a standardized reference for future analyses of neuronal chromatin architecture. More broadly, the same comparative framework could be applied to other iPSC-derived lineages, including cardiomyocytes, to assess their similarity to primary tissues and their suitability as disease models. Ultimately, understanding and potentially bridging the chromatin architecture gap between cultured neurons and the adult brain could enhance the fidelity of disease models and advance our understanding of brain genomics.

## METHODS

### Hi-C open source data collection

Open source Hi-C datasets were collected using the NCBI GEO DataSets portal(Edgar 2002) with the following search expressions: “(Neuron) AND “Mus musculus” [porgn: txid10090]” for mouse data and “(Neuron) AND Homo sapiens”[porgn: txid9606]” for human data. Additionally, the “Other” study type filter was added to limit irrelevant queries. All the datasets were annotated manually (Supplemental Table S1).

### Cultivation and differentiation of iPSCs into cortical neurons

iPSC line IPSRG4S was maintained in mTeSR1 (Stemcell Technologies) or GibriS-8 (PanEco) media according to the manufacturer’s instructions. Generation of the IPSRG4S cell line was previously described in Nekrasov et al. (2016) and Holmqvist et al. (2016). iPSCs were differentiated into cortical neurons using a stepwise protocol with sequential culture in neural differentiation medium, neuronal progenitor medium, and NB and NBA maturation media. Detailed media compositions and differentiation procedures are provided in the Supplemental Methods.

### Processing of postmortem human brain tissue

Postmortem brain samples were obtained from the National BioService Russian Biospecimen CRO (St. Petersburg, Russia) with written informed consent and verified as neurologically healthy. Frontal blocks (1–1.5 cm) were stored at −80°C. The posterior superior temporal gyrus of the left hemisphere (BA22p, Wernicke’s area) was dissected using thse Atlas of the Human Brain as a guide. Blocks at AHB level 60 were equilibrated in a cryostat at −20°C, and 600–800 mg tissue was sectioned into 200–300 mg portions using sterile, pre-cooled tools and immediately frozen on dry ice. Detailed procedures are provided in the Supplemental Methods.

### Nuclei isolation from postmortem human brain tissue

Cell nuclei from adult donor brain tissue were isolated and sorted as previously described (Nott et al. 2021; Pletenev et al. 2024), with modifications (see Supplemental Methods). The collected nuclei were either snap-frozen for storage at −80 °C, or used immediately for Hi-C library preparation.

### Hi-C library preparation

Hi-C experiments were conducted on cultured iPSC-derived neurons (whole cells) and adult neuronal and non-neuronal FANS-sorted nuclei. Two biological replicates were performed in each case. Approximately 1 million cells or nuclei per sample were used. We followed the previously described protocol (Pletenev et al. 2024; Lieberman-Aiden et al. 2009) for the subsequent steps. Hi-C libraries were sequenced on Illumina NovaSeq 6000 by Skoltech Genomics Core Facility and Evrogen Joint Stock Company.

### Hi-C data processing

Raw reads of each Hi-C library were mapped to the human *hg38* reference genome with the distiller-nf pipeline, v.0.3.3 (Open2C et al. 2024), filtered, and converted into binned Hi-C interaction matrices. Most downstream analyses were performed on Hi-C matrices merged by cell type at 5 kb resolution. Values along the main diagonal were removed to minimize potential ligation artifacts, and the resulting matrices were ICE normalized (Imakaev et al. 2012). Sample metadata is detailed in the Supplemental materials (Supplemental Table S3).

Full details of library preparation and computational processing are provided in the Supplemental Methods.

### Snm3C-seq data processing

Publicly available snm3C-seq datasets were used to analyze 3D chromatin organization across neural cell types and developmental stages (Supplemental Table S4). Contact pairs were obtained from GSE215353 and GSE213950 and grouped according to cell-type annotations from the original studies. For the developing brain data (Heffel et al. 2024), cortical neurons were divided into four stages (2T, 3T, infant, adult), and for adult brain data (Tian et al. 2023), cells were grouped by neuronal class and brain region; groups with insufficient cell numbers were excluded.

For each group, contact pairs were subsampled to match bulk Hi-C depth and aggregated into pseudobulk contact maps using cooler library, v.0.10.2. Matrices were processed analogously to Hi-C data, including removal of diagonal values and ICE normalization. Adult brain snm3C-seq data were merged across regions and cell types for downstream analyses (see Supplemental Methods for details).

### Downstream analysis inclusion criteria

To minimize batch effects and ensure comparability across the large number of collected Hi-C datasets, all samples underwent a multistep selection procedure. The inclusion criteria included the absence of experimental treatments or genetic modifications, consistent cell sorting conditions, adequate sequencing quality, and a sufficient number of unique contact pairs.

In total, 211 human and 83 mouse Hi-C maps passed this filtering procedure. These maps were subsequently merged by cell type and brain region, yielding 62 human and 34 mouse Hi-C maps for downstream analyses. Full details of the sample selection procedure are provided in the Supplemental Methods. Number of merged samples and datasets used for the downstream analysis provided in Supplemental Table S5.

### Contact probability analysis

Distance-dependent contact probability was computed from 5-kb binned Hi-C maps using the “expected_cis” function from the cooltools library, v.0.5.1, averaging intra-chromosomal contacts across autosomes. Samples were grouped by cell type or developmental stage, and mean contact probability at each genomic separation was calculated to generate representative scaling curves, visualized as log-log plots (normalized at 10 kb where indicated). Chromosomes X, Y, and mitochondria were excluded.

Scaling slopes were derived by differentiating log-transformed contact frequencies with respect to log genomic distance using numpy “np.gradient” function, and local minima and maxima were identified with the “find_peaks” function from the scipy.signal module of SciPy v. 1.11.4 (Virtanen et al. 2020). Comparisons of slope extrema between groups were assessed using Mann–Whitney *U* tests (see Supplemental Methods for full workflow).

### Identification and analysis of compartments

To identify compartments from Hi-C data, we first used Inspectro, v.0.2.0 (Spracklin et al. 2023) to cluster genomic bins based on eigenvector decomposition. For each Hi-C map, clustering was performed with the following parameters: “binsize: 50000”, “n_eigs: 5”, “n_clusters: [16]”, and “decomp_mode:’trans’”. We then selected the first 5 principal components that correlated with genomic GC content using the eigs_cis() function from the cooltools package with parameters: “n_eigs=5”, “sort_metric=’spearmanr’”. From the compartment tracks obtained by the eigenvector decomposition, we retained the one showing the highest Pearson’s correlation with the cooltools principal components. Chromosome Y, M, and the short arms of acrocentric chromosomes (Chr13p, Chr14p, Chr15p, Chr21p, and Chr22p) were excluded from analysis. Code for saddle plot generation was adapted from cooltools tutorial(cooltools). To calculate compartment strength, we used the mean intensity of the top 20% of corresponding interactions from the saddle matrix.

### Identification and analysis of TAD borders

TAD borders were identified from Hi-C maps using the “insulation” module from the cooltools package, v.0.6.1 (Open2C et al. 2022), with insulation scores computed at 15-kb resolution using a 150-kb window and a minimum valid pixel fraction of 0.75. To facilitate comparison across samples and account for minor positional fluctuations, TAD borders were clustered using the DBSCAN algorithm from the scikit-learn package, v.1.5.2 (Pedregosa et al. 2018), grouping borders within 30 kb into the same cluster. Clustered borders were treated as shared entities in downstream analyses. Sample-specific borders were identified by comparing border strength values between groups using the Mann–Whitney *U* test. Gene Ontology enrichment for genes located at TAD borders was performed using the “enrichGO” function from the clusterProfiler package (Yu et al. 2012). Detailed parameters are provided in the Supplemental Methods.

### TAD strength analysis

TAD strength (domain score) was calculated following Flyamer et al. (2020) by generating rescaled pileups using the coolpuppy package, v.1.0.0 (Flyamer et al. 2020) (see Supplemental Methods). Only TADs >10 bins from low-quality regions and < 1.5 Mb were included. Shared TADs were defined as those with borders ≤ 1.5 bins apart, and pairwise Jaccard similarity and Pearson’s correlation of strength scores were computed across samples. Border and corner dot enrichment were quantified as log_2_ ratios of contact intensities relative to the TAD interior. TADs with log₂FC > 0 were considered enriched. Detailed procedures, including pixel coordinates and enrichment calculations, are provided in the Supplemental Methods.

### Identification of chromatin loops

Chromatin loops were identified using the cooler and cooltools packages using *hg38* assembly of the human genome. Expected contact frequencies were computed with the “expected_cis” function from cooltools and used as a background for loop detection detecting significant loop interactions. To improve sensitivity across genomic scales, four custom kernels of different sizes were designed and applied as masks to detect contact enrichment.

Loop calling was performed using the “dots” function from cooltools with a maximum genomic distance of 12 Mb, FDR threshold of 0.16, tolerance of up to 5 missing bins, and clustering radius of 2.5 bin sizes. Loop sets obtained with different kernels were compared and filtered using custom Python scripts and pybedtools package (Dale et al. 2011) to ensure robustness and uniqueness of detected loops (see Supplemental Methods for details).

### Comparisons of chromatin loops between samples

To enable robust comparisons between samples, chromatin loops detected in individual datasets were grouped using the DBSCAN function from the scikit-learn package. Loops located within 2.5 bins (37.5 kb) at 15-kb resolution were assigned to the same group and considered equivalent.

For analyses requiring a complete matrix of loop intensities across all samples, such as PCA and correlation analyses, each cluster was assigned an intensity value in every sample. If a loop was not detected, intensity was estimated as the maximum median signal within a 3×3 bin window centered at the cluster coordinates. Clusters were further filtered using technical and intensity-based criteria, and loop anchors were intersected with genomic features using pybedtools.

Differential loop analysis between postmortem and iPSC-derived neurons was performed by comparing median loop intensities per cluster using the Mann–Whitney *U* test (see Supplemental Methods).

### Transcriptomics data processing and differential gene expression analysis

Publicly available RNA-seq datasets for postmortem adult and fetal brain samples, as well as iPSCs, NPCs, and iPSC-derived neurons, were used (Supplemental Table S6). When available, transcriptomic data from the same studies as the Hi-C datasets were used. For other cell types, the most closely matched datasets were selected from publicly available resources. Reads were trimmed, mapped, and quantified using standard pipelines (see Supplemental Methods for details). Only protein-coding genes were included in downstream analyses.

For downstream analyses, when gene expression data included biological replicates within a given cell group, rlog values were averaged across replicates to generate a single representative expression profile per group. Differential gene expression analysis was performed with DESeq2 (Love et al. 2014), v.1.36.0, with dataset, cell type, and donor included as variables in the design formula. Genes with an adjusted *p*-value ≤ 0.01 and |log2FoldChange| ≥ 2 were considered differentially expressed. Gene Ontology enrichment was performed using the clusterProfiler package, v.4.4.4 (Yu et al. 2012; Wu et al. 2021a), with Benjamini–Hochberg correction (see Supplemental Methods for full workflow).

### Principal component analysis (PCA) of Hi-C and gene expression data

Genome-wide chromatin insulation scores from Hi-C data and normalized RNA-seq expression matrices were used to compare cell types in mouse and human samples. PCA was performed using the PCA module from scikit-learn v.1.5.2, and the first two components were retained for visualization. PCA coordinates were plotted with ggplot2 v.3.5.1, with points colored by sample group; 95% confidence ellipses, derived from the PCA coordinates, were calculated using the “stat_ellipse” function. Arrows indicating differentiation trajectories were added manually. Detailed parameters and procedures are provided in the Supplemental Methods. To quantify multivariate separation between cell type groups, Mahalanobis distances were calculated between group centroids in 4-dimensional PC space (PC1-PC4), accounting for covariance structure. Statistical significance was assessed via permutation testing (10,000 iterations).

### Analysis of long-range PcG-contacts

The coordinates of long-range PcG-contacts were obtained from Pletenev et al. (2024). Contact frequencies, extracted from 100-kb resolution Hi-C matrices, were used for PCA and scaling plots, and mean heatmaps were generated from 20-kb scaling-normalized maps. Genes were grouped by differential expression between iPSC-derived and postmortem neurons (upregulated, downregulated, or not significant), and PcG-contacts were stratified by anchor expression category. Compartment transitions at PcG-contact anchors were assessed using PC1 values.

Polycomb group genes were clustered based on rlog-normalized expression using hierarchical clustering using correlation distance and average linkage. H3K27me3 enrichment at PcG-contact anchors was quantified by comparing ChIP-seq peak scores within and outside anchors, and associations with gene expression were evaluated. Quality control metrics (FRiP, RSC, NSC) were assessed to confirm robustness of the enrichment analysis (see Supplemental Methods for full workflow).

### FIREs and SNP analysis

Positions and scores of FIREs, defined as the total number of cis-interactions within the nearest 200 kb for each 10-kb bin passing default filters, were identified using the FIREcaller package (Crowley et al. 2021). FIREs were detected independently for each sample, and a union set of positions across all samples was compiled for downstream analyses. PCA of FIRE scores was used to examine dataset relationships, followed by a pairwise analysis to distinguish batch effects from genuine biological similarity.

Differential FIRE analysis between iPSC-derived and postmortem neurons was performed using the limma package (Ritchie et al. 2015) with covariates to account for potential technical bias and Benjamini–Hochberg correction. Genes were assigned to FIREs based on TSS positions, Gene Ontology enrichment was performed, and enhancers were linked to FIREs based on coordinate overlap.

SNP-associated chromatin interactions were identified at 20-kb resolution, aggregated for PCA and differential analysis, and visualized using heatmaps and contact profiles. Full methodological details are provided in the Supplemental Methods.

### ChIP-seq analysis

ChIP–seq datasets for H3K27me3 and H3K27ac were obtained from multiple sources (Supplemental Table S7). Broad peaks calling and generation of signal tracks were performed using the nf-core/chipseq pipeline (Patel et al. 2024) with the *hg38* genome assembly. For H3K27me3, average signal was computed in 5-kb genomic bins, whereas H3K27ac signal was calculated in 10-kb bins to match the resolution used for FIRE analysis. To exclude promoter-associated signals, we masked bins in the H3K27ac track that overlapped with H3K4me3 narrow peaks.

Average signal profiles around PcG-contact anchors were generated from the highest-quality datasets: Wu et al. (2021b) and Ciceri et al. (2024) for iPSC-derived neurons and Kozlenkov et al. (2018) for postmortem neurons. To highlight local enrichment, zero-centered average profiles shown in the main figures were adjusted by subtracting the signal baseline (see Supplemental Methods for full processing details).

## DATA ACCESS

All Hi-C data generated in this study have been submitted to the NCBI Gene Expression Omnibus (GEO; https://www.ncbi.nlm.nih.gov/geo/) under accession number GSE291967.

## CODE AVAILABILITY

Custom code for Hi-C maps processing and data analyses is available at https://github.com/DianaZagirova/cultural_vs_adult_neurons and is also provided as a Supplemental Code file.

## ACKNOWLEDGEMENTS

We thank the Center for Precision Genome Editing and Genetic Technologies for Biomedicine, IGB RAS for the cell sorting, Ilya Pletenev for the help with ChIP-seq and Snm3C-seq data processing, Genomics Core Facility and Dr. Elena Shagimardanova for sequencing and quality control of Hi-C libraries, and National BioService Russian Biospecimen CRO (St. Petersburg, Russia) for providing brain samples. H3K27me3 ChIP-seq data for this publication were obtained from NIMH Repository & Genomics Resource, a centralized national biorepository for genetic studies of psychiatric disorders.

## AUTHOR CONTRIBUTIONS

D.Z. processed Hi-C maps, analyzed global chromatin changes, TAD borders and chromatin loops, and contributed to manuscript writing. A.K. conceived the main idea of the work, analyzed FIRES and SNP data, investigated long-range Polycomb loops, processed ChIP-seq data and contributed to manuscript writing. K.M. collected and processed Hi-C data, created a database, analyzed compartments, conducted transcriptomic analysis, and contributed to manuscript writing. M.M. obtained Hi-C data from in-house iPSC-derived neurons and postmortem neurons and contributed to manuscript writing. N.V. analyzed TAD borders and contributed to manuscript writing. A.D. analyzed compartments. P.D. conducted transcriptomic analysis. O.E. performed Hi-C sample dissection from brain tissue. A.T. carried out neuronal nuclei sorting. I.P. processed ChIP-seq data and contributed to discussion. K.U. obtained Hi-C data from postmortem neurons. M.L. differentiated iPSCs into neurons and contributed to conceptualization. P.K. and S.R. contributed to conceptualization. S.U. contributed to conceptualization, manuscript editing, and writing. E.K. led the study, designed analyses, contributed to conceptualization, manuscript editing and writing, and provided resources. All authors contributed to the general discussion.

## REFERENCES

Baker LR, Weasner BM, Nagel A, Neuman SD, Bashirullah A, Kumar JP. 2018. Eyeless/Pax6 initiates eye formation non-autonomously from the peripodial epithelium. Development dev.163329.

Ballarino R, Bouwman BAM, Agostini F, Harbers L, Diekmann C, Wernersson E, Bienko M, Crosetto N. 2022. An atlas of endogenous DNA double-strand breaks arising during human neural cell fate determination. Sci Data 9: 400.

Banigan EJ, Mirny LA. 2020. Loop extrusion: theory meets single-molecule experiments. Curr Opin Cell Biol 64: 124–138.

Bardy C, Van Den Hurk M, Kakaradov B, Erwin JA, Jaeger BN, Hernandez RV, Eames T, Paucar AA, Gorris M, Marchand C, et al. 2016. Predicting the functional states of human iPSC-derived neurons with single-cell RNA-seq and electrophysiology. Mol Psychiatry 21: 1573–1588.

Bauer M, Vidal E, Zorita E, Üresin N, Pinter SF, Filion GJ, Payer B. 2021. Chromosome compartments on the inactive X guide TAD formation independently of transcription during X-reactivation. Nat Commun 12: 3499.

Bonev B, Mendelson Cohen N, Szabo Q, Fritsch L, Papadopoulos GL, Lubling Y, Xu X, Lv X, Hugnot J-P, Tanay A, et al. 2017. Multiscale 3D Genome Rewiring during Mouse Neural Development. Cell 171: 557–572.e24.

Bonnard C, Strobl AC, Shboul M, Lee H, Merriman B, Nelson SF, Ababneh OH, Uz E, Güran T, Kayserili H, et al. 2012. Mutations in IRX5 impair craniofacial development and germ cell migration via SDF1. Nat Genet 44: 709–713.

Burke EE, Chenoweth JG, Shin JH, Collado-Torres L, Kim S-K, Micali N, Wang Y, Colantuoni C, Straub RE, Hoeppner DJ, et al. 2020. Dissecting transcriptomic signatures of neuronal differentiation and maturation using iPSCs. Nat Commun 11: 462.

Cao R, Wang L, Wang H, Xia L, Erdjument-Bromage H, Tempst P, Jones RS, Zhang Y. 2002. Role of Histone H3 Lysine 27 Methylation in Polycomb-Group Silencing. Science 298: 1039–1043.

Chandrasekaran S, Espeso-Gil S, Loh Y-HE, Javidfar B, Kassim B, Zhu Y, Zhang Y, Dong Y, Bicks LK, Li H, et al. 2021. Neuron-specific chromosomal megadomain organization is adaptive to recent retrotransposon expansions. Nat Commun 12: 7243.

Chatzidaki A, Fouillet A, Li J, Dage J, Millar NS, Sher E, Ursu D. 2015. Pharmacological Characterisation of Nicotinic Acetylcholine Receptors Expressed in Human iPSC-Derived Neurons. PloS One 10: e0125116.

Childs CJ, Eiken MK, Spence JR. 2022. Approaches to benchmark and characterize *in vitro* human model systems. Development 149: dev200641.

Choi W-Y, Hwang J-H, Cho A-N, Lee AJ, Jung I, Cho S-W, Kim LK, Kim Y-J. 2020. NEUROD1 Intrinsically Initiates Differentiation of Induced Pluripotent Stem Cells into Neural Progenitor Cells. Mol Cells 43: 1011–1022.

Ciceri G, Baggiolini A, Cho HS, Kshirsagar M, Benito-Kwiecinski S, Walsh RM, Aromolaran KA, Gonzalez-Hernandez AJ, Munguba H, Koo SY, et al. 2024. An epigenetic barrier sets the timing of human neuronal maturation. Nature 626: 881–890.

cooltools. Compartments & Saddleplots - Cooltools Documentation. https://cooltools.readthedocs.io/en/latest/notebooks/compartments_and_saddles.html.

Cordazzo Vargas B, Shioda T. 2024. Association Between Activated Loci of HML-2 Primate-Specific Endogenous Retrovirus and Newly Formed Chromatin Contacts in Human Primordial Germ Cell-like Cells. Int J Mol Sci 25: 13639.

Crowley C, Yang Y, Qiu Y, Hu B, Abnousi A, Lipiński J, Plewczyński D, Wu D, Won H, Ren B, et al. 2021. FIREcaller: Detecting frequently interacting regions from Hi-C data. Comput Struct Biotechnol J 19: 355–362.

Czermin B, Melfi R, McCabe D, Seitz V, Imhof A, Pirrotta V. 2002. Drosophila Enhancer of Zeste/ESC Complexes Have a Histone H3 Methyltransferase Activity that Marks Chromosomal Polycomb Sites. Cell 111: 185–196.

Dale RK, Pedersen BS, Quinlan AR. 2011. Pybedtools: a flexible Python library for manipulating genomic datasets and annotations. Bioinformatics 27: 3423–3424.

Dixon JR, Selvaraj S, Yue F, Kim A, Li Y, Shen Y, Hu M, Liu JS, Ren B. 2012. Topological domains in mammalian genomes identified by analysis of chromatin interactions. Nature 485: 376–380.

Domanskyi A, Alter H, Vogt MA, Gass P, Vinnikov IA. 2014. Transcription factors Foxa1 and Foxa2 are required for adult dopamine neurons maintenance. Front Cell Neurosci 8. http://journal.frontiersin.org/article/10.3389/fncel.2014.00275/abstract (Accessed March 16, 2026).

Dong D, Ahmed W, Sagar R, Boyd RJ, Mahairaki V. 2025. Mapping key mitochondrial genes in Alzheimer’s disease through human tissue and iPSC derived neurons. Sci Rep 15: 42766.

Dong P, Hoffman GE, Apontes P, Bendl J, Rahman S, Fernando MB, Zeng B, Vicari JM, Zhang W, Girdhar K, et al. 2022. Population-level variation in enhancer expression identifies disease mechanisms in the human brain. Nat Genet 54: 1493–1503.

Duan R-S, Tang G-B, Du H-Z, Hu Y-W, Liu P-P, Xu Y-J, Zeng Y-Q, Zhang S-F, Wang R-Y, Teng Z-Q, et al. 2018. Polycomb protein family member CBX7 regulates intrinsic axon growth and regeneration. Cell Death Differ 25: 1598–1611.

Edgar R. 2002. Gene Expression Omnibus: NCBI gene expression and hybridization array data repository. Nucleic Acids Res 30: 207–210.

Eres IE, Luo K, Hsiao CJ, Blake LE, Gilad Y. 2019. Reorganization of 3D genome structure may contribute to gene regulatory evolution in primates ed. H.S. Malik. PLOS Genet 15: e1008278.

Eysert F, Coulon A, Boscher E, Vreulx A-C, Flaig A, Mendes T, Hughes S, Grenier-Boley B, Hanoulle X, Demiautte F, et al. 2021. Alzheimer’s genetic risk factor FERMT2 (Kindlin-2) controls axonal growth and synaptic plasticity in an APP-dependent manner. Mol Psychiatry 26: 5592–5607.

Fasken MB, Leung SW, Cureton LA, Al-Awadi M, Al-Kindy A, van Hoof A, Khoshnevis S, Ghalei H, Al-Maawali A, Corbett AH. 2024. A biallelic variant of the RNA exosome gene, EXOSC4, associated with neurodevelopmental defects impairs RNA exosome function and translation. J Biol Chem 300: 107571.

Feng J, Chuah YH, Liang Y, Cipta NO, Zeng Y, Warrier T, Elfar GARE, Yoon J, Grinchuk OV, Tay EXY, et al. 2024. PHF2 regulates genome topology and DNA replication in neural stem cells via cohesin. Nucleic Acids Res 52: 7063–7080.

Fishilevich S, Nudel R, Rappaport N, Hadar R, Plaschkes I, Iny Stein T, Rosen N, Kohn A, Twik M, Safran M, et al. 2017. GeneHancer: genome-wide integration of enhancers and target genes in GeneCards. Database 2017. https://academic.oup.com/database/article/doi/10.1093/database/bax028/3737828 (Accessed January 17, 2025).

Flyamer IM, Illingworth RS, Bickmore WA. 2020. Coolpup.py: versatile pile-up analysis of Hi-C data. Bioinformatics 36: 2980–2985.

Fraser J, Ferrai C, Chiariello AM, Schueler M, Rito T, Laudanno G, Barbieri M, Moore BL, Kraemer DC, Aitken S, et al. 2015. Hierarchical folding and reorganization of chromosomes are linked to transcriptional changes in cellular differentiation. Mol Syst Biol 11: 852.

Fudenberg G, Imakaev M, Lu C, Goloborodko A, Abdennur N, Mirny LA. 2016. Formation of Chromosomal Domains by Loop Extrusion. Cell Rep 15: 2038–2049.

Garcez RC, Le Douarin NM, Creuzet SE. 2013. Combinatorial activity of Six1-2-4 genes in cephalic neural crest cells controls craniofacial and brain development. Cell Mol Life Sci. http://link.springer.com/10.1007/s00018-013-1477-z (Accessed March 17, 2026).

Ge M, zhang J, Chen S, Huang Y, Chen W, He L, Zhang Y. 2022. Role of Calcium Homeostasis in Alzheimer’s Disease. Neuropsychiatr Dis Treat 18: 487–498.

Gevorgyan MM, Zhanaeva SYa, Alperina EL, Lipina TV, Idova GV. 2020. The composition of peripheral immunocompetent cell subpopulations and cytokine content in the brain structures of mutant Disc1-Q31L mice. Vavilov J Genet Breed 24: 770–776.

Grant ZL, Kuang S, Zhang S, Horrillo AJ, Rao KS, Kameswaran V, Joubran C, Lau PK, Dong K, Yang B, et al. 2025. Dose-dependent sensitivity of human 3D chromatin to a heart disease-linked transcription factor. http://biorxiv.org/lookup/doi/10.1101/2025.01.09.632202 (Accessed May 6, 2025).

Gu X, Wang X, Su D, Su X, Lin L, Li S, Wu Q, Liu S, Zhang P, Zhu X, et al. 2018. CBX2 Inhibits Neurite Development by Regulating Neuron-Specific Genes Expression. Front Mol Neurosci 11: 46.

Guan S, Tang J, Ma X, Miao R, Cheng B. 2024. CBX7C⋅PHC2 interaction facilitates PRC1 assembly and modulates its phase separation properties. iScience 27: 109548.

Han W, Wang H, Li J, Zhang S, Lu W. 2017. Ferric Chelate Reductase 1 Like Protein (FRRS1L) Associates with Dynein Vesicles and Regulates Glutamatergic Synaptic Transmission. Front Mol Neurosci 10: 402.

Handel AE, Chintawar S, Lalic T, Whiteley E, Vowles J, Giustacchini A, Argoud K, Sopp P, Nakanishi M, Bowden R, et al. 2016. Assessing similarity to primary tissue and cortical layer identity in induced pluripotent stem cell-derived cortical neurons through single-cell transcriptomics. Hum Mol Genet 25: 989–1000.

Heffel MG, Zhou J, Zhang Y, Lee D-S, Hou K, Pastor-Alonso O, Abuhanna KD, Galasso J, Kern C, Tai C-Y, et al. 2024. Temporally distinct 3D multi-omic dynamics in the developing human brain. Nature 635: 481–489.

Henstridge CM, Hyman BT, Spires-Jones TL. 2019. Beyond the neuron-cellular interactions early in Alzheimer disease pathogenesis. Nat Rev Neurosci 20: 94–108.

Hjelm BE, Salhia B, Kurdoglu A, Szelinger S, Reiman RA, Sue LI, Beach TG, Huentelman MJ, Craig DW. 2013. In vitro-differentiated neural cell cultures progress towards donor-identical brain tissue. Hum Mol Genet 22: 3534–3546.

Hoffman GE, Hartley BJ, Flaherty E, Ladran I, Gochman P, Ruderfer DM, Stahl EA, Rapoport J, Sklar P, Brennand KJ. 2017. Transcriptional signatures of schizophrenia in hiPSC-derived NPCs and neurons are concordant with post-mortem adult brains. Nat Commun 8: 2225.

Holmqvist S, Lehtonen Š, Chumarina M, Puttonen KA, Azevedo C, Lebedeva O, Ruponen M, Oksanen M, Djelloul M, Collin A, et al. 2016. Creation of a library of induced pluripotent stem cells from Parkinsonian patients. Npj Park Dis 2: 16009.

Hu B, Won H, Mah W, Park RB, Kassim B, Spiess K, Kozlenkov A, Crowley CA, Pochareddy S, Li Y, et al. 2021. Neuronal and glial 3D chromatin architecture informs the cellular etiology of brain disorders. Nat Commun 12: 3968.

Huang C, Su T, Xue Y, Cheng C, Lay FD, McKee RA, Li M, Vashisht A, Wohlschlegel J, Novitch BG, et al. 2017. Cbx3 maintains lineage specificity during neural differentiation. Genes Dev 31: 241–246.

Imakaev M, Fudenberg G, McCord RP, Naumova N, Goloborodko A, Lajoie BR, Dekker J, Mirny LA. 2012. Iterative correction of Hi-C data reveals hallmarks of chromosome organization. Nat Methods 9: 999–1003.

Jezierski A, Baumann E, Aylsworth A, Costain WJ, Corluka S, Banderali U, Sodja C, Ribecco-Lutkiewicz M, Alasmar S, Martina M, et al. 2022. Electrophysiological-and Neuropharmacological-Based Benchmarking of Human Induced Pluripotent Stem Cell-Derived and Primary Rodent Neurons. Stem Cell Rev Rep 18: 259–277.

Ji P, Yeh V, Ramirez T, Murata-Hori M, Lodish HF. 2010. Histone deacetylase 2 is required for chromatin condensation and subsequent enucleation of cultured mouse fetal erythroblasts. Haematologica 95: 2013–2021.

Jiang Y, Loh Y-HE, Rajarajan P, Hirayama T, Liao W, Kassim BS, Javidfar B, Hartley BJ, Kleofas L, Park RB, et al. 2017. The methyltransferase SETDB1 regulates a large neuron-specific topological chromatin domain. Nat Genet 49: 1239–1250.

Jiao J, Yang Y, Shi Y, Chen J, Gao R, Fan Y, Yao H, Liao W, Sun X-F, Gao S. 2013. Modeling Dravet syndrome using induced pluripotent stem cells (iPSCs) and directly converted neurons. Hum Mol Genet 22: 4241–4252.

Kayserili H, Uz E, Niessen C, Vargel I, Alanay Y, Tuncbilek G, Yigit G, Uyguner O, Candan S, Okur H, et al. 2009. ALX4 dysfunction disrupts craniofacial and epidermal development. Hum Mol Genet 18: 4357–4366.

Keough KC, Whalen S, Inoue F, Przytycki PF, Fair T, Deng C, Steyert M, Ryu H, Lindblad-Toh K, Karlsson E, et al. 2023. Three-dimensional genome rewiring in loci with human accelerated regions. Science 380: eabm1696.

Khodosevich K, Sellgren CM. 2023. Neurodevelopmental disorders—high-resolution rethinking of disease modeling. Mol Psychiatry 28: 34–43.

Kiefer L, Gaudin S, Rajkumar SM, Servito GIF, Langen J, Mui MH, Nawsheen S, Canzio D. 2024. Tuning cohesin trajectories enables differential readout of the Pcdhα cluster across neurons. Science 385: eadm9802.

Kloiber S, Rosenblat JD, Husain MI, Ortiz A, Berk M, Quevedo J, Vieta E, Maes M, Birmaher B, Soares JC, et al. 2020. Neurodevelopmental pathways in bipolar disorder. Neurosci Biobehav Rev 112: 213–226.

Koehler KR, Tropel P, Theile JW, Kondo T, Cummins TR, Viville S, Hashino E. 2011. Extended passaging increases the efficiency of neural differentiation from induced pluripotent stem cells. BMC Neurosci 12: 82.

Kozlenkov A, Li J, Apontes P, Hurd YL, Byne WM, Koonin EV, Wegner M, Mukamel EA, Dracheva S. 2018. A unique role for DNA (hydroxy)methylation in epigenetic regulation of human inhibitory neurons. Sci Adv 4: eaau6190.

Kraft K, Yost KE, Murphy SE, Magg A, Long Y, Corces MR, Granja JM, Wittler L, Mundlos S, Cech TR, et al. 2022. Polycomb-mediated genome architecture enables long-range spreading of H3K27 methylation. Proc Natl Acad Sci 119: e2201883119.

Kriukov D, Eremenko E, Smirnov D, Stein D, Tsitrina A, Golova A, Einav M, Khrameeva E, Toiber D. 2024. Nuclear expansion and chromatin structure remodeling in mouse aging neurons. NAR Mol Med 1: ugae011.

Li L, Chao J, Shi Y. 2018. Modeling neurological diseases using iPSC-derived neural cells: iPSC modeling of neurological diseases. Cell Tissue Res 371: 143–151.

Li P, Marshall L, Oh G, Jakubowski JL, Groot D, He Y, Wang T, Petronis A, Labrie V. 2019. Epigenetic dysregulation of enhancers in neurons is associated with Alzheimer’s disease pathology and cognitive symptoms. Nat Commun 10: 2246.

Lieberman-Aiden E, van Berkum NL, Williams L, Imakaev M, Ragoczy T, Telling A, Amit I, Lajoie BR, Sabo PJ, Dorschner MO, et al. 2009. Comprehensive mapping of long-range interactions reveals folding principles of the human genome. Science 326: 289–293.

Liu Z, Zhang S, James BT, Galani K, Mangan RJ, Fass SB, Liang C, Wagle MM, Boix CA, Tanigawa Y, et al. 2025. Single-cell multiregion epigenomic rewiring in Alzheimer’s disease progression and cognitive resilience. Cell 188: 4980–5002.e29.

Love MI, Huber W, Anders S. 2014. Moderated estimation of fold change and dispersion for RNA-seq data with DESeq2. Genome Biol 15: 550.

Lu L, Liu X, Huang W-K, Giusti-Rodríguez P, Cui J, Zhang S, Xu W, Wen Z, Ma S, Rosen JD, et al. 2020. Robust Hi-C maps of enhancer-promoter interactions reveal the function of non-coding genome in neural development and diseases. Mol Cell 79: 521–534.e15.

Makki N, Capecchi MR. 2010. Hoxa1 lineage tracing indicates a direct role for Hoxa1 in the development of the inner ear, the heart, and the third rhombomere. Dev Biol 341: 499–509.

Malachowski T, Chandradoss KR, Boya R, Zhou L, Cook AL, Su C, Pham K, Haws SA, Kim JH, Ryu H-S, et al. 2023. Spatially coordinated heterochromatinization of long synaptic genes in fragile X syndrome. Cell 186: 5840–5858.e36.

Mendez EF, Grimm SL, Stertz L, Gorski D, Movva SV, Najera K, Moriel K, Meyer TD, Fries GR, Coarfa C, et al. 2023. A human stem cell-derived neuronal model of morphine exposure reflects brain dysregulation in opioid use disorder: Transcriptomic and epigenetic characterization of postmortem-derived iPSC neurons. Front Psychiatry 14: 1070556.

Nekrasov ED, Vigont VA, Klyushnikov SA, Lebedeva OS, Vassina EM, Bogomazova AN, Chestkov IV, Semashko TA, Kiseleva E, Suldina LA, et al. 2016. Manifestation of Huntington’s disease pathology in human induced pluripotent stem cell-derived neurons. Mol Neurodegener 11: 27.

Niekamp S, Marr SK, Oei TA, Subramanian R, Kingston RE. 2024. Modularity of PRC1 composition and chromatin interaction define condensate properties. Mol Cell 84: 1651–1666.e12.

Nott A, Schlachetzki JCM, Fixsen BR, Glass CK. 2021. Nuclei isolation of multiple brain cell types for omics interrogation. Nat Protoc 16: 1629–1646.

Okhovat M, VanCampen J, Nevonen KA, Harshman L, Li W, Layman CE, Ward S, Herrera J, Wells J, Sheng RR, et al. 2023. TAD evolutionary and functional characterization reveals diversity in mammalian TAD boundary properties and function. Nat Commun 14: 8111.

O’Loghlen A, Muñoz-Cabello AM, Gaspar-Maia A, Wu H-A, Banito A, Kunowska N, Racek T, Pemberton HN, Beolchi P, Lavial F, et al. 2012. MicroRNA Regulation of Cbx7 Mediates a Switch of Polycomb Orthologs during ESC Differentiation. Cell Stem Cell 10: 33–46.

Open2C, Abdennur N, Abraham S, Fudenberg G, Flyamer IM, Galitsyna AA, Goloborodko A, Imakaev M, Oksuz BA, Venev SV. 2022. Cooltools: enabling high-resolution Hi-C analysis in Python. 2022.10.31.514564. https://www.biorxiv.org/content/10.1101/2022.10.31.514564v1 (Accessed February 24, 2026).

Open2C, Abdennur N, Fudenberg G, Flyamer IM, Galitsyna AA, Goloborodko A, Imakaev M, Venev SV. 2024. Pairtools: From sequencing data to chromosome contacts. PLoS Comput Biol 20: e1012164.

Owen MJ, O’Donovan MC, Thapar A, Craddock N. 2011. Neurodevelopmental hypothesis of schizophrenia. Br J Psychiatry 198: 173–175.

Page SC, Sripathy SR, Farinelli F, Ye Z, Wang Y, Hiler DJ, Pattie EA, Nguyen CV, Tippani M, Moses RL, et al. 2022. Electrophysiological measures from human iPSC-derived neurons are associated with schizophrenia clinical status and predict individual cognitive performance. Proc Natl Acad Sci 119: e2109395119.

Paşca AM, Sloan SA, Clarke LE, Tian Y, Makinson CD, Huber N, Kim CH, Park J-Y, O’Rourke NA, Nguyen KD, et al. 2015. Functional cortical neurons and astrocytes from human pluripotent stem cells in 3D culture. Nat Methods 12: 671–678.

Patel H, Espinosa-Carrasco J, Wang C, Ewels P, bot nf-core, Silva TC, Peltzer A, Langer B, Guinchard S, Garcia MU, et al. 2024. nf-core/chipseq: nf-core/chipseq v2.1.0 - Platinum Willow Sparrow. https://zenodo.org/records/13899404 (Accessed February 24, 2026).

Pedregosa F, Varoquaux G, Gramfort A, Michel V, Thirion B, Grisel O, Blondel M, Müller A, Nothman J, Louppe G, et al. 2018. Scikit-learn: Machine Learning in Python. http://arxiv.org/abs/1201.0490 (Accessed February 24, 2026).

Pletenev IA, Bazarevich M, Zagirova DR, Kononkova AD, Cherkasov AV, Efimova OI, Tiukacheva EA, Morozov KV, Ulianov KA, Komkov D, et al. 2024. Extensive long-range polycomb interactions and weak compartmentalization are hallmarks of human neuronal 3D genome. Nucleic Acids Res 52: 6234–6252.

Polovnikov KE, Brandão HB, Belan S, Slavov B, Imakaev M, Mirny LA. 2023. Crumpled Polymer with Loops Recapitulates Key Features of Chromosome Organization. Phys Rev X 13: 041029.

Prè D, Nestor MW, Sproul AA, Jacob S, Koppensteiner P, Chinchalongporn V, Zimmer M, Yamamoto A, Noggle SA, Arancio O. 2014. A time course analysis of the electrophysiological properties of neurons differentiated from human induced pluripotent stem cells (iPSCs). PloS One 9: e103418.

Rahman S, Dong P, Apontes P, Fernando MB, Kosoy R, Townsley KG, Girdhar K, Bendl J, Shao Z, Misir R, et al. 2023. Lineage specific 3D genome structure in the adult human brain and neurodevelopmental changes in the chromatin interactome. Nucleic Acids Res 51: 11142–11161.

Rajarajan P, Borrman T, Liao W, Schrode N, Flaherty E, Casiño C, Powell S, Yashaswini C, LaMarca EA, Kassim B, et al. 2018. Neuron-specific signatures in the chromosomal connectome associated with schizophrenia risk. Science 362: eaat4311.

Rawdin BJ, Lindqvist D, Bush N, Hamilton S, Boparai R, Mackin RS, Reus VI, Mellon SH, Wolkowitz OM. 2014. Neurodevelopmental and Neurobiological Aspects of Major Depressive Disorder. In Treatment of Neurodevelopmental Disorders (eds. R. Hagerman and R. Hendren), pp. 73–104, Oxford University Press https://academic.oup.com/book/35458/chapter/303479430 (Accessed March 17, 2026).

Ritchie ME, Phipson B, Wu D, Hu Y, Law CW, Shi W, Smyth GK. 2015. limma powers differential expression analyses for RNA-sequencing and microarray studies. Nucleic Acids Res 43: e47.

Rocks D, Shukla M, Ouldibbat L, Finnemann SC, Kalluchi A, Rowley MJ, Kundakovic M. 2022. Sex-specific multi-level 3D genome dynamics in the mouse brain. Nat Commun 13: 3438.

Rosenthal SB, Wang H, Shi D, Liu C, Abagyan R, McEvoy LK, Chen C-H. 2022. Mapping the gene network landscape of Alzheimer’s disease through integrating genomics and transcriptomics. PLoS Comput Biol 18: e1009903.

Rowley MJ, Corces VG. 2018. Organizational principles of 3D genome architecture. Nat Rev Genet 19: 789–800.

Saha O, Melo de Farias AR, Pelletier A, Siedlecki-Wullich D, Landeira BS, Gadaut J, Carrier A, Vreulx A-C, Guyot K, Shen Y, et al. 2024. The Alzheimer’s disease risk gene BIN1 regulates activity-dependent gene expression in human-induced glutamatergic neurons. Mol Psychiatry 29: 2634–2646.

Saleem K, Xiao Z, Zhu B, Ren Y, Yan Z, Feng J. 2026. Elevated SGK1 increases Tau phosphorylation and microtubule instability in Alzheimer’s patient-derived cortical neurons. Mol Psychiatry 31: 332–342.

Sandor C, Robertson P, Lang C, Heger A, Booth H, Vowles J, Witty L, Bowden R, Hu M, Cowley SA, et al. 2017. Transcriptomic profiling of purified patient-derived dopamine neurons identifies convergent perturbations and therapeutics for Parkinson’s disease. Hum Mol Genet 26: 552–566.

Spracklin G, Abdennur N, Imakaev M, Chowdhury N, Pradhan S, Mirny LA, Dekker J. 2023. Diverse silent chromatin states modulate genome compartmentalization and loop extrusion barriers. Nat Struct Mol Biol 30: 38–51.

Stein JL, de la Torre-Ubieta L, Tian Y, Parikshak NN, Hernández IA, Marchetto MC, Baker DK, Lu D, Hinman CR, Lowe JK, et al. 2014. A quantitative framework to evaluate modeling of cortical development by neural stem cells. Neuron 83: 69–86.

Stewart M, Lau P, Banks G, Bains RS, Castroflorio E, Oliver PL, Dixon CL, Kruer MC, Kullmann DM, Acevedo-Arozena A, et al. 2019. Loss of *Frrs1l* disrupts synaptic AMPA receptor function, and results in neurodevelopmental, motor, cognitive and electrographical abnormalities. Dis Model Mech dmm.036806.

Takahashi K, Tanabe K, Ohnuki M, Narita M, Ichisaka T, Tomoda K, Yamanaka S. 2007. Induction of Pluripotent Stem Cells from Adult Human Fibroblasts by Defined Factors. Cell 131: 861–872.

Tian W, Zhou J, Bartlett A, Zeng Q, Liu H, Castanon RG, Kenworthy M, Altshul J, Valadon C, Aldridge A, et al. 2023. Single-cell DNA methylation and 3D genome architecture in the human brain. Science 382: eadf5357.

Titus KR, Simandi Z, Chandrashekar H, Paquet D, Phillips-Cremins JE. 2023. Cell type-specific loops linked to RNA polymerase II elongation in human neural differentiation. http://biorxiv.org/lookup/doi/10.1101/2023.12.04.569731 (Accessed May 6, 2025).

Vanhove J, Pistoni M, Welters M, Eggermont K, Vanslembrouck V, Helsen N, Boon R, Najimi M, Sokal E, Collas P, et al. 2016. H3K27me3 Does Not Orchestrate the Expression of Lineage-Specific Markers in hESC-Derived Hepatocytes In Vitro. Stem Cell Rep 7: 192–206.

Virtanen P, Gommers R, Oliphant TE, Haberland M, Reddy T, Cournapeau D, Burovski E, Peterson P, Weckesser W, Bright J, et al. 2020. SciPy 1.0: fundamental algorithms for scientific computing in Python. Nat Methods 17: 261–272.

von Maydell D, Wright SE, Pao P-C, Staab C, King O, Spitaleri A, Bonner JM, Liu L, Yu CJ, Chiu C-C, et al. 2025. ABCA7 variants impact phosphatidylcholine and mitochondria in neurons. Nature 647: 462–471.

Wightman DP, Jansen IE, Savage JE, Shadrin AA, Bahrami S, Holland D, Rongve A, Børte S, Winsvold BS, Drange OK, et al. 2021. A genome-wide association study with 1,126,563 individuals identifies new risk loci for Alzheimer’s disease. Nat Genet 53: 1276–1282.

Wolozin B. 2004. Cholesterol and the Biology of Alzheimer’s Disease. Neuron 41: 7–10.

Won H, de la Torre-Ubieta L, Stein JL, Parikshak NN, Huang J, Opland CK, Gandal M, Sutton GJ, Hormozdiari F, Lu D, et al. 2016. Chromosome conformation elucidates regulatory relationships in developing human brain. Nature 538: 523–527.

Wu T, Hu E, Xu S, Chen M, Guo P, Dai Z, Feng T, Zhou L, Tang W, Zhan L, et al. 2021a. clusterProfiler 4.0: A universal enrichment tool for interpreting omics data. Innov Camb Mass 2: 100141.

Wu W, Kargbo-Hill SE, Nathan WJ, Paiano J, Callen E, Wang D, Shinoda K, Van Wietmarschen N, Colón-Mercado JM, Zong D, et al. 2021b. Neuronal enhancers are hotspots for DNA single-strand break repair. Nature 593: 440–444.

Yeganeh Markid T, Hosseinpour Feizi MA, Talebi M, Rezazadeh M, Khalaj-Kondori M. 2024. Gene expression investigation of four key regulators of polyadenylation and alternative adenylation in the periphery of late-onset Alzheimer’s disease patients. Gene 895: 148013.

Yu G, Wang L-G, Han Y, He Q-Y. 2012. clusterProfiler: an R package for comparing biological themes among gene clusters. Omics J Integr Biol 16: 284–287.

Zaghi M, Banfi F, Massimino L, Volpin M, Bellini E, Brusco S, Merelli I, Barone C, Bruni M, Bossini L, et al. 2023. Balanced SET levels favor the correct enhancer repertoire during cell fate acquisition. Nat Commun 14: 3212.

Zenk F, Zhan Y, Kos P, Löser E, Atinbayeva N, Schächtle M, Tiana G, Giorgetti L, Iovino N. 2021. HP1 drives de novo 3D genome reorganization in early Drosophila embryos. Nature 593: 289–293.

Zhen CY, Tatavosian R, Huynh TN, Duc HN, Das R, Kokotovic M, Grimm JB, Lavis LD, Lee J, Mejia FJ, et al. 2016. Live-cell single-molecule tracking reveals co-recognition of H3K27me3 and DNA targets polycomb Cbx7-PRC1 to chromatin. eLife 5: e17667.

