## Supplemental Materials for "Fetal signatures in the 3D genome of iPSC-derived neurons and their implications for disease modeling"

### Supplemental Methods

#### Cultivation and differentiation of iPSCs into cortical neurons

##### Compositions of cell culture media

Neural differentiation medium: DMEM/F12 (GIBCO), 1% serum replacement, 1% N2 supplement (GIBCO), 1× Glutamax, 50 units/ml penicillin-streptomycin, 10 μM SB431542, 2 μ M dorsomorphin and 0.5 μM LDN-193189. Neuronal progenitor medium: DMEM/F12, 2% B27 supplement (GIBCO), 1× Glutamax, 50 units/mL penicillin-streptomycin, 10 ng/mL, FGF2. NB maturation medium: Neurobasal (GIBCO), 2% B27 supplement, 1× Glutamax, 50 units/mL penicillin-streptomycin, 20 ng/mL BDNF, 20 ng/mL GDNF, 200 μM ascorbic acid. NBA maturation medium: Neurobasal-A (GIBCO), 2% B27 supplement, 1× Glutamax, 50 units/mL penicillin-streptomycin, 20 ng/mL BDNF, 20 ng/mL GDNF, 200 μM ascorbic acid and 10 μM forskolin.

##### iPSCs cultivation

iPSC line IPSRG4S was maintained in mTeSR1 (Stemcell Technologies) or GibriS-8 (PanEco) media according to the manufacturer's instructions. Generation of the IPSRG4S cell line was previously described in (Nekrasov et al. 2016; Holmqvist et al. 2016). Matrigel™ substrate (BD Biosciences) was used to coat the culture plates. Matrigel-coated culture dishes and plates were prepared according to the manufacturer's instructions. Cells were passaged using 0.05% trypsin (Gibco). ROCK inhibitor Y27632 (Stemgent) was added to the culture medium at a concentration of 5 μM for 1 day after seeding.

##### Differentiation of iPSCs into cortical neurons

iPSCs were detached with trypsin and plated at a density of 40,000 cells/cm$^2$ in mTeSR1 or GibriS-8 medium in the presence of 5 μM of ROCK inhibitor. At cell confluency, the mTeSR1 (GibriS-8) medium was replaced by the neural differentiation medium. The cells were cultured for 14 days, with the medium changed every other day. The resulting neural progenitors were detached with Versene solution (PanEco) by incubating the cells for 10 min in a $CO_2$ incubator at 37 °C. The cells were then centrifuged for 5 min at 250× g and washed with DMEM/F12 medium. The cells were plated at a density of 250,000–400,000 cells/cm$^2$ in cell culture-treated dishes coated with Matrigel and cultured in neuronal progenitor medium for 10 days. The medium was changed every other day. The resulting neuronal progenitors were detached with the Versene solution as described above. Then the cells were again plated at a density of 250,000–400,000 cells/cm$^2$ in cell culture-treated dishes coated with Matrigel and cultured for 7 days in NB maturation medium. The medium was changed every other day. After this stage, the cells could no longer be reseeded. The cells were cultured in NBA maturation medium for at least 2 weeks.

#### Processing of postmortem human brain tissue

This study was approved by the Bioethics Committee of the Institute of Gene Biology, Russian Academy of Sciences, and conducted in accordance with the Declaration of Helsinki. Postmortem human brain samples were obtained from the National BioService Russian Biospecimen CRO (St. Petersburg, Russia) with written informed consent from donors or their next of kin. All samples were verified by medical pathologists as neurologically healthy, with no history of neurological, psychiatric, or substance use disorders. Frontal brain blocks (1–1.5 cm thick) were stored at −80°C in foil-wrapped, zip-locked bags. The posterior superior temporal gyrus of the left hemisphere (BA22p, Wernicke's area) was identified using the Atlas of the Human Brain (AHB) to guide the dissection.

Blocks corresponding to AHB level 60 (MNI: −36.57; ICL: 34.82; MCP: 20.08) were placed in a Leica CM1950 cryostat at −20°C for 15–20 minutes before dissection. Using a metal scalpel, approximately 600–800 mg of frozen brain tissue was cut out and then sectioned into 200–300 mg portions. Samples were collected with sterile tweezers, transferred to tubes, and immediately placed on dry ice.

##### Nuclei isolation from postmortem human brain tissue

The tissue pieces were homogenized with an automated pestle homogenizer in 100 ul of fixative solution (1x PBS, 2% formaldehyde (Sigma-Aldrich)). The homogenate was rocked for 10 minutes at room temperature in a total volume of 10 ml of fixative solution, followed by the addition of 2 M glycine to a final concentration of 125 mM to quench the cross-linking reaction. The homogenate was then centrifuged at 1100g for 5 min at 4°C. The pellet was washed twice with NF1 buffer (10 mM Tris–HCl, pH 8.0; 1 mM EDTA; 5 mM $MgCl_2$; 100 mM sucrose; 0.5% Triton X-100) and incubated in NF1 buffer for 30 min. The mixture was homogenized on ice using a Dounce homogenizer and filtered through a 70 um cell strainer. A sucrose cushion (10 mM Tris-HCl, pH 8.0; 1.6 M sucrose, 3 mM MgCl2; 1 mM DTT) was placed under the homogenate. The mixture was then centrifuged at 3200g for 30 min at 4°C with the brakes set to “low” to separate the nuclei from the debris. The nuclei pellet was then washed sequentially with NF1 buffer and PBS, and resuspended in PBST with 5% BSA and 3% bovine serum. The nuclei were incubated with fluorescent antibodies (Abcam, ab196184) to the neuronal nuclear marker NeuN (at 100 ul per 1 million nuclei, dilution 1:500) overnight at 4°C in the dark. The next day, the nuclei were washed and resuspended in 1x PBS, and DAPI dye was added to a final concentration of 2 ug/ml. The nuclei were passed through a 35 μm cell strainer followed by fluorescence-activated nuclei sorting (FANS). The sorted NeuN-positive and NeuN-negative nuclei were collected in FANS buffer (1x PBS, 0.1% Tween-20, 5% BSA), spun at 2,300g for 10 min at 4°C, and washed in PBS. The collected nuclei were either snap-frozen for storage at −80 °C, or used immediately for Hi-C library preparation.

##### Hi-C library preparation

Hi-C experiments were conducted on cultured iPSC-derived neurons (whole cells) and adult neuronal and non-neuronal FANS-sorted nuclei. Two biological replicates were performed in each case. Approximately 1 million cells or nuclei per sample were used. We followed the previously described protocol (Pletenev et al. 2024; Lieberman-Aiden et al. 2009) for the subsequent steps. Hi-C libraries were sequenced on Illumina NovaSeq 6000 by Skoltech Genomics Core Facility and Evrogen Joint Stock Company.

##### Hi-C data processing

Raw reads of each Hi-C library were mapped to the human *hg38* reference genome with the distiller-nf pipeline, v.0.3.3 (Open2C et al. 2024), which uses BWA for genome mapping. The pipeline performed read filtering, binning, and generation of valid pair contact lists, which were subsequently converted into binned Hi-C interaction matrices. Sample metadata is detailed in the Supplemental materials (Supplemental Table S3).

Most of the Hi-C analyses were performed on Hi-C matrices merged by cell type. Merging was done at a 5-kb resolution using the "cooler.merge_coolers" function from the cooler library, v.0.8.11 (Abdennur and Mirny 2020). The resulting merged Hi-C maps were sampled to an equal number of contact pairs using the "cooltools.sample" function of the cooltools library, v.0.5.184. To avoid possible bias due to Hi-C ligation artifacts, values on the main diagonal were removed prior to sampling using cooler and custom Python scripts. The resulting .cool files containing the Hi-C matrices were ICE normalized (Imakaev et al. 2012) using the "cooler balance" command with default parameters.

#### Snm3C-seq data processing

To study 3D chromatin organization in different neural cell types at various developmental timepoints, we used publicly available snm3C-seq data (Supplemental Table S4). To generate aggregated contact maps, we downloaded files containing contact pairs per cell: ".3C.contact.tsv.gz" from GSE215353 and "_contacts.tar.gz" from GSE213950. Using the cell annotations provided in the metadata tables of the original studies, we combined cells into several groups. For the developing brain data (Heffel et al. 2024), we used cortical neurons divided into four stages: 2T, 3T, infant, and adult. Only neurotypical control samples were included. For adult brain data (Tian et al. 2023), we categorized cells by class: telencephalic excitatory neurons, inhibitory/non-telencephalic neurons, and non-neurons. Then, we further subdivided the classes by brain area: temporal cortex, frontal cortex, occipital cortex, paleocortex, basal nuclei, and basal forebrain. Three out of 18 resulting groups contained too few cells and were excluded from further analysis: non-neurons of the occipital cortex, inhibitory/non-telencephalic neurons of the paleocortex, and telencephalic excitatory neurons of the basal forebrain. For each group in both the adult and developing datasets, corresponding contact pairs were randomly sampled to match the number of pairs in the Hi-C data. The contact pairs were then aggregated into pseudobulk contact maps using the "cooler cload pairs" command from the cooler library, v.0.10.2. Consistent with Hi-C data processing, values along the main diagonal were removed, and the matrices were ICE normalized. Contact maps for each analyzed brain region and cell group are shown in Figure 1B. For subsequent analyses, the adult brain data from the Tian et al. (2023) dataset were merged across area and neuron types to ensure consistency with other samples that lacked this detailed stratification.

#### Downstream analysis inclusion criteria

In total, we collected 228 human and 89 mouse uniformly processed Hi-C and snm3C-seq maps (full metadata for all collected samples are provided in Supplemental Tables 1 and 3). To minimize batch effects in publicly available Hi-C and snm3C-seq datasets, we applied a sequential sample selection procedure that considered sequencing quality, library preparation metrics, cell type annotation, and experimental conditions. First, we retained only intact control samples (without any treatment or modifications) that represent a single cell type. Bulk tissue samples were excluded, except for fetal brain datasets, which are extremely scarce. Second, we filtered samples based on sequencing quality based on FastQC(Andrews 2010) reports generated by the distiller-nf pipeline (median Phred score ≥ 25, absence of highly overrepresented duplicates or singleton reads, and approximately uniform read coverage). Third, we verified sample annotations by examining consistency within each cell type using metadata from the original publications and PCA clustering of insulation score profiles. In this analysis, variability between cell types was expected to exceed variability within a given cell type. During this step, one neuronal sample (HSB181) from the Hu et al. (2021) dataset was identified as a mixture of neuronal and non-neuronal cells and was excluded from further analysis. Finally, we retained only samples and datasets in which Hi-C maps merged by cell type contained more than 250 million unique contact pairs for human samples and 200 million for mouse samples after removal of the main diagonal of the contact matrix, as reported by the distiller-nf pipeline. After filtering, 211 human and 83 mouse individual Hi-C maps were retained and subsequently merged by cell type to generate 62 human and 34 mouse Hi-C maps for downstream analyses. Number of merged samples and datasets used for the downstream analysis provided in Supplemental Table S5.

#### Contact probability analysis

To analyze distance-dependent contact probability, Hi-C maps binned at 5 kb resolution were processed using the "expected_cis" function from the cooltools library, v.0.5.1. This function computes the average intra-chromosomal contact frequency as a function of genomic distance, while applying smoothing and aggregation across all autosomes. For each Hi-C sample, resulting profiles were saved for subsequent analysis. Samples were grouped by cell type or developmental stage, and within each group, mean expected contact frequency was calculated at each genomic separation to obtain representative scaling curves. These average contact probability decay profiles were visualized as log-log plots, displaying contact probability as a function of genomic distance; where indicated, contact probabilities were normalized to the value at 10 kb. Chromosomes X, Y, and mitochondrial sequences were excluded from all analyses.

For derivative (scaling slope) analysis, the log-transformed contact frequency was differentiated with respect to the log of genomic separation using the numpy "np.gradient" function. Local minima and maxima in the scaling slope, corresponding to different hierarchical levels of chromatin organization, were identified with the "find_peaks" function from the scipy.signal module of SciPy v. 1.11.4 (Virtanen et al. 2020). Comparisons of slope extrema between groups were assessed using Mann–Whitney U tests.

#### Identification and analysis of TAD borders

To identify TADs from Hi-C data, we used the "insulation" module from the cooltools package, v.0.6.1 (Open2C et al. 2022). This module calculates the insulation profile using a diamond-window score approach. For each Hi-C map, insulation scores (IS) were computed with parameters chosen to optimize the balance between sensitivity and specificity in detecting TAD borders. Specifically, we used a resolution of 15 kb, a window size of 150 kb, a minimum valid pixel fraction of 0.75, and maintained a minimum distance of 4 bins from low-quality or missing bins.

To facilitate comparison of TAD borders across different Hi-C maps and to account for minor positional fluctuations, we implemented a clustering strategy based on the Density-Based Spatial Clustering of Applications with Noise (DBSCAN) algorithm from the scikit-learn package, v.1.5.2 (Pedregosa et al. 2018). Genomic coordinates of the initially identified TAD borders from the Hi-C maps were organized into clusters based on proximity. This distance threshold is set by our clustering parameters: a 15-kb resolution window and a clustering factor of 2, thus allowing boundaries within a 30-kb interval to be assigned to the same cluster. The "cityblock" distance metric and a minimum point threshold of 3 were used to define valid clusters. This approach accommodates minor positional variation across samples and enables us to identify TAD borders consistently across different samples. Borders present within the same cluster from different samples were treated as single entities in downstream comparative analyses. We identified sample-specific TAD borders by statistically comparing border strength values between groups using the Mann Whitney U test, defining borders as specific to a sample group when they showed significant differences in strength. GSEA for genes located at TAD borders, as well as other genes in this study, was performed in R using the "enrichGO" function from the clusterProfiler package (Yu et al. 2012).

##### TAD strength analysis

TAD strength, also known as domain score, was calculated following the approach of Flyamer et al. (2020) by generating rescaled pileups using the coolpuppy package, v.1.0.0 (Flyamer et al. 2020). In brief, each TAD along with flanking regions of equivalent size was extracted and rescaled into a 99x99 pixel matrix. TAD strength was then computed as the mean intensity in the middle 33x33 square, normalized by the mean intensity of the top and right 33x33 squares. For this analysis, we included only TADs located more than 10 bins away from any low-quality bins and under 1.5 Mb in length. TADs were considered shared between two maps if their borders were no more than 1.5 bins apart. We computed pairwise Jaccard similarities as the ratio of the intersection (number of common TADs) to the union of TADs, and then calculated Pearson's correlation coefficients between the strength scores of the common TADs.

For each TAD, we calculated border and corner enrichment scores. Corner enrichment was calculated as the $\log_2$ ratio of the intensity within a 7x7 square (20% of TAD pileup size) centered at 32x65 pixel (apex of the TAD triangle) to the inner region intensity. Border enrichment was computed as the $\log_2$ ratio of the mean intensity in 5x19 (reduced by a 2-pixel margin relative to the corner region *x* 60% of TAD pileup size) and 19x5 rectangles centered at 33x49 pixel and 49x33 pixels (the midpoints of the TAD boundaries) respectively to the inner region intensity. The intensity of the inner region was defined as an average of pixel values in a trapezoid separated from the TAD borders by a 1-pixel gap and by 4-pixel padding from the main diagonal. TADs were divided into four classes based on the presence of corner or border enrichment (TADs were considered enriched if the corresponding $\log_2$FC values were greater than 0). Genes, located within the boundaries of the TADs which were assigned to disjoint classes in iPSC-derived and postmortem samples were selected for additional analysis. The $\log_2$FC values of their gene expression between iPSC-derived and postmortem samples were tested for correlation with the difference in corner enrichment and border enrichment scores between iPSC-derived and postmortem Hi-C maps using scipy.stats.pearsonr from SciPy v. 1.11.4.

##### Identification of chromatin loops

To detect chromatin loop positions, we employed multiple Python libraries and custom scripts. The primary computational steps were performed using the cooler and cooltools packages. The analysis was conducted using the *hg38* assembly of the human genome. To estimate the baseline interaction frequencies, we generated expected contact maps using the "expected_cis" function from cooltools. This function calculates expected contact frequencies within chromosome arms, which serve as a reference for detecting significant loop interactions. To enhance the sensitivity and specificity of loop detection, we developed four custom kernels for the loop calling procedure. These kernels were then used with the cooltools package as masks to search for enrichment of interaction contacts. The kernels differed in the number of bins analyzed, ranging from narrow kernels for short-range interactions to broader kernels for long-range interactions. This approach improved the method's capacity to capture contact enrichment relative to the genomic background.

Chromatin loop identification was performed using the “dots” function from the cooltools package. Parameters used for loop calling included: a maximum genomic separation of 12,000,000 bp, a false discovery rate (FDR) threshold of 0.16, and tolerance for up to 5 bins of missing data. The clustering radius was set to 2.5 times the bin size to facilitate efficient grouping of proximal significant interactions. To ensure the robustness and uniqueness of the identified loops across various kernel configurations, we created custom functions for loop comparison and filtering. These functions utilized the pybedtools package (Dale et al. 2011) to perform pairwise comparisons between loop sets obtained with different kernels.

##### Comparisons of chromatin loops between samples

Due to limitations of automated annotation procedures, the position of chromatin loops often varies slightly between samples. To enable robust comparisons between samples, we developed a strategy that clusters loops automatically detected in individual samples and treats them as identical if they are spatially proximate. Specifically, we used the DBSCAN function from the scikit-learn package to group loops located within 2.5 bins (37.5 kb) of each other at 15 kb resolution. Loops located within the same cluster were considered equivalent, facilitating the identification of consistent looping patterns across datasets. For analyses requiring a complete matrix of loop intensities across all samples, such as PCA and correlation analyses, we assigned an intensity value to every cluster in every sample. If a sample did not have a loop annotated by the detection algorithm in a given cluster, we estimated the loop intensity by searching within a 3 by 3 bin window centered at the cluster coordinates and selecting the region with the highest median intensity.

For other analyses, we further refined the clusters through a series of filtering steps to ensure data reliability. Technical filters were applied to exclude clusters lacking sufficient data points, and intensity-based thresholds were imposed. Clusters with fewer than a predetermined number of data points or with intensities below threshold were excluded from downstream analyses. The final set of clusters was used for subsequent analyses. We intersected loop anchors with other genomic features provided in .bed or .bedpe files using the pybedtools package.

For differential chromatin loop analysis, we aimed to identify loops that are upregulated or downregulated between two groups: "postmortem" and "iPSC-derived" neurons. We employed an approach based on statistical assessments of loop intensities. First, the median loop intensity for each cluster was calculated within each group. Then, the Mann-Whitney $U$ test was applied to identify significant intensity differences. Loops demonstrating a statistically significant increase in the "postmortem neurons" group were classified as upregulated, while those with a significant decrease were considered downregulated.

##### Transcriptomics data processing and differential gene expression analysis

Publicly available RNA-seq datasets for postmortem adult and fetal brain samples, as well as iPSCs, NPCs, and iPSC-derived neurons, were used (Supplemental Table S6). Sequencing adapters were removed from raw FASTQ files using the Trimmomatic package, v.0.39 (Bolger et al. 2014), with the following parameters: "ILLUMINACLIP:NexteraPE-PE.fa:4:30:10", "LEADING:3", "TRAILING:3", "MINLEN:45".

Trimmed FASTQ files were processed using the Salmon tool, v.1.10.0 (Patro et al. 2017), with the parameters "--validateMappings" and "-l A", using indexed genome *hg38* (for human data) or mm10 (for mouse data). Only protein-coding genes were included in subsequent analyses. GENCODE gene annotation, v.41 (Frankish et al. 2021) was used. Transcript abundance (TPM) and regularized log-transformed (rlog) expression values were obtained using the tximport (Soneson et al. 2016) and DESeq2 (Love et al. 2014) packages, respectively. Rlog normalization was performed with the blind = TRUE setting, meaning that the transformation did not account for experimental design or condition-specific effects.

For downstream analyses, when gene expression data included biological replicates within a given cell group, rlog values were averaged across replicates to generate a single representative expression profile per group.

Differential gene expression analysis was performed using the DEseq2 package, v.1.36.0. Genes were retained if they had at least five read counts in at least 10 samples. Three variables (dataset, cell type and donor ID) were included in the DESeq2 design formula. Genes with an adjusted $p$-value $\leq 0.01$ and $|\log2FoldChange| \geq 2$ were considered differentially expressed and used for further analysis. Gene Ontology enrichment analysis was performed on protein-coding genes using the clusterProfiler package, v.4.4.4 (Yu et al. 2012; Wu et al. 2021a), with "enrichGO" function and the following parameters: "OrgDb = org.Hs.eg.db", "pAdjustMethod = "BH"", "pvalueCutoff = 0.05".

##### Principal component analysis (PCA) of Hi-C and gene expression data

To analyze the relationships between cell types, we performed PCA on genome-wide chromatin insulation scores and gene expression profiles derived from both mouse and human samples. Insulation scores for each sample were determined from Hi-C data as described above, producing a genome-wide profile per sample. For gene expression, normalized matrices were generated from RNA-seq data. PCA was carried out in Python using the PCA module from scikit-learn, v.1.5.2 (Pedregosa et al. 2018). Prior to PCA, all input matrices were scaled and centered to ensure comparability across samples. The first two principal components were retained for subsequent visualization, as they captured the largest fraction of variance in the data.

To visualize the results, PCA coordinates were plotted with the ggplot2 package, v. 3.5.1. Each sample was colored according to its assigned group. Ellipses denoting the 95 percent confidence interval for each group were calculated in an unsupervised manner with the "stat_ellipse" function in ggplot2, based on sample distributions in principal component space. Arrows in Figure 1B were added manually to indicate key differentiation trajectories and to help guide visualization of progression between cell types.

#### Analysis of long-range PcG-contacts

The coordinates of long-range PcG-contacts were obtained from Pletenev et al. (2024). Contact frequencies, extracted from 100-kb resolution Hi-C matrices, were used for PCA and scaling plots. The mean heatmaps centered on long-range PcG-contacts were generated using 20-kb scaling-normalized Hi-C maps by averaging contact map fragments across all long-range PcG-contacts, followed by per-group averaging.

To validate the low prevalence of long-range PcG-mediated contacts in iPSC-derived neurons, we built upon previous observations that such contacts occur between extended genomic regions marked by H3K27me3 in neurons. We generated average heatmaps centered on all possible intrachromosomal interactions between bins containing H3K27me3 ChIP-seq peaks greater than 10 kb in width and separated by more than 3 Mb. Heatmaps were created at 10-kb resolution, using cell type–matched ChIP-seq and Hi-C datasets, ensuring an unbiased comparison. Because ChIP-seq peak quality varied between samples, resulting in different numbers of broad H3K27me3 peaks and potentially influencing the analysis, we produced two versions of average heatmaps for iPSC-derived neurons, based on datasets with different quality metrics (Ciceri et al. 2024; Wu et al. 2021b). For postmortem neurons, data from Dong et al. (2022) were used. Group-level heatmaps were generated by averaging per-sample mean Hi-C maps within the iPSC-derived and postmortem groups.

To examine how PcG-contacts relate to gene expression, we analyzed transcriptomic data from postmortem neurons isolated from the dorsolateral prefrontal cortex and anterior cingulate cortex (Dong et al. 2024; Rizzardi et al. 2019), as well as from iPSC-derived neurons (Ballarino et al. 2022; Li 2016; Lu et al. 2020; Zaghi et al. 2023; Ciceri et al. 2024). For the transcriptomic data from Ciceri et al. (2024), we used a single time point corresponding to 50 days of differentiation, as the authors reported that the most pronounced transcriptional changes occurred between days 25 and 50, indicating that neurons at day 50 represent a sufficiently differentiated state. Genes were assigned to contact anchors if their TSSs fell within 100-kb bins overlapping H3K27me3 peaks identified in postmortem neurons in Dong et al. (2024). This step was based on the idea that long-range PcG-contacts tend to form between H3K27me3-enriched regions. PcG-contacts involving loci with low gene expression in both iPSC-derived and postmortem neurons were selected, based on the gene expression level (TPM<1) at both 100-kb anchor bins within H3K27me3 peaks from postmortem neurons (Dong et al. 2024). In contrast to rlog units, TPM values could be unevenly distributed across samples. Therefore, we first determined the rlog value corresponding to 1 TPM in postmortem neurons and then used this value as a threshold to exclude highly expressed regions in both iPSC-derived and postmortem neurons.

To assess PcG-contact intensity across gene expression categories, genes assigned to PcG-contact anchors were annotated based on differential expression analysis of the full protein-coding transcriptome (see transcriptome analysis section), comparing iPSC-derived and postmortem neurons. Genes were grouped into three categories: upregulated, downregulated, or not significantly differentially expressed. PcG-contacts were then stratified into three groups according to anchor composition. Each group included contacts where at least one anchor contained genes from only one expression category, with neither anchor overlapping genes from the other two categories.

To analyze compartmental transitions associated with PcG-contacts, we used PC1 values derived from compartment annotations. Only genomic bins with an absolute PC1 value greater than 0.05 were included to minimize ambiguous assignments. Average heatmaps were generated for PcG-contacts where both anchors were assigned to the same category of compartment transition. In the analysis of the relationship between compartment transitions and gene expression, consistent with the properties of A- and B-compartments (Lieberman-Aiden et al. 2009) (gene-rich and gene-poor), the average number of genes per bin varied across transitions (0.33 for B→A, 0.85 for A→A, 0.68 for A→B, and 0.38 for B→B). To account for potential bias introduced by unequal group sizes, we additionally compared random gene subsets of equal size from the A→A and B→B transitions. The expression distribution and number of genes in these subsets were matched to those of the B→A group in postmortem neurons. Using these expression-matched groups, we calculated the p-value for the difference in gene expression between iPSC-derived and postmortem neurons.

A clustermap of Polycomb group genes was generated using rlog-normalized expression values. Hierarchical clustering was performed using correlation distance and average linkage (metric="correlation", method="average").

To compare H3K27me3 levels within PcG-contact anchors between iPSC-derived and postmortem neurons, ChIP-seq peaks from each sample were categorized into two groups: those within PcG-contact anchor bins and those outside. For each sample, the mean peak score (MACS3 fold enrichment) within anchor bins was divided by the mean score outside the anchors, yielding a relative measure of H3K27me3 enrichment at PcG-contacts. The analysis was repeated using median peak scores to test robustness. Since ChIP-seq quality could influence enrichment estimates, we also assessed the relationship between enrichment ratios and standard quality control metrics, including FRiP score, relative strand cross-correlation (RSC), and normalized strand cross-correlation (NSC) (Supplemental Fig. S29H). Additional analyses of transcription and H3K27me3 levels were performed using the combined set of ChIP-seq peaks from both cell types overlapping PcG-contact anchors. For genes with TSSs within these peaks, rlog expression values were used to assess the association between H3K27me3 occupancy and gene expression.

#### FIREs and SNP analysis

Positions and scores of FIREs, defined as the total number of cis-interactions within the nearest 200 kb for each 10-kb bin passing default filters, were identified using the FIREcaller package (Crowley et al. 2021). FIREs were detected independently for each sample, using default parameters, with mappability files specifying cut frequencies of the corresponding restriction enzymes as required input. A union set of FIRE positions across all samples was compiled, and the corresponding FIRE scores were used for PCA and subsequent analyses. PCA revealed a pronounced difference between iPSC-derived neurons from Lu et al. (2020) and all other samples (Supplemental Fig. S30A). This difference likely reflects use of the easy Hi-C protocol in that dataset, as opposed to conventional Hi-C used in other datasets; thus, this sample was excluded from downstream analyses.

PCA also showed that iPSC-derived neurons clustered closely with neural progenitor cells (NPCs) from the same studies. To test whether this reflects batch effects rather than biological similarity, we performed a pairwise enrichment analysis. Specifically, for each sample from the iPSC, NPC, fetal, and postmortem groups, we calculated the average cis-interactions at the genomic positions corresponding to FIREs detected in one of the iPSC-derived neuron samples. This value was then normalized to the average cis-interactions across all other FIRE positions from the same reference sample (belonging to iPSC, NPC, fetal or postmortem groups). The resulting ratio served as a measure of how strongly FIREs from iPSC-derived neurons were pronounced in other cell types. For each cell group containing *n* samples this yielded $n \times 7$ enrichment values, where 7 corresponds to the number of iPSC-derived neuron samples.

For the differential FIREs analysis, we focused on two cell groups: iPSC-derived and postmortem neurons. First, adjacent FIREs detected in any sample from either group were merged into composite regions. FIRE scores were then averaged per sample across each merged region, to produce a set of custom "superFIREs" with corresponding per-sample scores. Next, these scores were used as input for differential analysis using the limma package (Ritchie et al. 2015), with the trend = FALSE setting and BH-correction applied to adjust *p*-values. To account for potential technical bias due to differences in restriction enzymes, this factor was included as a covariate in the linear model. Regions with adjusted *p*-values below 0.05 were considered differentially enriched and were referred to as differential FIREs.

Genes were assigned to FIREs based on the location of their TSSs. For the postmortem group, only samples from cortical regions were included into analysis. Gene Ontology enrichment analysis was performed using the enrichGO function from the clusterProfiler package, with both *p*- and *q*-value cutoffs set to 0.05. The background gene set included all genes with a mean TPM > 1 across the respective group (either iPSC-derived or postmortem neurons). Enhancer coordinates were obtained from two sources: GeneHancer v5.18 for neural stem/progenitor cells (NSPCs) (Fishilevich et al. 2017) and Dong et al. (2022) for postmortem neurons. From the latter, only enhancers not labeled as "glia" were retained. Because GeneHancer does not distinguish enhancers from NPCs alone but aggregates data for both neural stem and progenitor cells under the NSPC label, we used NSPCs as a proxy for NPCs and referred to them accordingly in the Results section. Enhancers were linked to FIREs based on coordinate overlap. P-values for the association between postmortem-specific FIREs and enhancers were calculated from the following information: the number of postmortem enhancers falling into postmortem-specific FIREs (473), the number of NSPC enhancers falling into postmortem-specific FIREs (340), the number of postmortem enhancers falling into the rest FIREs (common and iPSC-derived-specific) (3023) and the number of NSPC enhancers falling into the rest FIREs (common and iPSC-derived-specific) (4791). Similarly, the association between iPSC-derived-specific FIREs and enhancers was estimated from the following information: the number of postmortem enhancers falling into iPSC-derived-specific FIREs (84), the number of NSPC enhancers falling into iPSC-derived-specific FIREs (403), the number of postmortem enhancers falling into the rest FIREs (common and postmortem-specific) (3412) and the number of NSPC enhancers falling into the rest FIREs (common and postmortem-specific) (4728).

Average heatmaps centered on FIREs were generated using 10-kb scaling-normalized Hi-C data as follows. First, adjacent FIREs were merged into super-FIREs. For each super-FIRE, we extracted a Hi-C map fragment encompassing the super-FIRE region and 10 flanking bins on each side. Each fragment was then rescaled by averaging the signal within the FIRE region while retaining the adjacent bins unchanged. Finally, mean Hi-C heatmaps were produced by averaging the rescaled fragments across all super-FIREs, followed by group-wise averaging.

Correspondence between FIREs and chromatin loops was assessed by intersecting aggregated loop coordinates with super-FIRE regions. UpSet plots illustrating GWAS SNP cross-disorder overlap and FIREs-loop intersections were generated using the UpSetPlot Python package.

To analyze chromatin interactions of genomic bins containing SNPs associated with neurological disorders, we used variants from the following studies: Grove et al. (2019) - for autism spectrum disorder (ASD), Pardinas et al. (2018) - for schizophrenia (SCZ), O'Connell et al. (2025) - for bipolar disorder (BD), Adams et al. (2025) - for major depressive disorder (MDD), and Wightman et al. (2021) - for Alzheimer's disease (AD). These variants were fine-mapped in the respective studies and were used in our analysis as credible SNPs. Specifically, for MDD, we selected SNPs with a posterior inclusion probability > 0.5, since a list of SNPs associated with BD was presented for this threshold.

SNP coordinates were retrieved by rs IDs from Ensembl using R package biomart for ASD, SCZ, MDD and BD. For Alzheimer's disease, rsIDs were obtained based on SNP coordinates reported in the hg19 genome build and subsequently converted to hg38 coordinates. Significant SNP-associated chromatin interactions were identified at 20-kb resolution in each sample, following the approach from Won et al. (2016), with minor modifications. Specifically, we fitted the interaction data using an exponentially modified Gaussian distribution, instead of the Weibull distribution used in the original study. We also estimated background interaction frequency from all genomic bins at a given distance, rather than considering only bins that contain GWAS-imputed SNPs. For each SNP-containing bin, we analyzed chromatin interactions within a ±5 Mb flanking region, excluding those separated by less than 40 kb. We then calculated the mean interaction intensity within a 3 × 3 bin window centered on each SNP-associated contact that was detected as significant in at least one sample. These averaged values were subsequently used for principal component analysis (PCA) and differential interaction detection. Differential analysis between iPSC-derived and postmortem neurons was conducted, using the limma package with trend = FALSE and BH-correction (adjusted $p < 0.05$). To ensure all samples are unpaired between the two groups and the number of samples is equal, we excluded one iPSC-derived sample from the Rahman et al. (2023) dataset from the differential analysis. Average heatmaps of shared and differentially enriched SNP-associated interactions were generated using scaling-normalized Hi-C maps at 20-kb resolution. For individual SNPs, we computed mean interaction profiles using 10-kb resolution Hi-C maps. For the GWAS locus, which contains 15 SCZ-associated SNPs linked to the *CHRNA2* gene and spanning several 10-kb bins, we averaged interaction profiles across all corresponding bins to obtain the final contact profile.

All group-level heatmaps and interaction profiles were produced by averaging per-sample means within each group.

##### Cell type proportion estimation and expression in different cell types.

Cell type proportions for four iPSC-derived samples from Li (2016), Lu et al. (2020), Zaghi et al. (2023), and Ballarino et al. (2022) were estimated from transcriptomic data using MuSiC (Wang et al. 2019) and CIBERSORT (Chen et al. 2018). For MuSiC deconvolution, snRNA-seq data from Jeffries et al. (2025) was used (specifically, adult samples from batches 210430, 210209, and 200128). Only cells meeting quality thresholds were included: total UMIs > 500, the number of expressed genes between 500 and 2500, and mitochondrial gene percentage < 5%. Decomposition relied on a gene signature compiled from multiple sources: McKenzie et al. (2018), Mesecar et al. (2023), and the Human Protein Atlas (Uhlén et al. 2015). From the first source, genes present in both the “top human enriched” and “top human specific” lists were selected. As the study McKenzie et al. (2018) represents a meta-analysis, genes with average global mean > 2 and $log_2FC > 1$ from the two included studies (Zhang et al. (2016) and Darmanis et al. (2015) were additionally selected from the initial list. From Mesecar et al. (2023), genes with $log_2FC > 4$ were chosen (limited to cell markers for excitatory neurons, inhibitory neurons, astrocytes, oligodendrocytes, OPCs, microglia, and endothelial cells). From the Human Protein Atlas, “cell type enhanced” genes related to excitatory neurons, inhibitory neurons, and OPCs were selected. The three gene lists were unified, and genes assigned to multiple cell types were removed, yielding a final signature of 1902 genes. To validate the signature's suitability, the quality of snRNA-seq cell type annotations was assessed using UMAP visualization (Supplemental Fig. S3C). For CIBERSORT, the gene signature based on Human Cell Atlas transcriptomic data (Hodge et al. 2019) was sourced from Sutton et al. (2022).

snRNA-seq data from Jeffries et al. (2025) were also used to estimate expression of genes carrying SNPs and genes interacting with SNP locus in neurons over different cell types. To this purpose, pseudobulk normalized expression was calculated over data from batches 210430, 210209, and 200128 , filtered as described above, with the Seurat package.

##### ChIP-seq analysis

ChIP–seq datasets were obtained from multiple sources (Supplemental Table S7). For postmortem neurons, we selected data generated from the dorsolateral prefrontal cortex (DLPFC), corresponding to the brain region represented in the Hi-C datasets. The dataset from Kozlenkov et al. (2018). included two neuronal subtypes: medial ganglionic eminence (MGE)-derived inhibitory GABAergic interneurons and excitatory glutamatergic neurons. ChIP-seq data for postmortem fetal neurons were not included in the analysis because, to our knowledge, such datasets are not currently available, including those accessible upon request.

Basic processing steps, including broad peak calling and generation of bigWig signal tracks, were performed using the nf-core/chipseq pipeline (2024) with the *hg38* genome assembly. For downstream analyses, bigWig files were converted to bedGraph format, and average signal intensity was computed for each 5-kb genomic bin for H3K27me3. To normalize ChIP–seq signals, we divided each signal track by its corresponding input track, using a pseudocount defined as the minimum non-zero value across all datasets for the given histone mark. For datasets of iPSC, NPC, or iPSC-derived cells that included replicates for both signal and input, we first merged replicate tracks by calculating the mean signal prior to normalization. Average signal profiles around PcG-contact anchors (shown in main figures) were then generated by averaging normalized ChIP–seq tracks from the highest-quality datasets: Wu et al. (2021b) and Ciceri et al. (2024) for iPSC-derived neurons and Kozlenkov et al. (2018) for postmortem neurons.

For H3K27ac, used as a marker of active enhancers, we followed a similar procedure, using 10-kb bins to match the resolution used for FIREs detection. To exclude promoter-associated signals, we masked bins in the H3K27ac track that overlapped with H3K4me3 narrow peaks.

To highlight local enrichment, zero-centered average profiles shown in the main figures were adjusted by subtracting the signal baseline. Specifically, for H3K27me3, the baseline was calculated as the mean signal within 50 kb from both ends of the profiled region; for H3K27ac, a 40-kb window was used. Integrative analyses presented in the main figures were performed on data averaged within four major groups (iPSCs, NPCs, iPSC-derived neurons, and postmortem neurons) without matching samples by individual study. For additional confirmation, we also calculated similar non-zero-centered individual profiles for each sample (Supplemental Fig. 29F, 30G).

##### Colocalisation of AD-associated SNPs and candidate cis-regulatory elements

To estimate occupancy of cCREs with AD-associated SNPs, cCREs coordinates generated for different cell types were retrieved from Liu et al. (2025). The genomic coordinates of cCREs and AD-associated SNPs were intersected using pybedtools package.

### Supplemental Figures

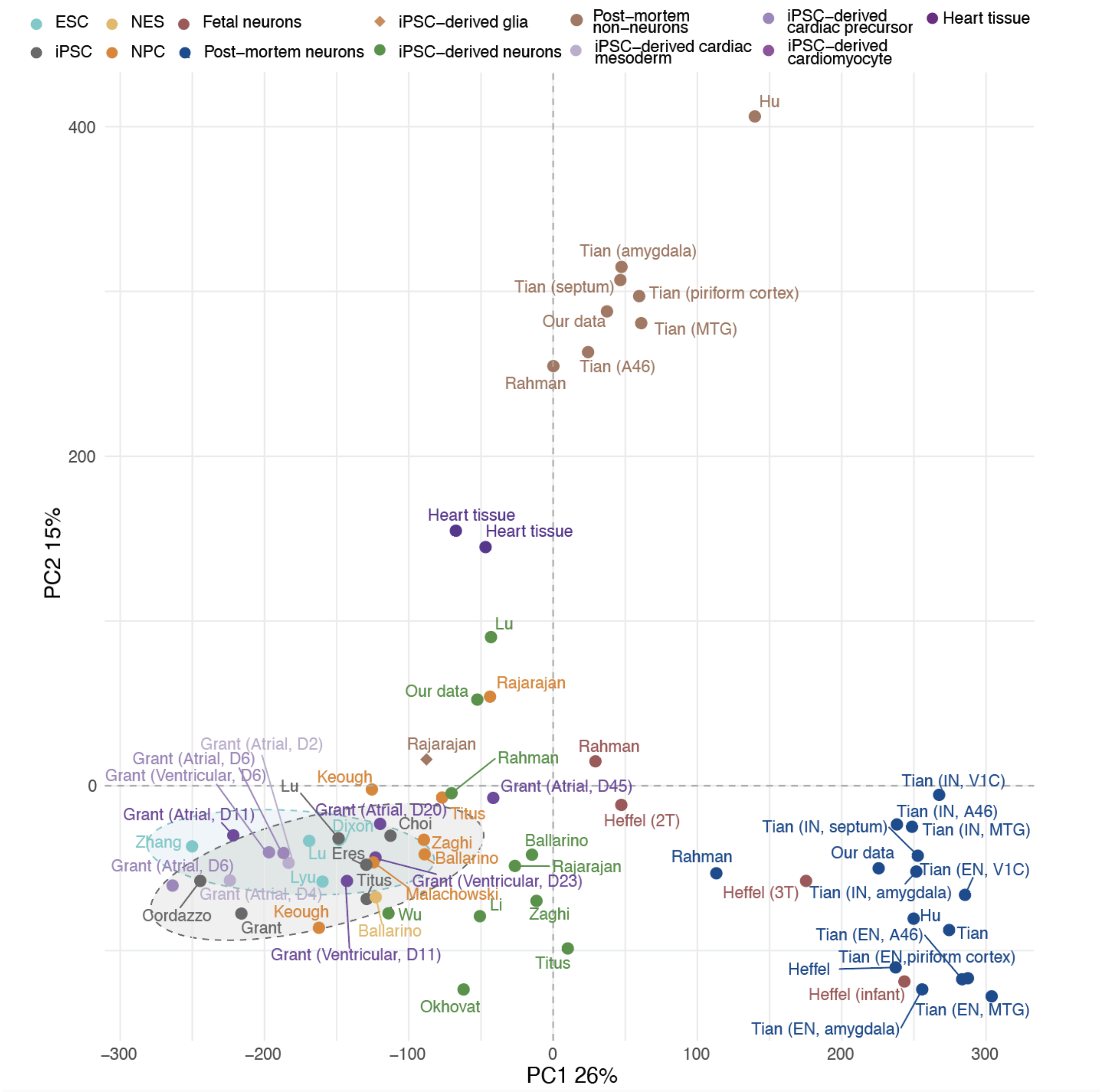


**Supplemental Fig. S1. PCA plots of Hi-C maps illustrating insulation score changes in human samples (including "Postmortem non-neurons" group).** Abbreviations for brain regions and neuron types are as follows: MTG, middle temporal gyrus (temporal cortex); V1C, primary visual cortex (occipital cortex); A46, prefrontal cortex (Brodmann area 46). EN and IN are used to indicate the neuron type, i.e., excitatory (EN) or inhibitory (IN), with the brain region specified in parentheses.

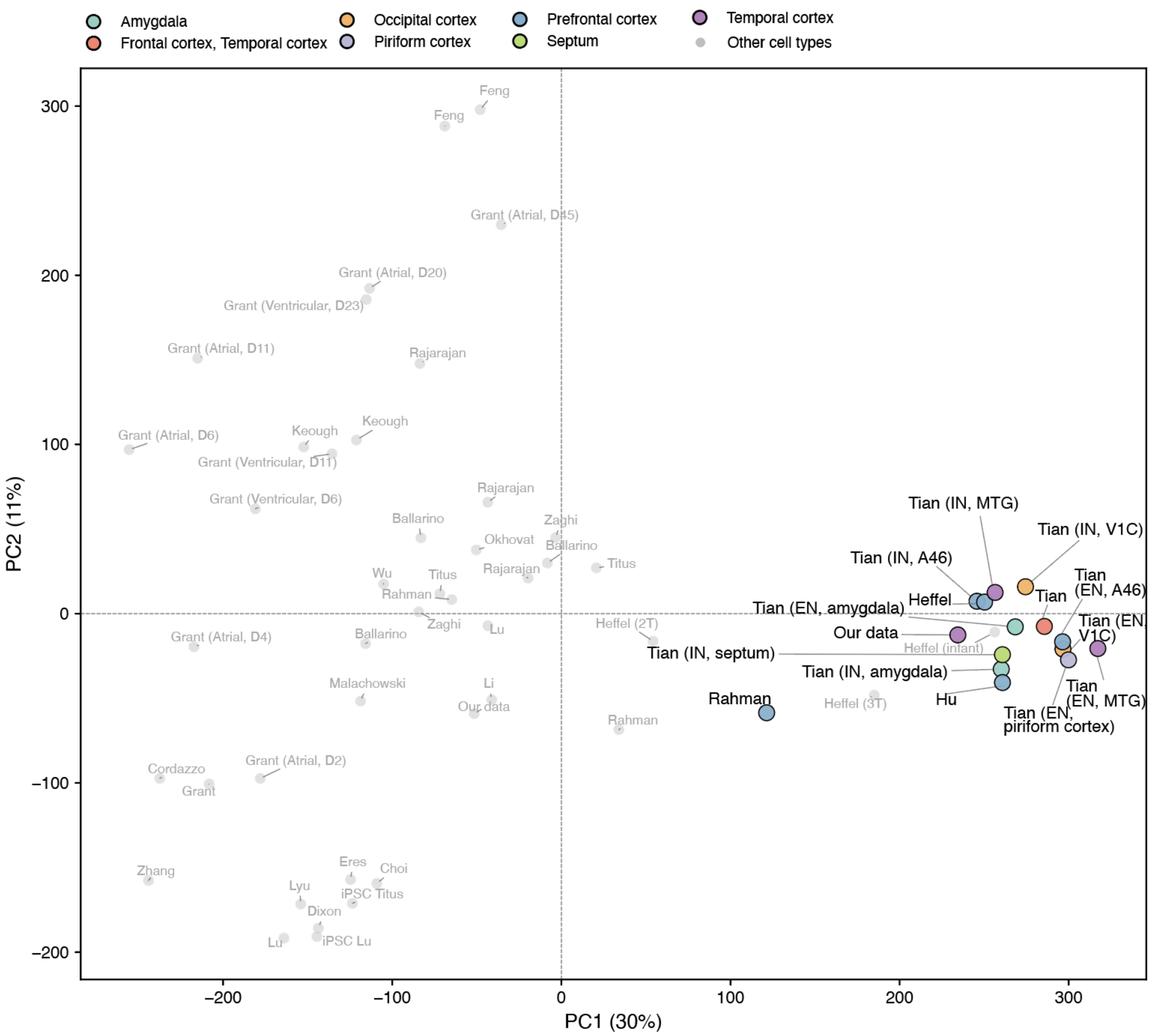


**Supplemental Fig. S2. PCA plot of Hi-C maps illustrating insulation score variation in human samples with postmortem neurons colored by brain region.** Abbreviations for brain regions and neuron types are as follows: MTG, middle temporal gyrus (temporal cortex); V1C, primary visual cortex (occipital cortex); A46, prefrontal cortex (Brodmann area 46). EN and IN are used to indicate the neuron type, i.e., excitatory (EN) or inhibitory (IN), with the brain region specified in parentheses.

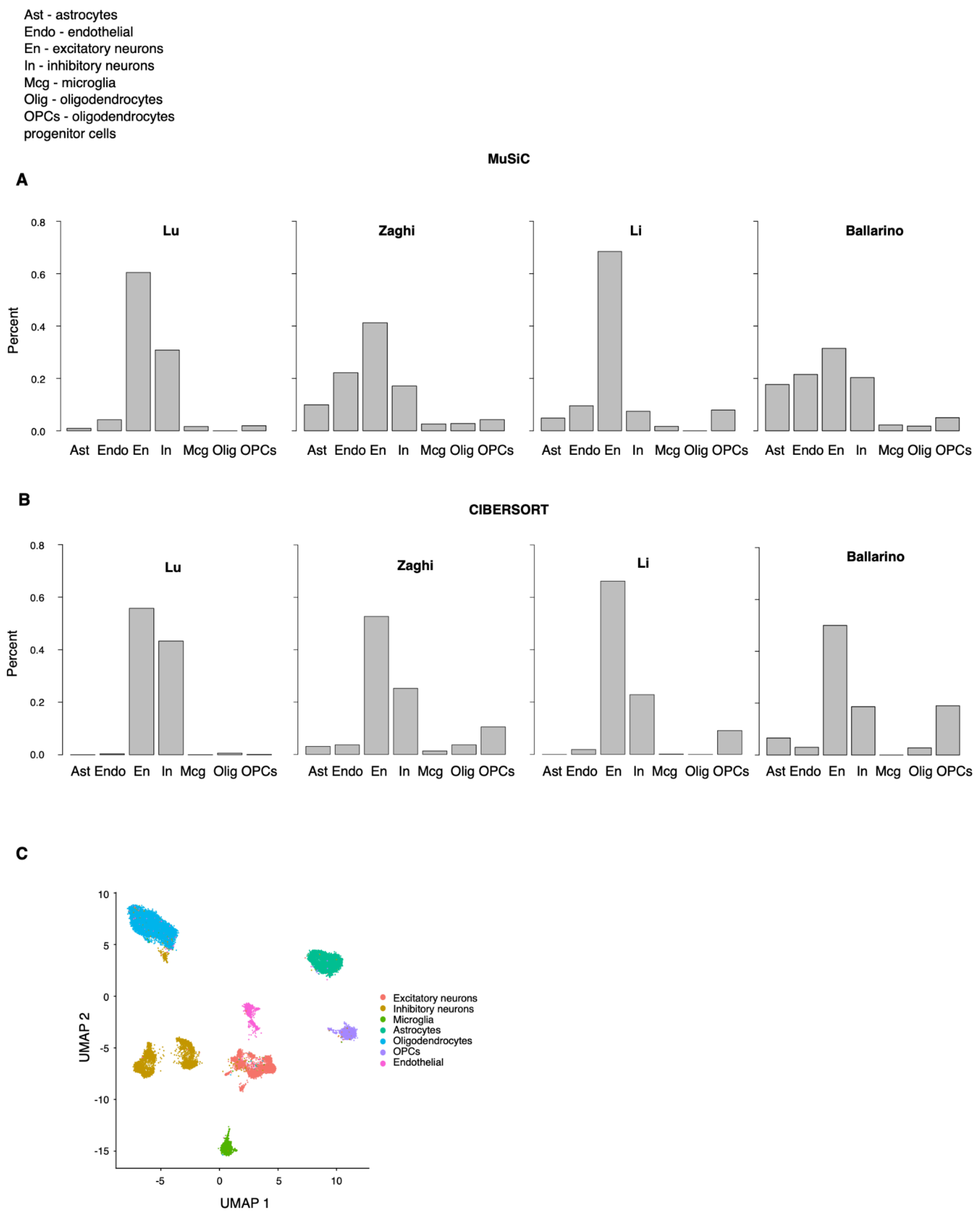


**Supplemental Fig. S3. Cell type composition of transcriptomic data from iPSC-derived samples.** (A–B) Cell type proportions estimated by decomposition using MuSiC (Wang et al. 2019) and CIBERSORT (Chen et al. 2018). (C) Dimensionality reduction of snRNA-seq nuclei performed using the gene signature applied for MuSiC decomposition.

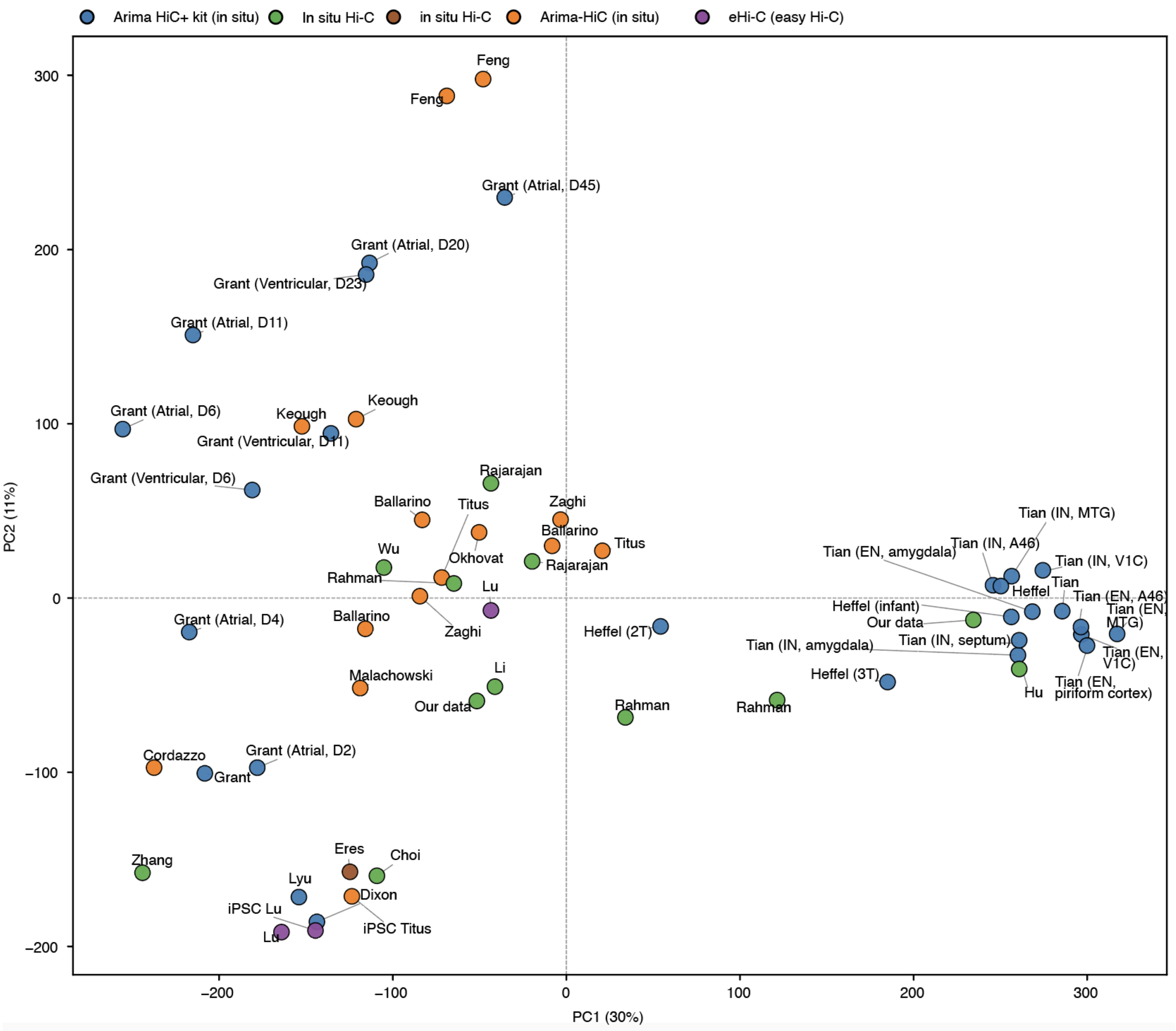


**Supplemental Fig. S4. PCA plot of Hi-C maps illustrating insulation score variation in human samples with all samples colored by Hi-C protocol used in the study.** Abbreviations for brain regions and neuron types are as follows: MTG, middle temporal gyrus (temporal cortex); V1C, primary visual cortex (occipital cortex); A46, prefrontal cortex (Brodmann area 46). EN and IN are used to indicate the neuron type, i.e., excitatory (EN) or inhibitory (IN), with the brain region specified in parentheses.

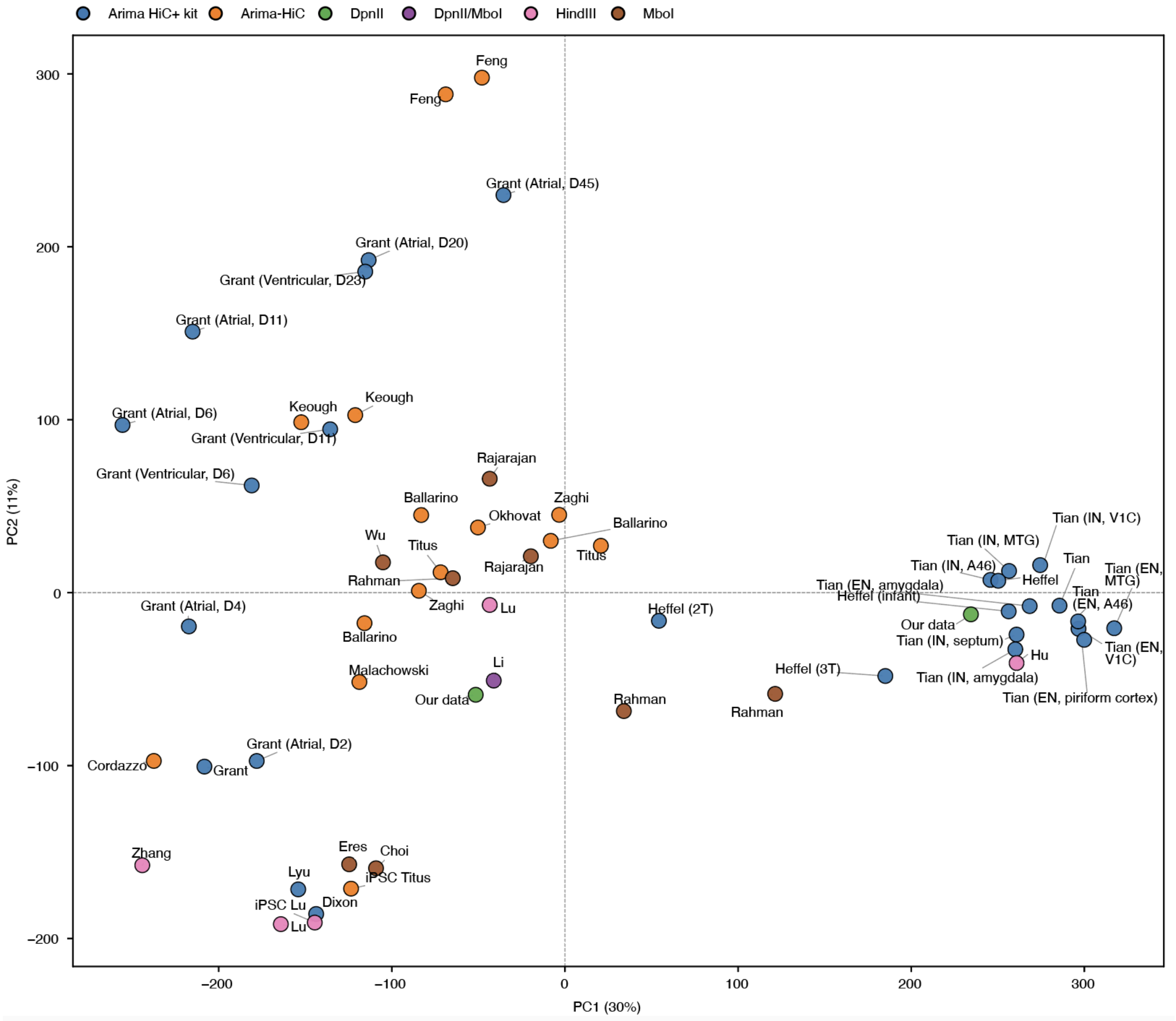


**Supplemental Fig. S5. PCA plot of Hi-C maps illustrates insulation score variation in human samples with all samples colored by restriction enzymes used in Hi-C protocols.** Abbreviations for brain regions and neuron types are as follows: MTG, middle temporal gyrus (temporal cortex); V1C, primary visual cortex (occipital cortex); A46, prefrontal cortex (Brodmann area 46). EN and IN are used to indicate the neuron type, i.e., excitatory (EN) or inhibitory (IN), with the brain region specified in parentheses.

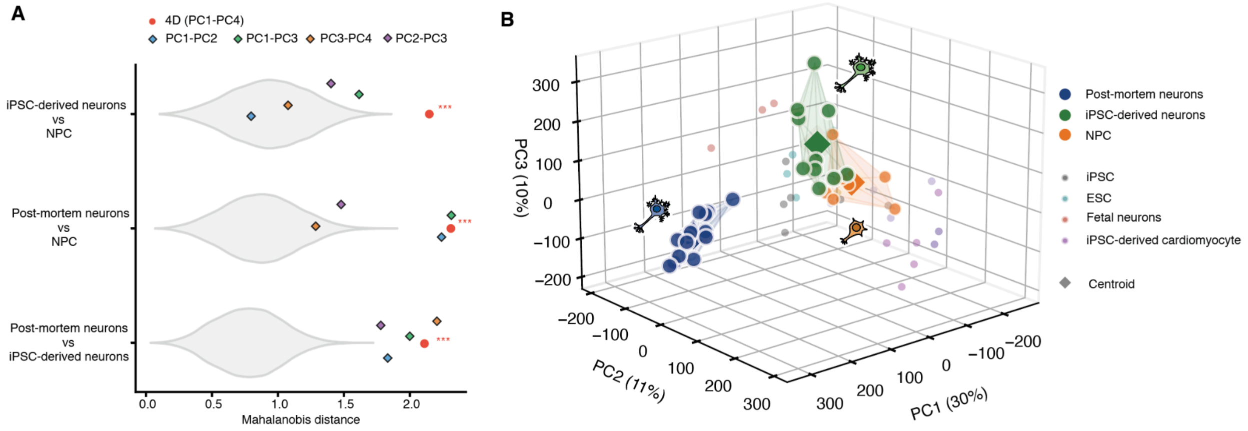


**Supplemental Fig. S6. Separation of postmortem and iPSC-derived neurons in a higher-dimensional space.** (A) Higher-dimensional separation analysis using Mahalanobis distances. Violin plots show the null distribution of pairwise distances between postmortem neurons, iPSC-derived neurons, and NPCs based on 10,000 permutations (distribution of distances expected by chance). Red circles indicate observed Mahalanobis distances calculated in 4-dimensional space (PC1-PC4). Colored diamonds show distances in 2-dimensional PC subspaces: PC1-PC2 (blue), PC3-PC4 (orange), PC1-PC3 (green), and PC2-PC3 (purple). Asterisks denote statistical significance from permutation testing (*** – $p < 0.001$). (B) Three-dimensional PCA visualization of cellular diversity. Main neuronal populations are highlighted: postmortem neurons (blue circles), iPSC-derived neurons (green circles), and NPCs (orange circles). Colored convex hulls delineate the spatial boundaries of the three main neuronal clusters. Diamond markers indicate group centroids calculated as the mean position in PC1-PC2-PC3 space.

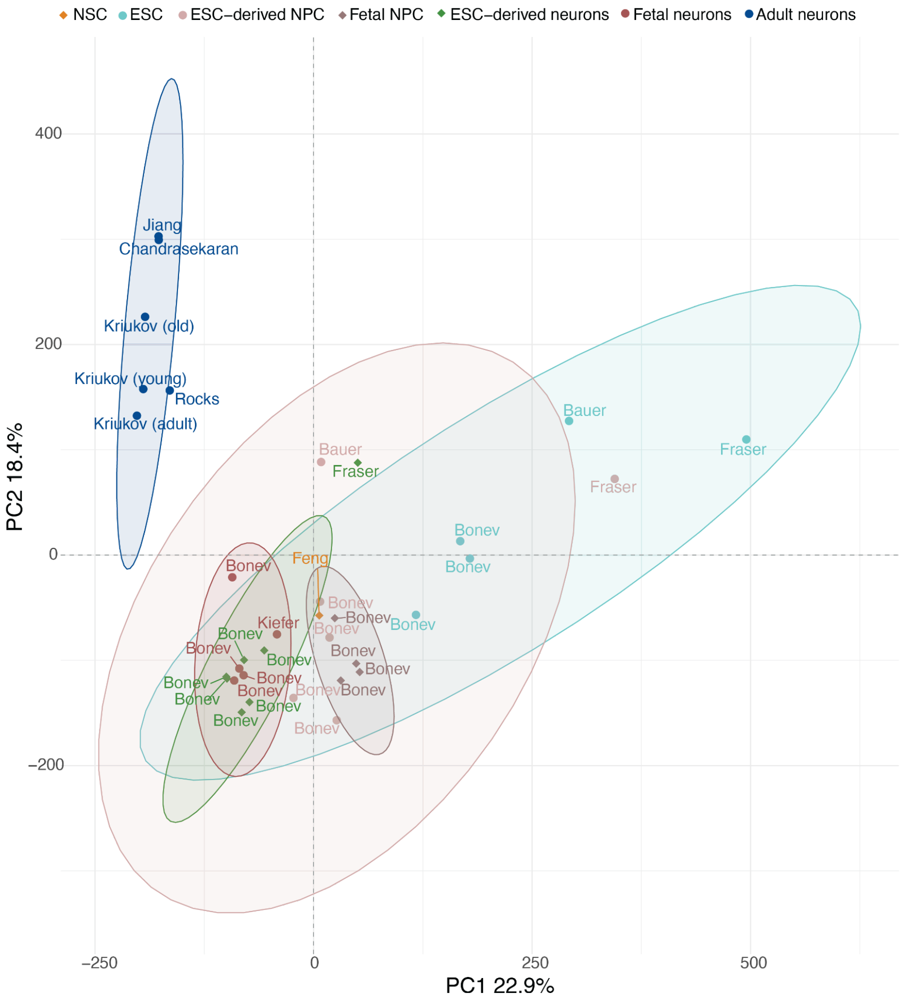


**Supplemental Fig. S7. PCA plots of Hi-C maps illustrating insulation score changes in mouse samples.**

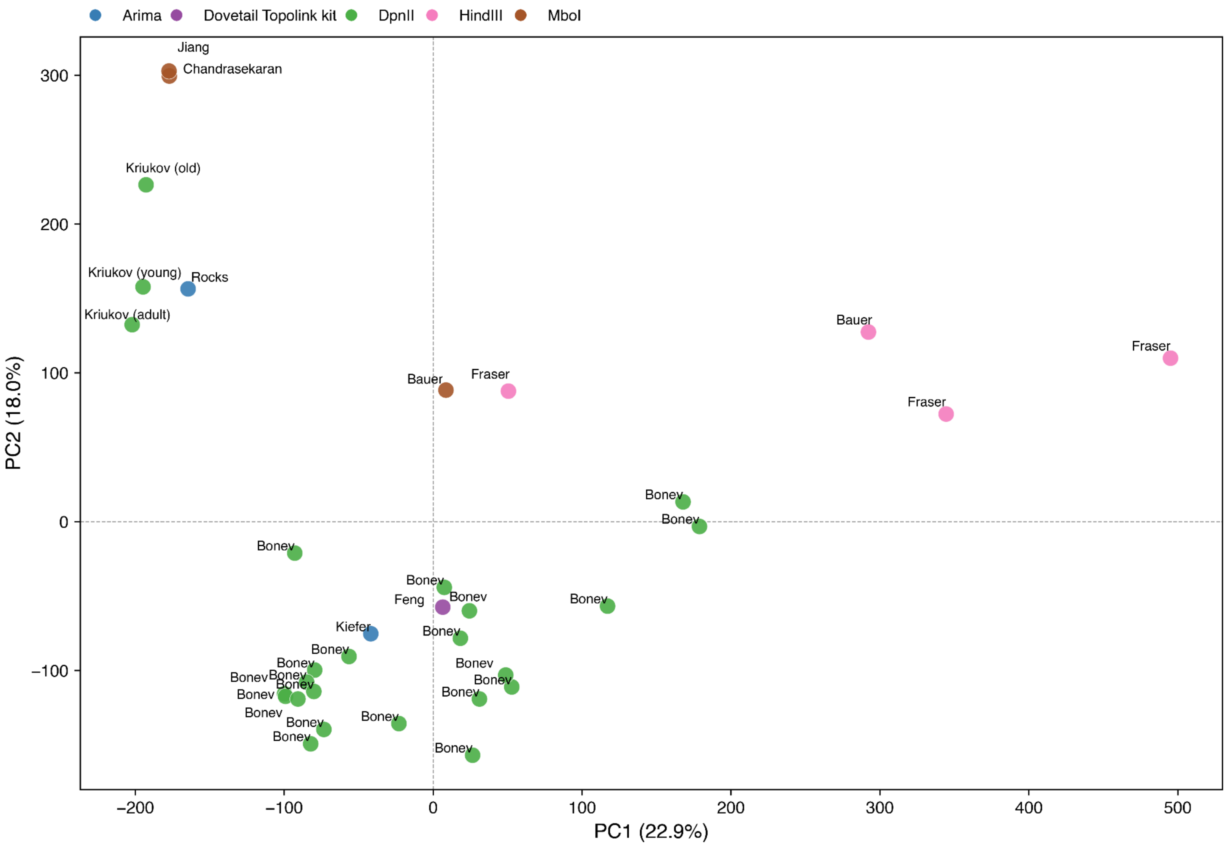


**Supplemental Fig. S8. PCA of Hi-C maps showing variation in insulation scores across mouse samples.** Samples are colored according to the restriction enzyme used in the Hi-C protocol. All datasets were generated using *in situ* Hi-C.

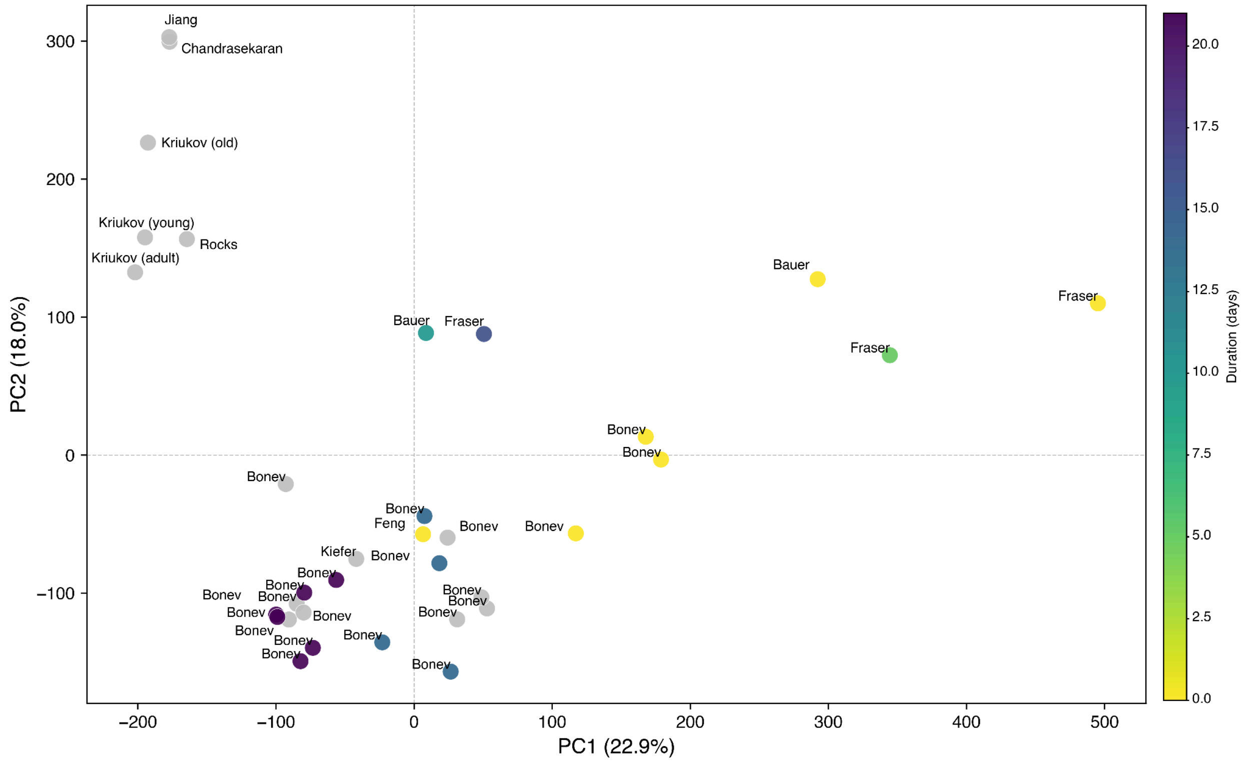


**Supplemental Fig. S9. PCA of Hi-C maps showing variation in insulation scores across mouse samples.** Samples are colored by the duration of the differentiation protocol in days.

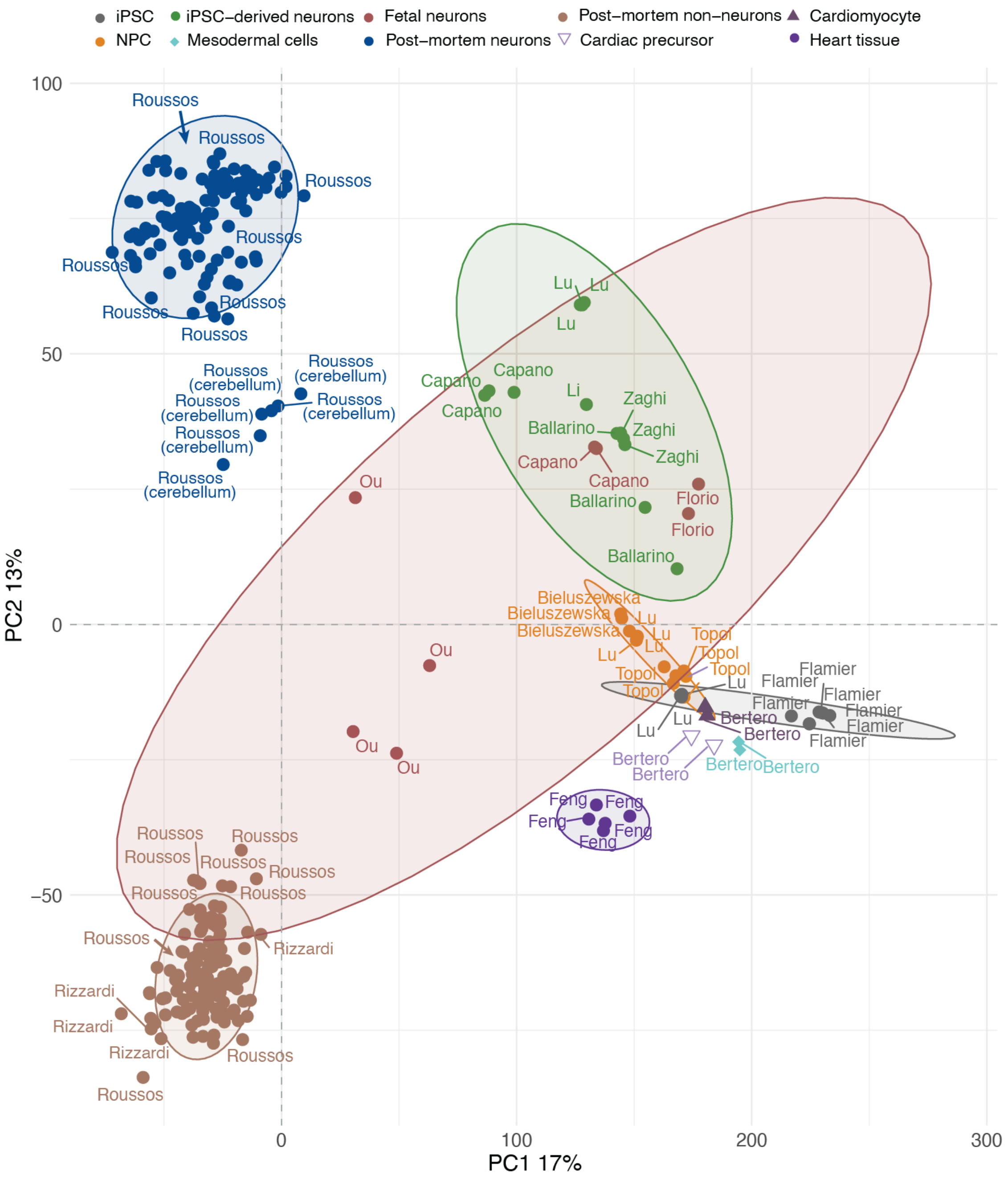


**Supplemental Fig. S10. PCA plots of gene expression changes in human samples.**

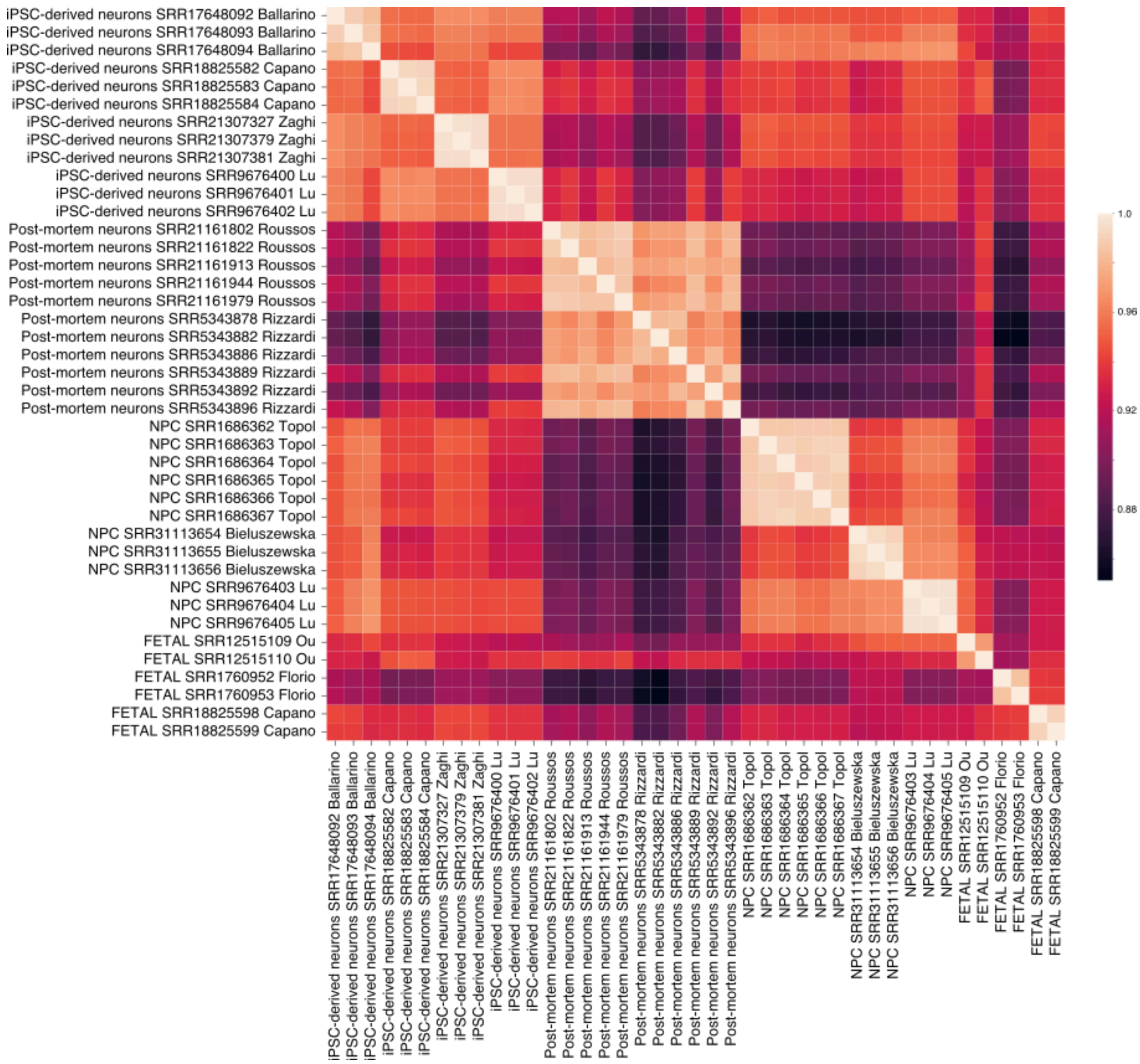


**Supplemental Fig. S11. Heatmap of Pearson's correlations between gene expression profiles across cell types and technical replicates.** Postmortem neurons are represented by prefrontal cortex samples, selected as a representative brain region and obtained from two independent datasets: Dong et al. (2024) and Rizzardi et al. (2019).

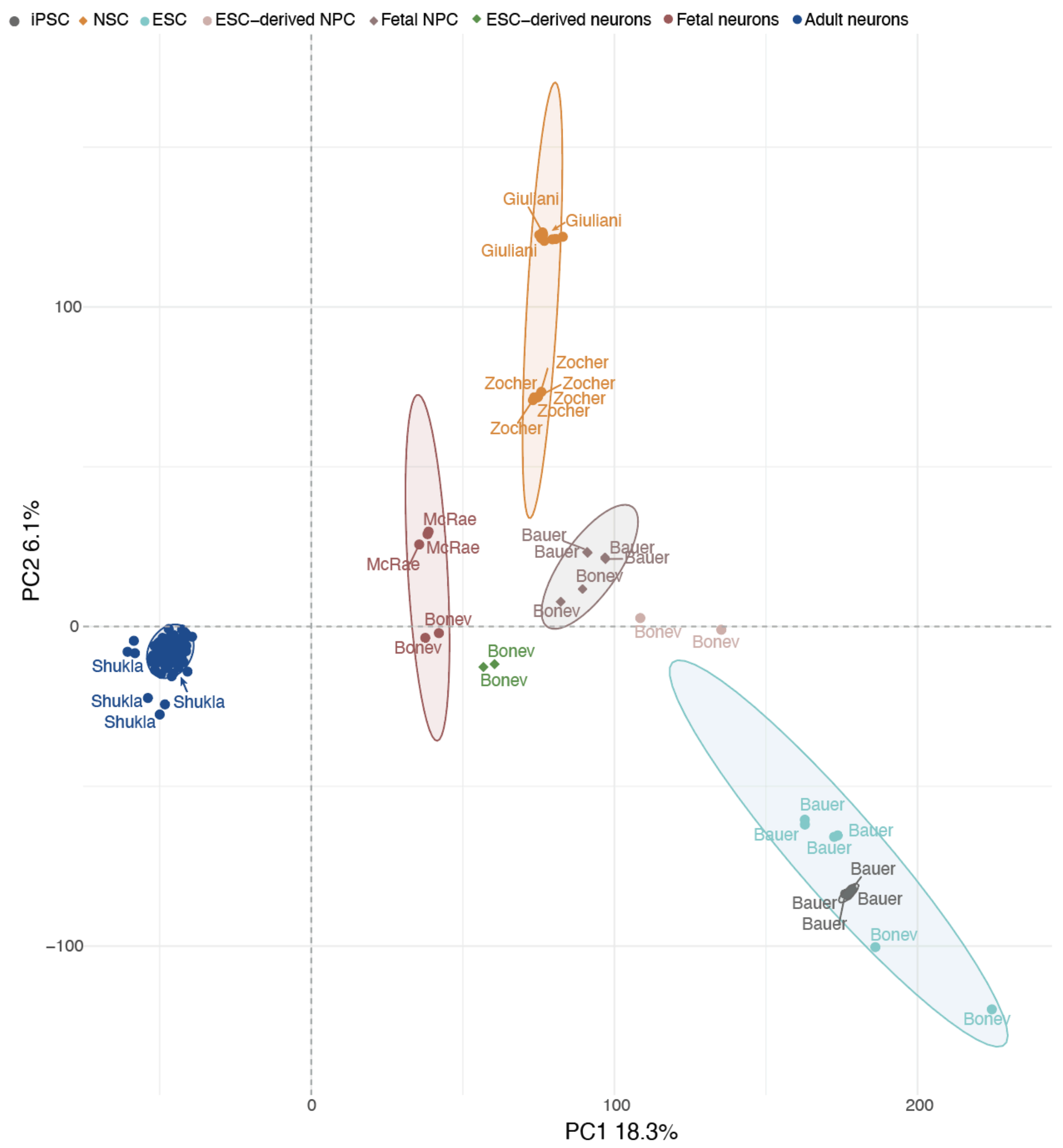


**Supplemental Fig. S12. PCA plots of gene expression changes in mouse samples.**

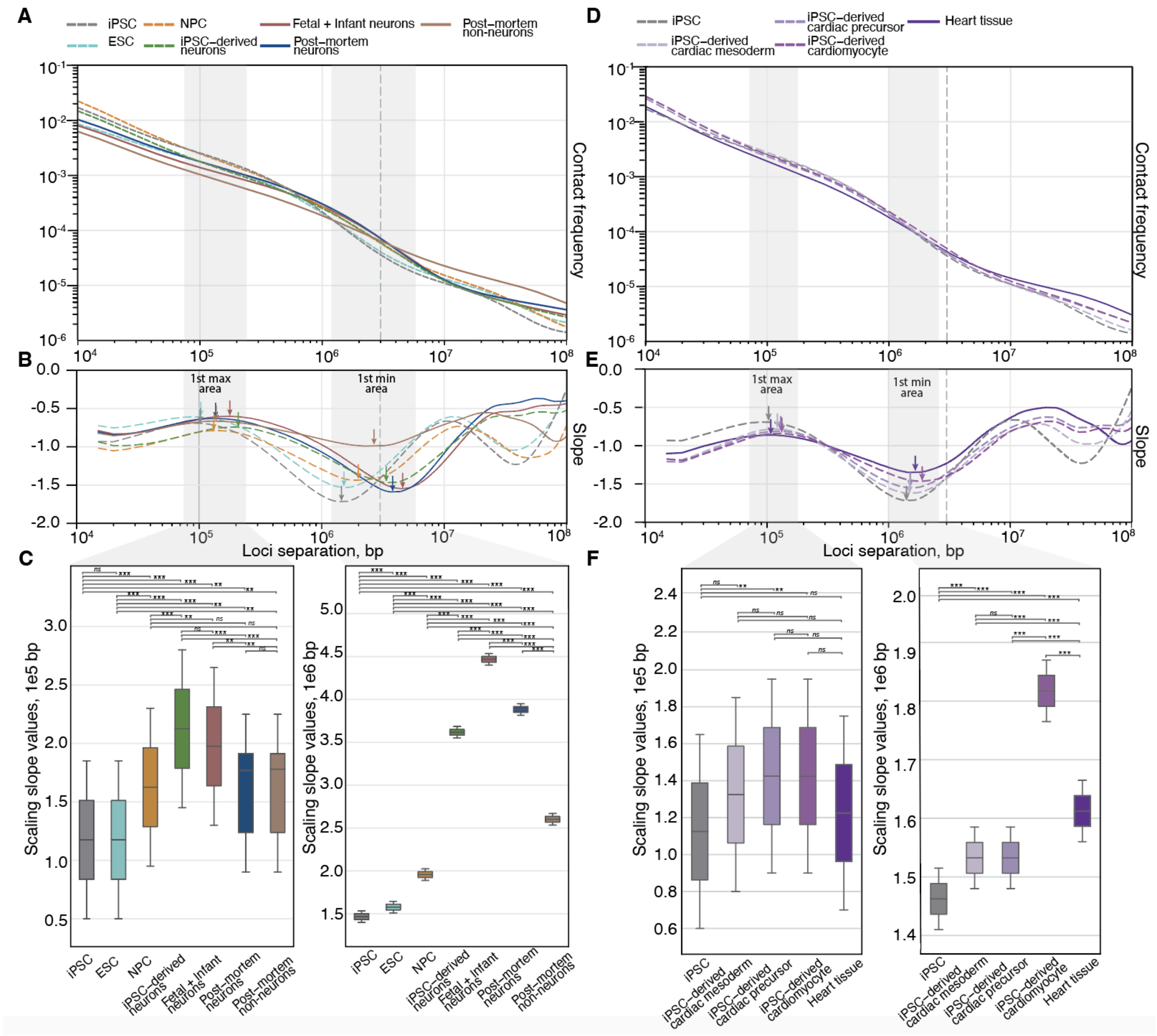


**Supplemental Fig. S13. Differential chromatin contact decay dynamics.** (A, D) Contact probability decay plot showing average interaction frequencies at various genomic distances for samples included in iPS-neuro and neuronal trajectories (A) and the cardiac differentiation trajectory (D). (B, E) Slopes of P(s) curves (depicting the relationship between chromatin contact probability and genomic distance) are shown for samples across neuronal and iPS-neuro differentiation trajectories (B) and the cardiac differentiation trajectory (E). (C, F) Box plots display the distribution of genomic distances for 10 bins upstream and 10 bins downstream (±10 bins) from the first slope maximum (left panels) and minimum (right panels) for samples across neuronal and iPS-neuro differentiation trajectories (C) and the cardiac differentiation trajectory (F). The position of the central bin (used as the reference for the ±10 bin window) varies between cell lines and is marked with a colored arrow matching each cell line; the region used to identify the minimum or maximum is indicated by a grey grey background in the figure. Asterisks indicate Mann-Whitney $U$ test $p$-values: ns - not significant change, ** - $p < 0.01$, *** - $p < 0.001$.

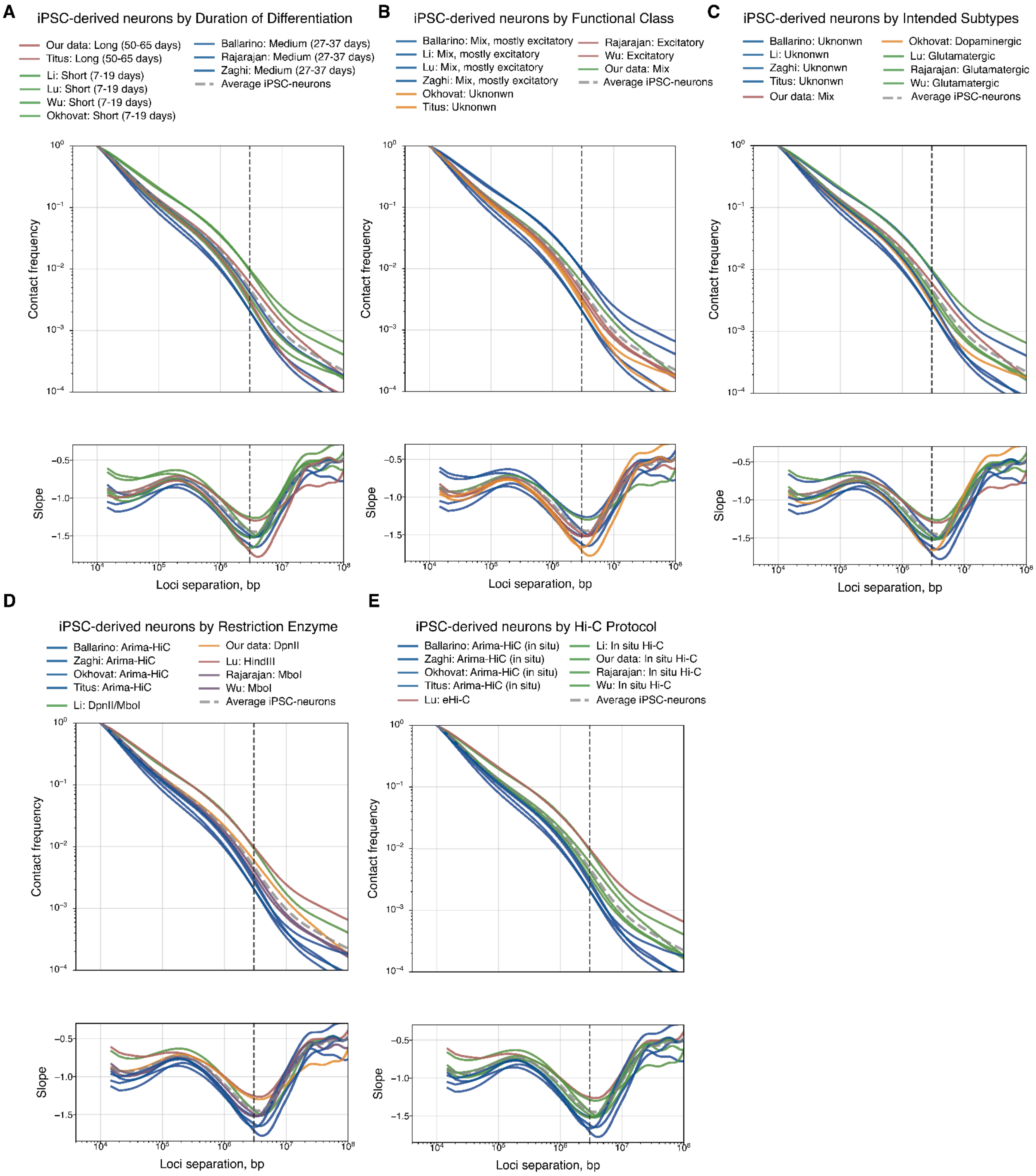


**Supplemental Fig. S14. Chromatin contact decay plots for iPSC-derived neurons.** Samples are grouped by different sample attributes: duration of differentiation protocol (A), neuronal functional class (B), intended neuronal subtype (C), Hi-C protocol (C), restriction enzyme used for the Hi-C library preparation (E). For each grouping, contact frequency (top) and slope (bottom) are plotted as a function of genomic distance. The dashed line indicates a separation of one megabase. The average across all iPSC-derived neuron samples is shown as the dashed grey line.

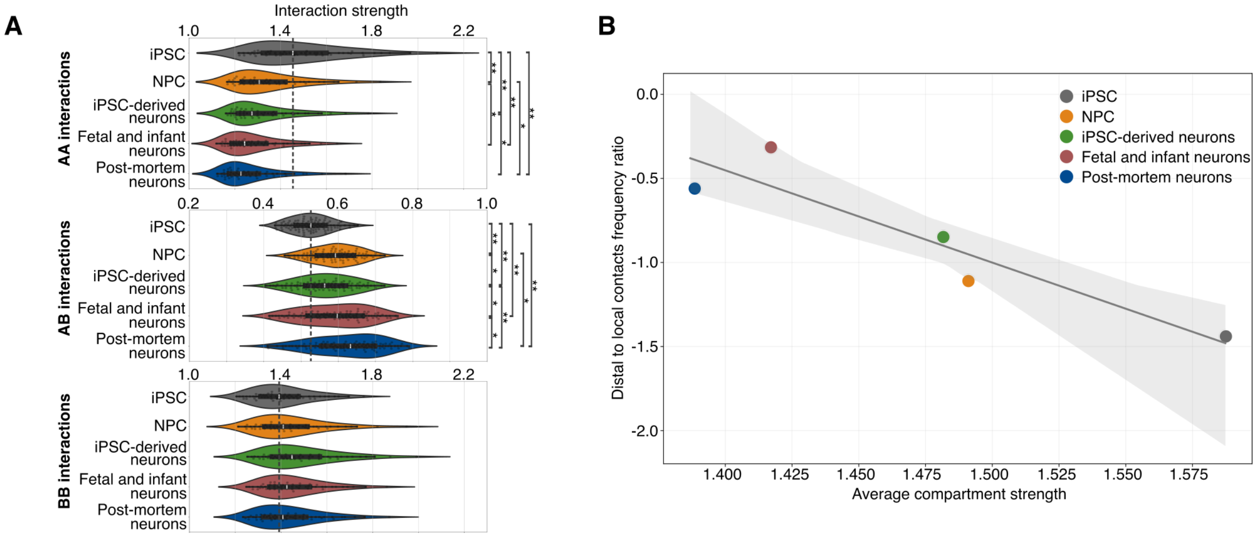


**Supplemental Fig. S15. Analysis of inter- and intracompartmental interactions.** (A) Intra- (AA, BB) and inter- (AB) compartmental interaction strength calculated within 5x5 corner areas of saddle plots for each cell group. Asterisks indicate BH-adjusted Mann-Whitney *U* test *p*-values: * - $p < 0.01$, ** - $p < 0.0001$. (B) Relationship between compartment strength and contact frequency. Linear regression of distal-to-local contact frequency ratio (DLR) on average compartment strength for selected cell groups. Shaded area - confidence interval = 95%.

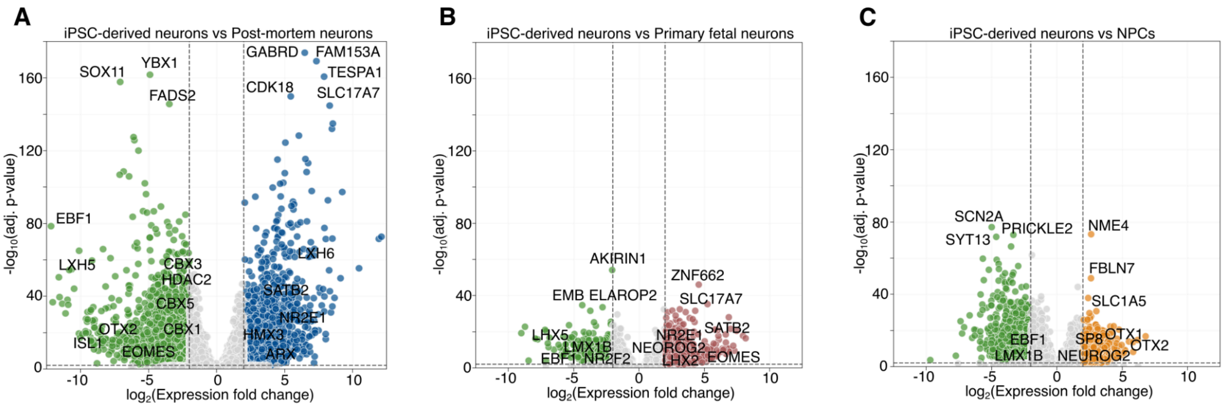


**Supplemental Fig. S16. Analysis of differentially expressed genes.** Volcano plots of differentially expressed genes (DEGs) for postmortem vs. iPSC-derived neurons (A), primary fetal vs. iPSC-derived neurons (B), and NPCs vs. iPSC-derived neurons (C). Colors indicate genes upregulated in iPSC-derived neurons (green), post-mortem neurons (blue), primary fetal neurons (red), or NPCs (orange).

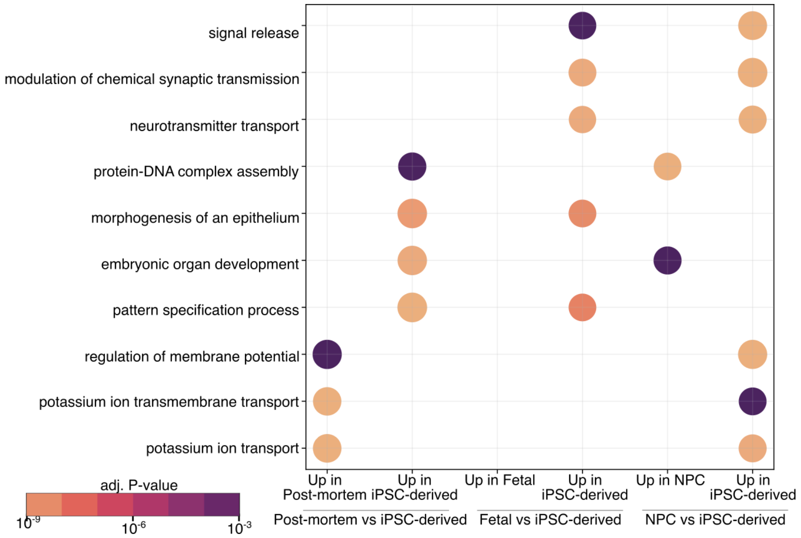


**Supplemental Fig. S17. Shared GO enrichment terms between all cell groups compared to iPSC-derived neurons.** Top of shared GO terms enrichment for differentially expressed genes specific for up- and downregulated genes in iPSC-derived neurons sorted by BH-adjusted *p*-value.

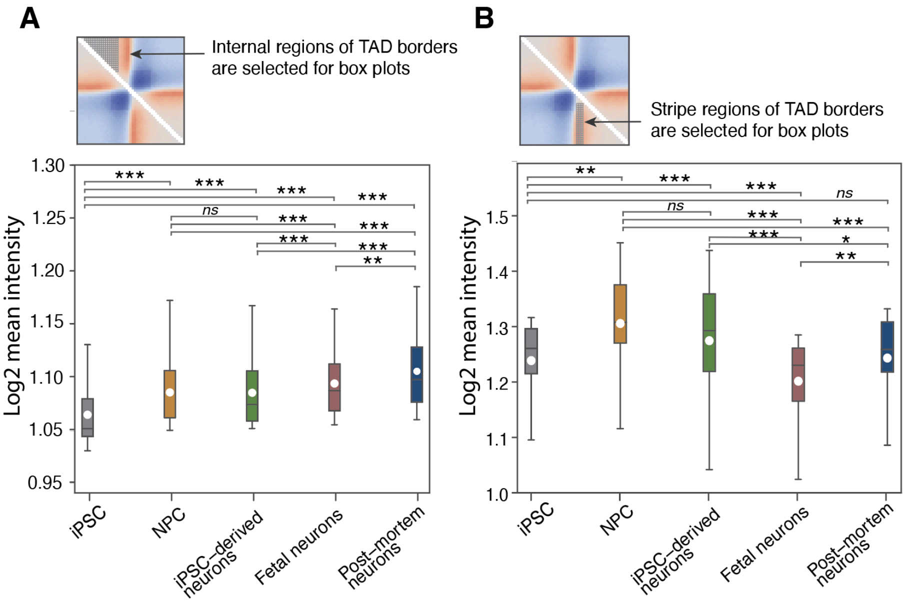


**Supplemental Fig. S18. TAD borders strength for the studied groups.** Box plots of TAD borders strength (observed/expected) in the internal regions of the borders (A) and stripe regions corresponding to the extrusion tracks (B). The values for the plot were selected from the corresponding regions noted on the average border figures (upper panel). Asterisks indicate Welch's *t*-test *p*-values: ns - not significant change,* -p < 0.05, ** -p < 0.01, *** - p < 0.001.

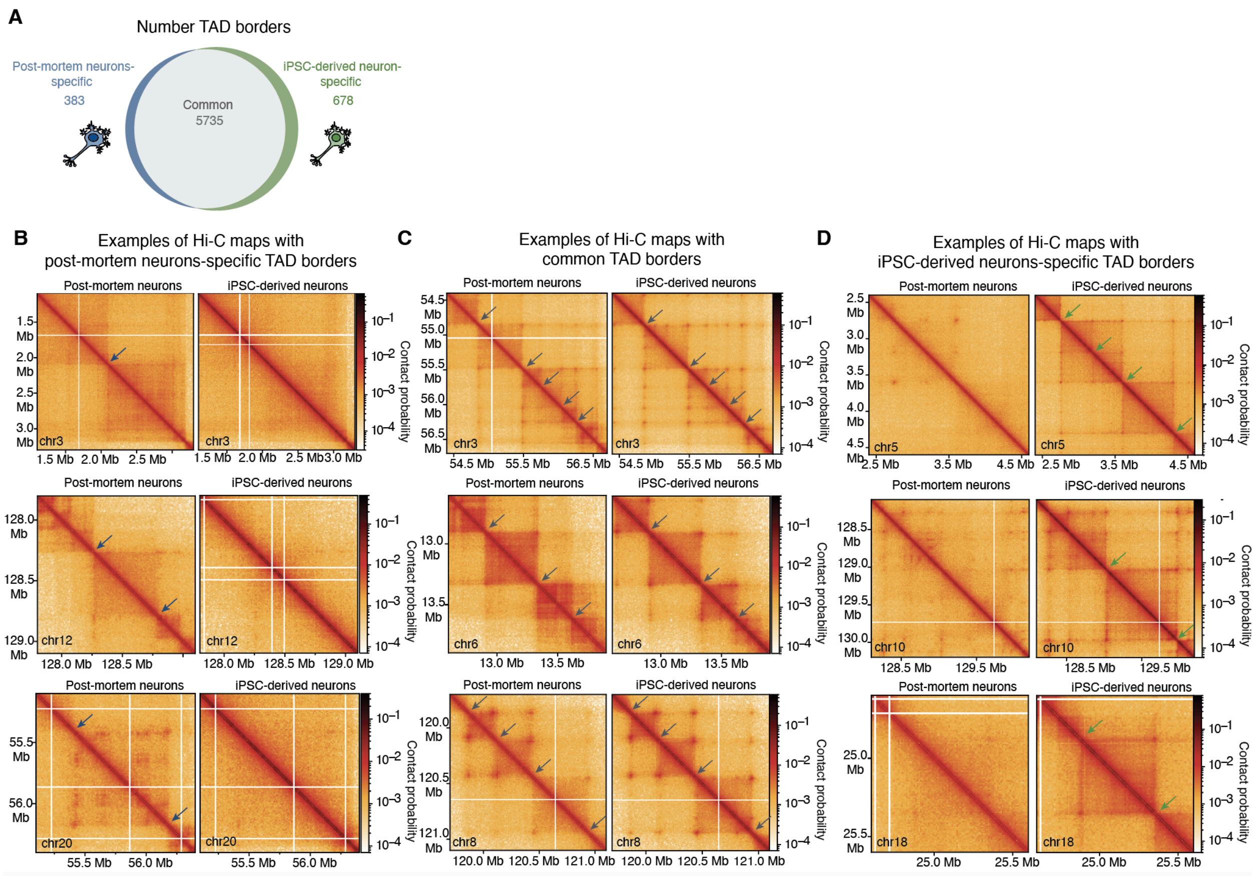


**Supplemental Fig. S19. Cell-type-specific TAD borders between postmortem and iPSC-derived neurons.** (A) Venn diagram showing the number of TAD borders that are cell-type-specific or shared between postmortem neurons and iPSC-derived neurons. (B-D) Representative Hi-C maps illustrating genomic regions with cell-type-specific and common TAD borders. Borders were classified as cell-type-specific based on significant differences in border strength identified by the Mann Whitney U test. Arrows indicate the location of cell-type-specific or common TAD borders, respectively.

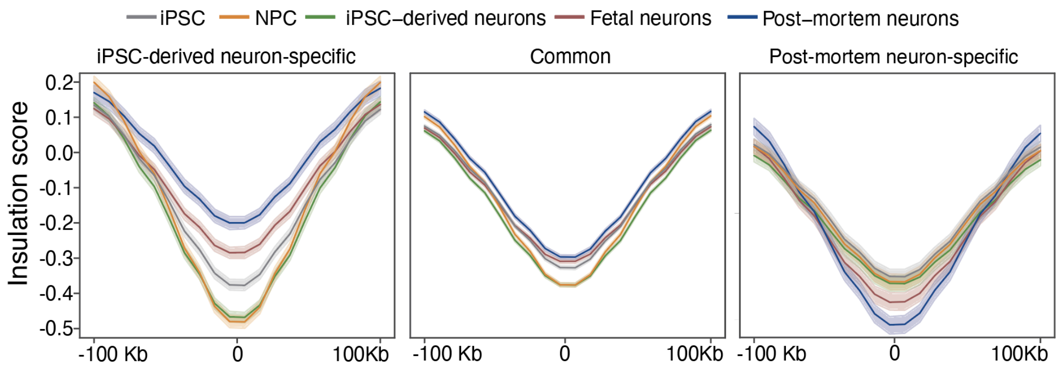


**Supplemental Fig. S20. Insulation score profiles at shared and cell type-specific TAD borders in iPSC-derived and postmortem neurons.** Insulation score profiles at TAD borders common to postmortem and iPSC-derived neurons (middle panel), and at differential borders: iPSC-derived neuron-specific (left panel) and postmortem neuron-specific (right panel).

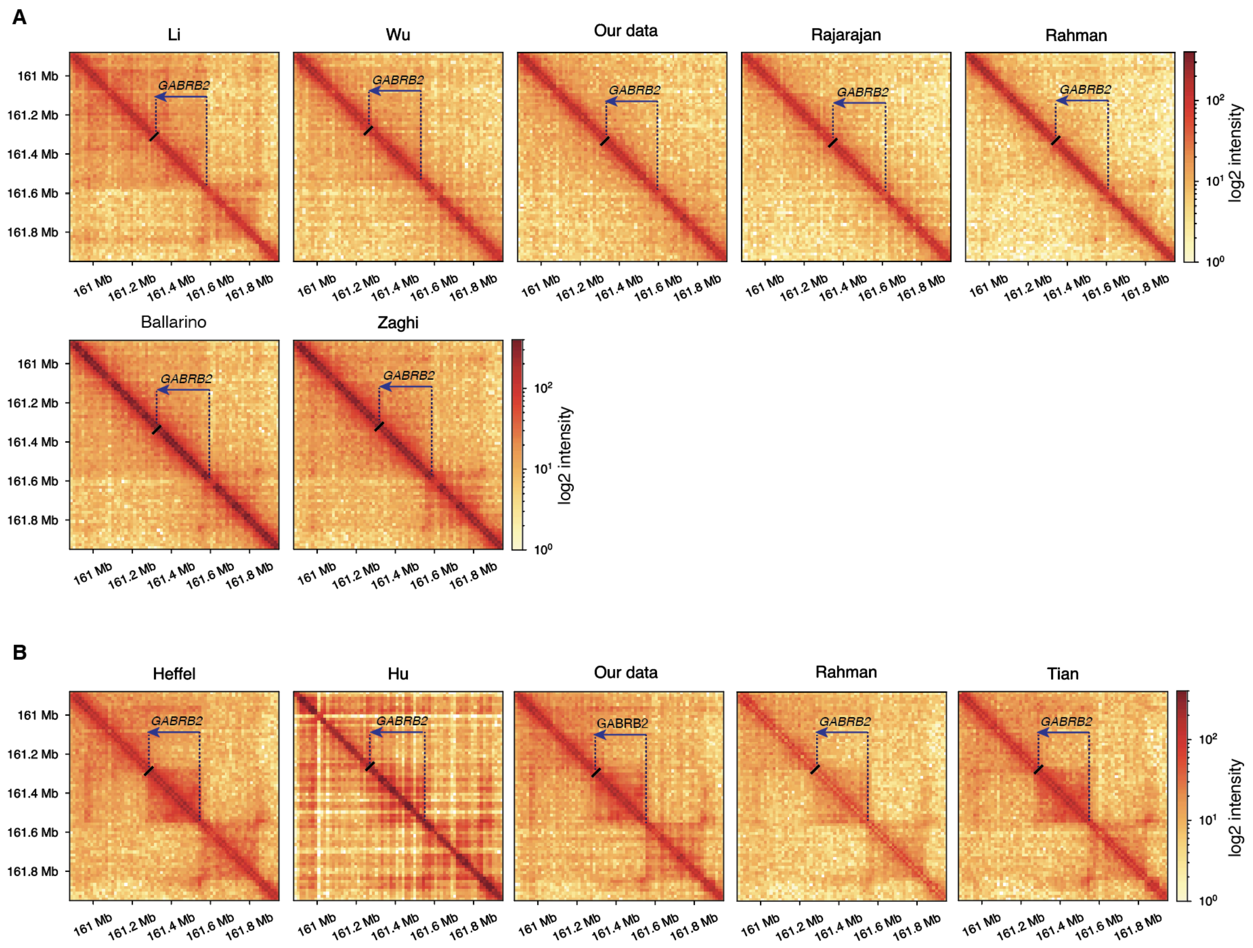


**Supplemental Fig. S21. Hi-C interaction maps surrounding the *GABRB2* locus in iPSC-derived and postmortem neurons.** Regions of the Hi-C maps around the *GABRB2* gene of iPSC-derived (A) and postmortem (B) neurons.

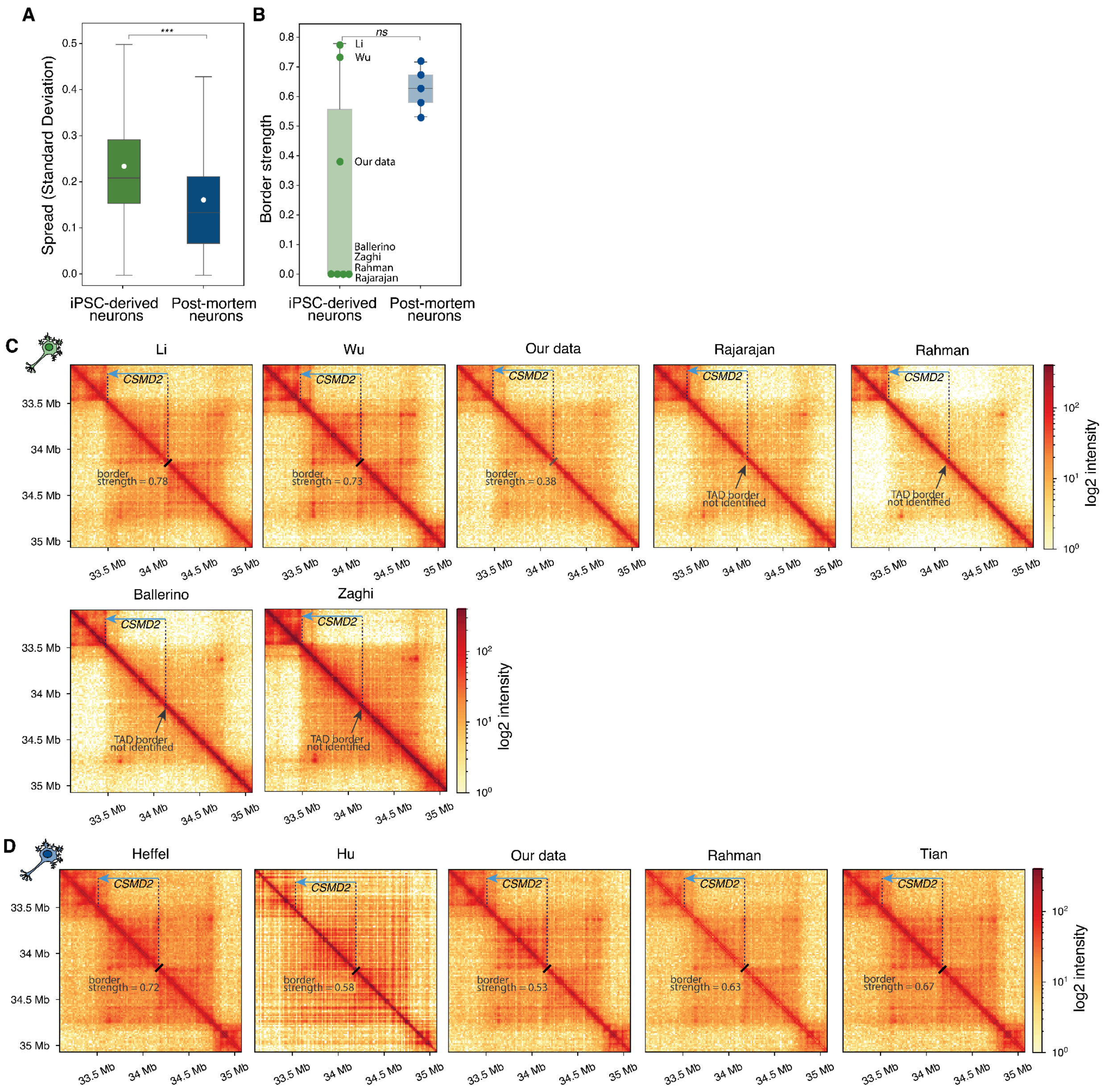


**Supplemental Fig. S22. Variability of TAD border strength in iPSC-derived and postmortem neurons.** (A) Box plot of standard deviation of the TAD borders strength for iPSC-derived and postmortem neurons across all identified TAD borders. Asterisks indicate Welch's *t*-test *p*-values: *** - $p < 0.001$. (B) Box plot of standard deviation of the strength of one selected TAD border located at *CSMD2* gene for studied samples. (C,D) Regions of Hi-C maps around the *CSMD2* gene for iPSC-derived (C) and postmortem (D) neurons. The position of the *CSMD2* gene is indicated by a blue arrow. Black lines mark TAD borders identified by the TAD calling algorithm. Absence of a border is indicated by grey arrows and corresponding annotations.

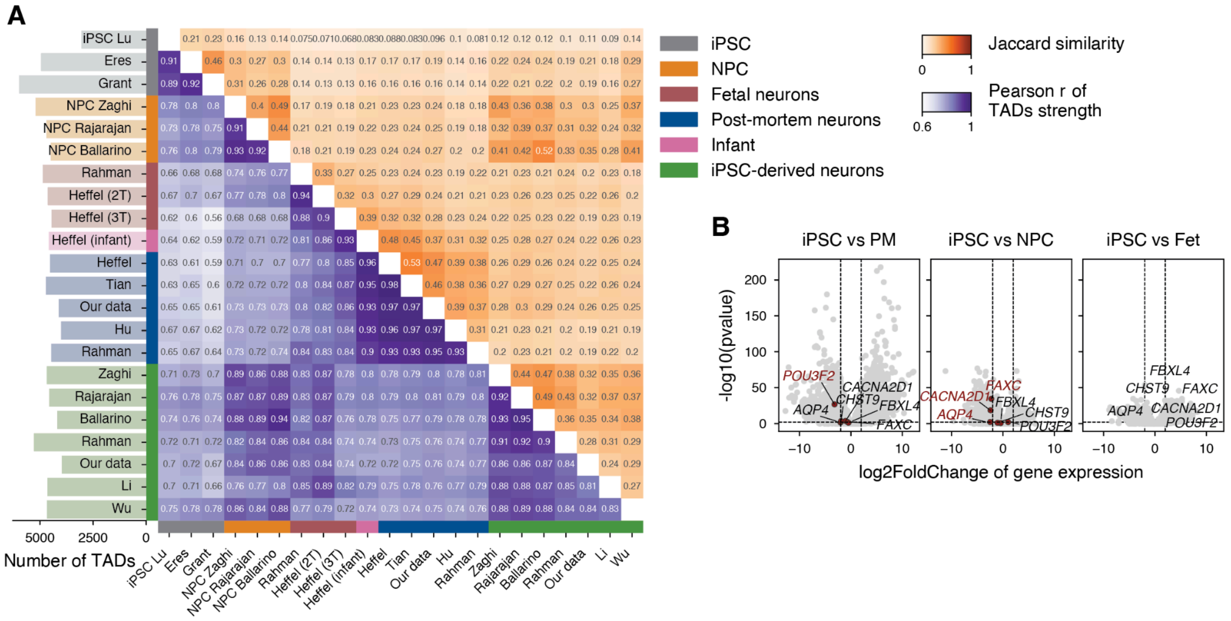


**Supplemental Fig. S23. Inner compactness of TADs.** (A) Heatmap showing pairwise comparisons of TAD organization across samples. The upper triangle represents Jaccard similarity of TADs with common borders, and the lower triangle shows Pearson's correlation of TAD strength. The adjacent bar plot indicates the total number of TADs identified in each map. (B) Volcano plot of differential gene expression between iPSC-derived neurons and other cell groups, highlighting example genes shown in Figure 3K.

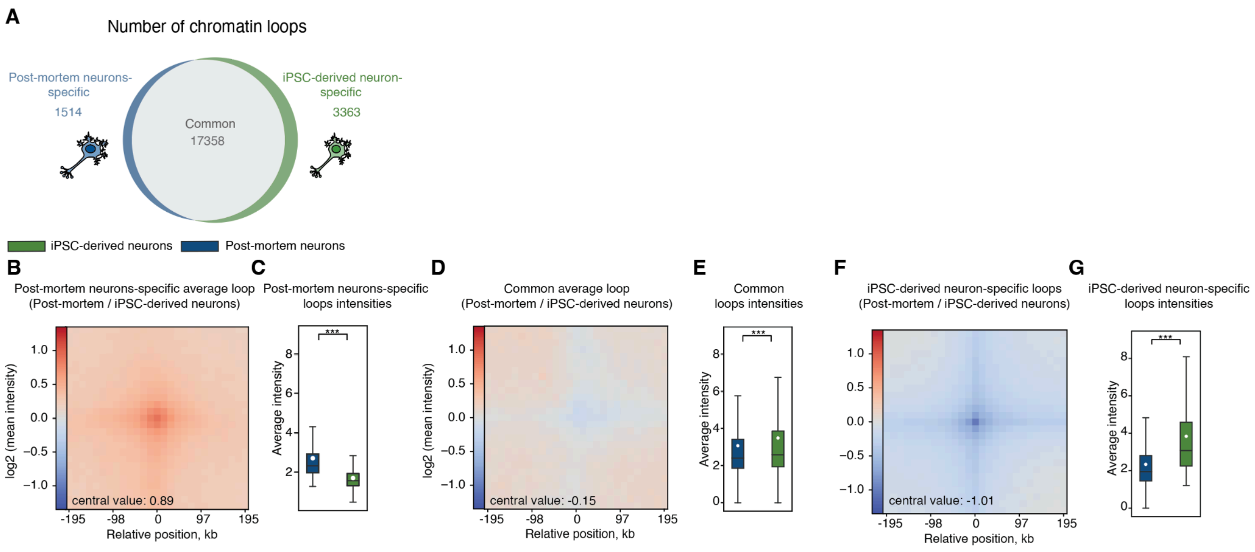


**Supplemental Fig. S24. Cell-type-specific chromatin loops between postmortem and iPSC-derived neurons.** (A) Venn diagram showing the number of chromatin loops that are cell-type-specific or shared between postmortem neurons and iPSC-derived neurons. (B, D, F) Average loop intensities (mean observed/expected) between postmortem and iPSC-derived neurons for (B) postmortem neuron-specific loops, (D) common loops, and (F) iPSC-derived neuron-specific loops. In each heatmap, the center represents the midpoint between the loop anchors, and the value at the center ("central value") is displayed. Color scale represents $\log_2$-transformed mean intensity ratio. (C, E, G) Box plots of average intensity for postmortem and iPSC-derived neurons for the corresponding loop sets shown in panels B, D, and F. Blue boxes represent postmortem neurons, and green boxes represent iPSC-derived neurons. White dots denote the mean values. Asterisks indicate *t*-test *p*-values: *** – $p < 0.001$.

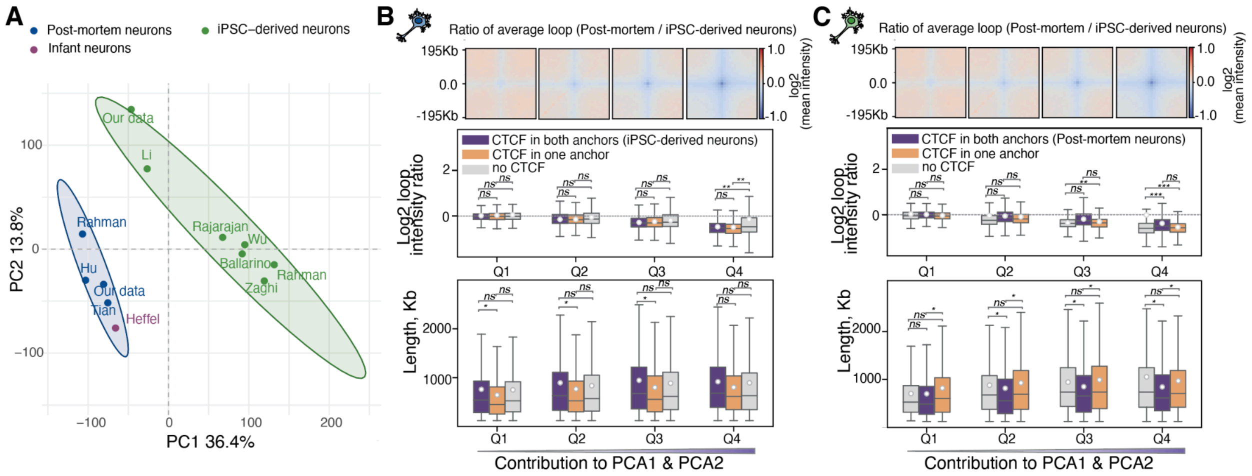


**Supplemental Fig. S25. Principal component analysis and categorization of chromatin loops in iPSC-derived and postmortem neurons.** (A) PCA of loop intensities for postmortem and iPSC-derived neurons. (B,C) Box plots categorizing loops based on their contribution to separation in PC1 and PC2 and the presence of CTCF based on layout for iPSC-derived (B) and postmortem (C) neurons; the upper plot shows the ratio of average loops, the middle panel details loop intensity ratios and the bottom plot displays loop lengths in kilobases. In all panels, asterisks indicate significance by two-sided *t*-test with FDR correction: **** – $p < 0.0001$, *** – $p < 0.001$, ** – $p < 0.01$, * – $p < 0.05$, *ns* – not significant.

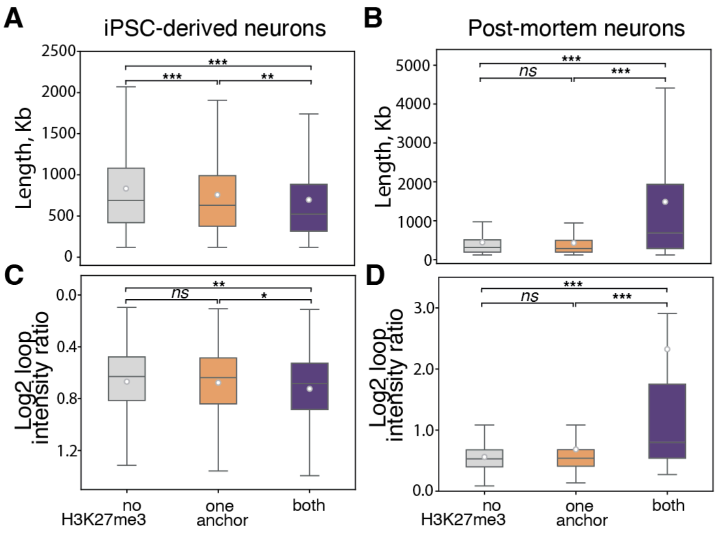


**Supplemental Fig. S26. Loops properties based on the intersection with H3K27me3 mark.** Box plot of loop length and logarithm of loop intensity ratio (iPSC-derived compared with postmortem neurons) for iPSC-derived (A,C) and postmortem (B,D) neurons. Asterisks indicate Welch's *t*-test *p*-values: *- $p < 0.05$, **- $p < 0.05$, *** - $p < 0.001$.

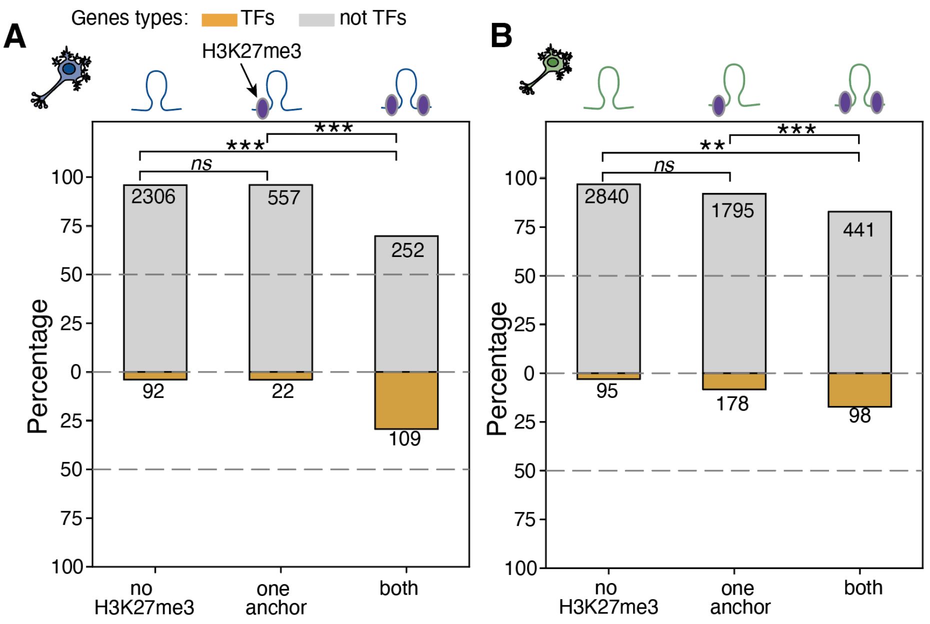


**Supplemental Fig. S27. Distribution of transcription factors (TFs) and non-TFs among genes located at chromatin loops with different H3K27me3 status at loop anchors.** Bar plots display the percentage of genes that are transcription factors (TFs, orange) or non-transcription factors (non-TFs, gray) for three groups of loops: with no H3K27me3, one anchor marked with H3K27me3, or both anchors marked with H3K27me3 for postmortem (A) and iPSC-derived (B) neurons. Numbers inside bars indicate total gene count per group (TFs in orange, non-TFs in gray). Asterisks denote Fisher's exact test *p*-values: ** – p < 0.01, * – p < 0.001, *ns* – not significant.

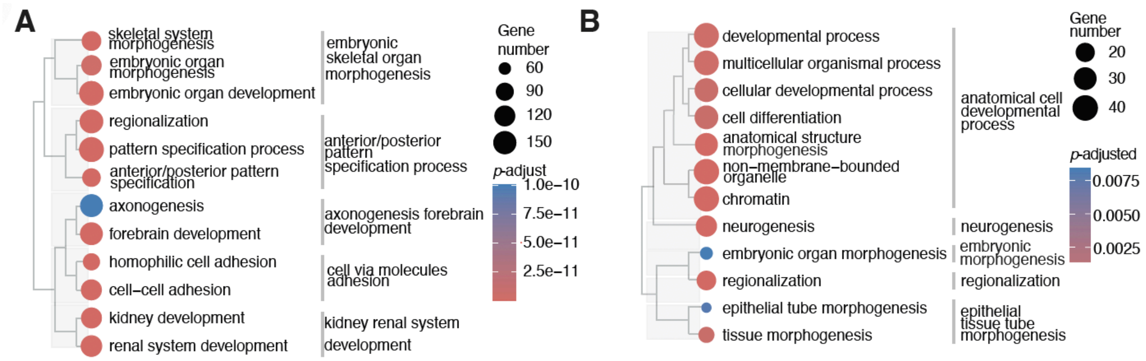


**Supplemental Fig. S28. GO term enrichment for transcription factors and genes located within H3K27me3-enriched chromatin loops.** (A) GO term enrichment analysis for downregulated TFs within H3K27me3-enriched chromatin loops in neurons. (B) GO term enrichment analysis for upregulated iPSC-derived genes located within chromatin loops detected in iPSC-derived neurons.

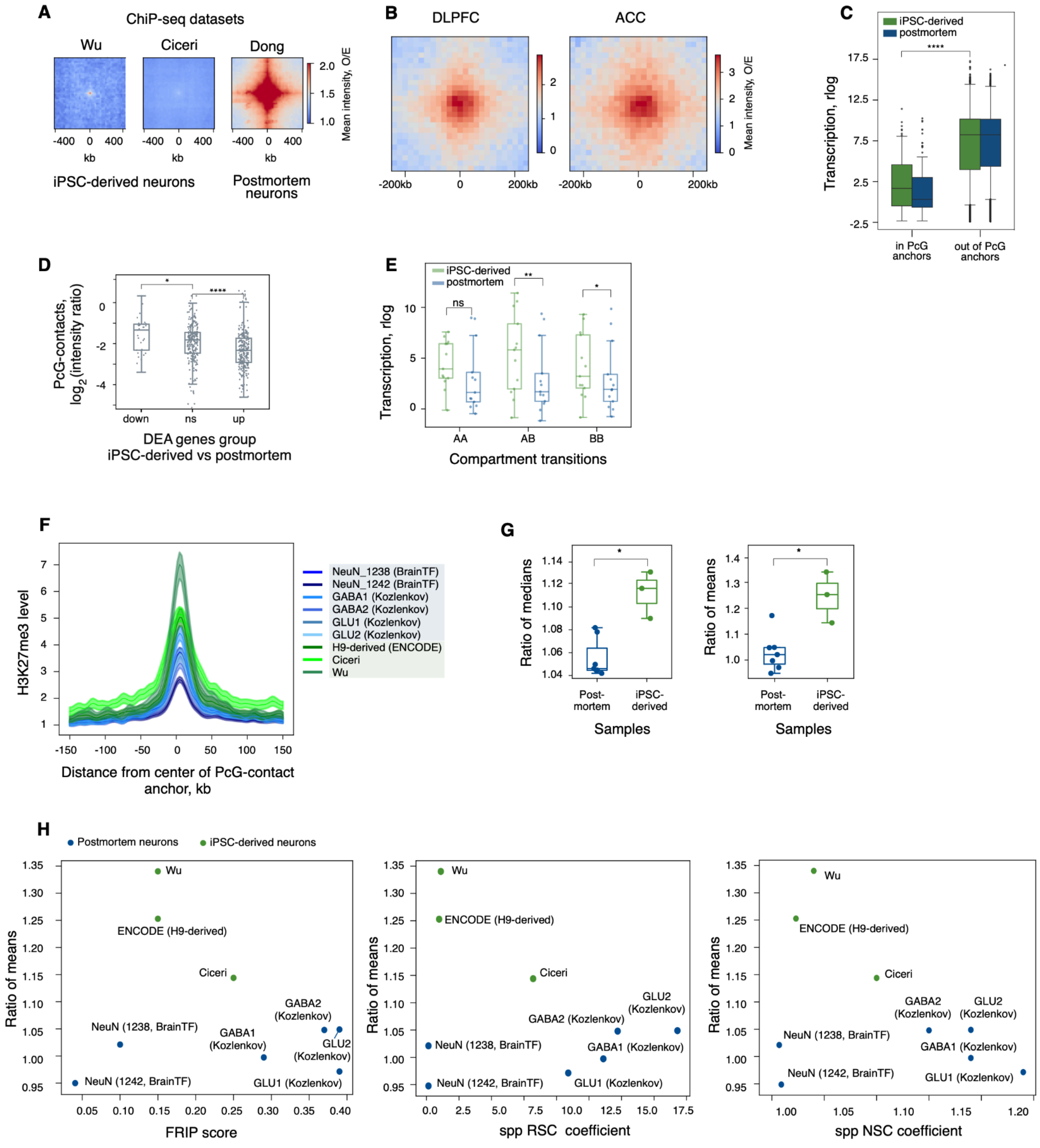

**Supplemental Fig. S29. Analysis of long-range PcG-contacts.** (A) Average contact frequencies between genomic bins containing broad H3K27me3 ChIP-seq peaks and separated by at least 3 Mb. For the left and middle heatmaps, peaks from Wu et al. (2021b) and Ciceri et al. (2024), respectively, were used for all iPSC-derived samples. The right heatmap corresponds to postmortem neurons using H3K27me3 peaks from Dong et al. (2022) (B) Average PcG-contact intensities for postmortem neurons from dorsolateral prefrontal cortex (Hu et al. (2021)) and anterior cingulate cortex (synapse.org: syn15137926). (C) Gene expression within and outside anchors of PcG-contacts. (D) The logarithmic ratio of PcG-contact intensities, comparing iPSC-derived to postmortem neurons, categorized by the presence of genes from DEA groups within anchors. Asterics indicate Mann-Whitney U-test *p*-value: * - *p*-value < 0.05, **** - *p*-value < 0.0001 (0.03 for comparison of intensity ratios (iPSC-derived/postmortem) of PcG-contacts containing down-regulated vs non-differential genes within anchors and $2.06*10^{-8}$ for comparison of intensity ratios (iPSC-derived/postmortem) of PcG-contacts containing non-differential vs upregulated genes within anchors). (E) Expression levels for random samples of protein-coding genes located at PcG-contact anchors, stratified by compartment transition categories. Expression distributions and sample sizes for A→A and B→B transitions were matched to those of the A→B transition group. (F). Average per-sample profile of H3K27me3 ChIP-seq, centered on PcG-contact anchor midpoints, without bringing to the common baseline. (G) Ratio of median (left) and mean (right) MACS3-defined ChIP-seq peaks fold-enrichment calculated within PcG-contact anchors to this median/mean value calculated outside PcG-contacts. Asterisks indicate Mann-Whitney U-test *p*-value: * - *p*-value < 0.05 (0.016 and 0.033 for left and right plots respectively). (H) Relationship between ratio of mean MACS3-defined ChIP-seq peaks fold-enrichment calculated within PcG-contact anchors to mean value calculated outside PcG-contacts and metrics of ChIP-seq quality (FRIP score, spp RSC coefficient, spp NSC coefficient – left, mid, right respectively).

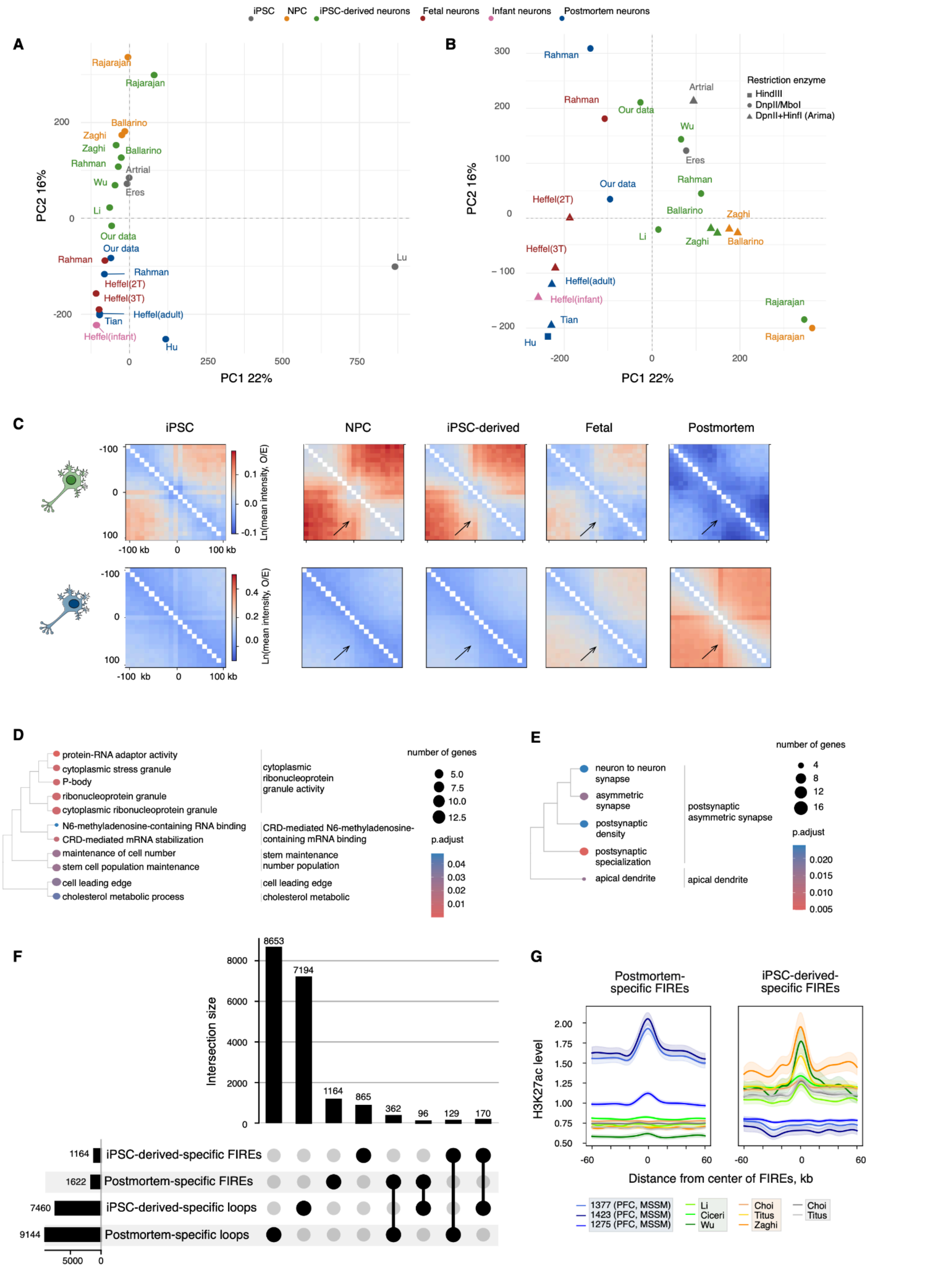


**Supplemental Fig. S30. Analysis of FIREs.** (A) PCA of FIREs scores across the analyzed cell groups, including a sample generated using the easy Hi-C protocol from Lu et al. (2020). (B) PCA of FIREs scores across the analyzed cell groups. Shapes of points indicate the restriction enzyme used in the Hi-C protocol. (C) Average intensities of iPSC-derived-specific (top) and postmortem-specific (bottom) FIREs in each cell group. (D) GO terms enrichment for genes located in postmortem-specific FIREs. (E) GO terms enrichment for genes located in iPSC-derived-specific FIREs. (F) UpSet plot of intersections between cell type-specific loops and FIREs positions. (G) Average per-sample profile of H3K27ac ChIP-seq, centered on postmortem-specific (left) and iPSC-derived-specific (right) FIREs without bringing to common baseline.

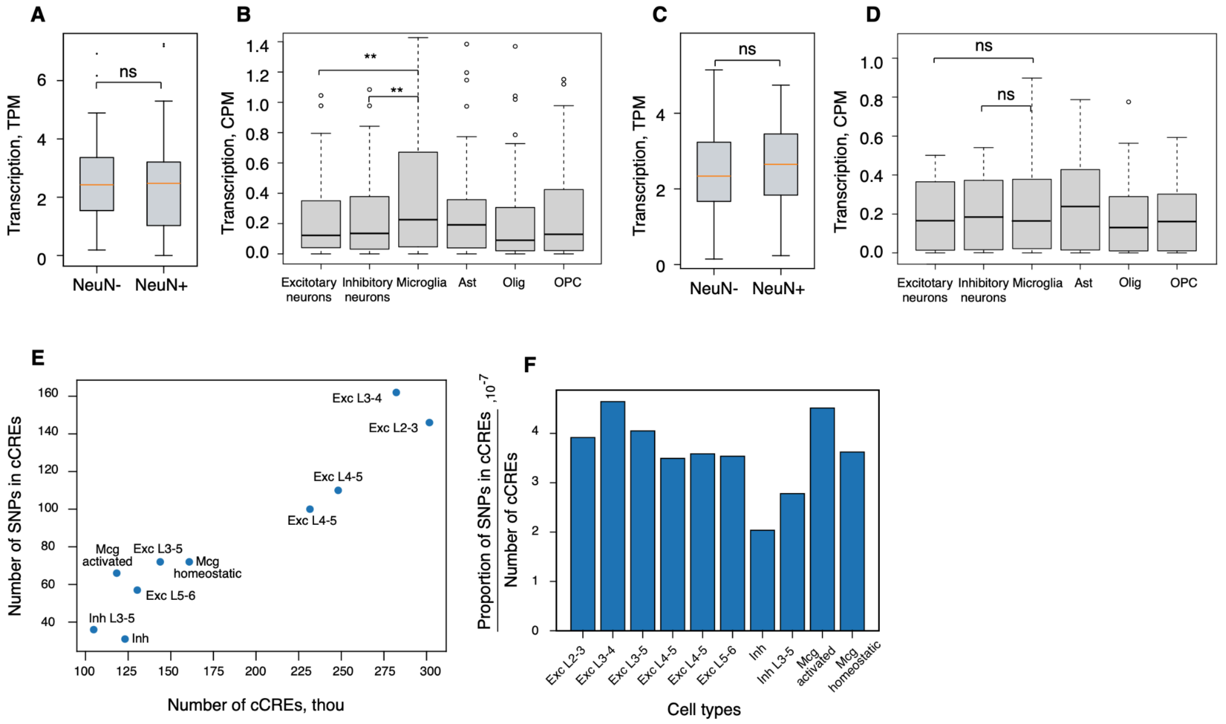


**Supplemental Fig. S31. Activity of GWAS AD variants in different cell types.** A. Expression of genes containing AD-associated SNPs in sorted NeuN- and NeuN+ nuclei from Rizzardi et al. (2019). B. Expression of genes containing AD-associated SNPs in snRNA-seq data from Jeffries et al. (2025). C. Expression of genes interacting with bins containing AD-associated SNPs in sorted NeuN- and NeuN+ nuclei from Rizzardi et al. (2019). D. Expression of genes interacting with bins containing AD-associated SNPs in snRNA-seq data from Jeffries et al. (2025). E. Scatter plot showing numbers of AD-associated SNPs located in cCREs of different cell types vs numbers of cCREs. F. Proportion of AD-associated SNPs located in cCREs, normalized by the number of cCREs across cell types.

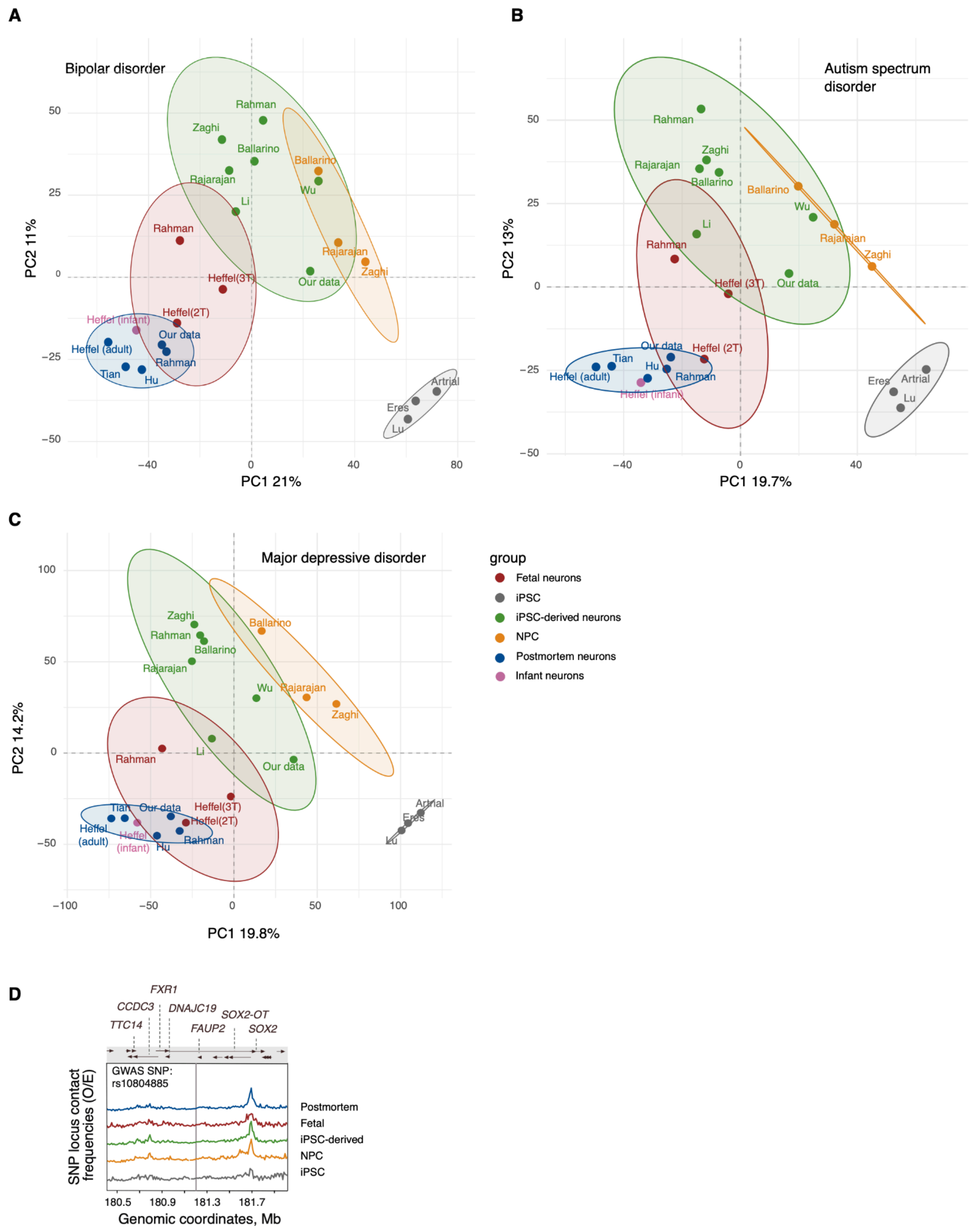


**Supplemental Fig. S32. Analysis of contacts involving disease-associated SNPs.** (A–C) PCA of intensities of significant contacts involving SNPs associated with bipolar disorder, autism spectrum disorder and major depressive disorder. (D) Average observed/expected contact frequency profiles for schizophrenia-associated GWAS loci interacting with *SOX2* genes.

### Supplemental Tables

**Supplemental Table S1. Bulk Hi-C data for human and mouse.**
(provided as a separate .xlsx file)

**Supplemental Table S2. Differentially expressed genes between iPSC-derived neurons and NPCs, fetal neurons and postmortem neurons.**
(provided as a separate .xlsx file)

**Supplemental Table S3. Hi-C sample metadata.**
(provided as a separate .xlsx file)

**Supplemental Table S4. Sn-m3C data for human and mouse.**
(provided as a separate .xlsx file)

**Supplemental Table S5. Number of merged samples and datasets for each cell type.**
(provided as a separate .xlsx file)

**Supplemental Table S6. Bulk RNA-seq data for human and mouse.**
(provided as a separate .xlsx file)

**Supplemental Table S7. ChIP-seq metadata.**
(provided as a separate .xlsx file)

### Supplemental References


Abdennur N, Mirny LA. 2020. Cooler: scalable storage for Hi-C data and other genomically labeled arrays. *Bioinformatics* **36**: 311–316.

Adams MJ, Streit F, Meng X, Awasthi S, Adey BN, Choi KW, Chundru VK, Coleman JRI, Ferwerda B, Foo JC, et al. 2025. Trans-ancestry genome-wide study of depression identifies 697 associations implicating cell types and pharmacotherapies. *Cell* **188**: 640-652.e9.

Andrews S. 2010. FastQC A Quality Control tool for High Throughput Sequence Data. https://www.bioinformatics.babraham.ac.uk/projects/fastqc/ (Accessed March 15, 2026).

Ballarino R, Bouwman BAM, Agostini F, Harbers L, Diekmann C, Wernersson E, Bienko M, Crosetto N. 2022. An atlas of endogenous DNA double-strand breaks arising during human neural cell fate determination. *Sci Data* **9**: 400.

Bolger AM, Lohse M, Usadel B. 2014. Trimmomatic: a flexible trimmer for Illumina sequence data. *Bioinformatics* **30**: 2114–2120.

Chen B, Khodadoust MS, Liu CL, Newman AM, Alizadeh AA. 2018. Profiling Tumor Infiltrating Immune Cells with CIBERSORT. In *Cancer Systems Biology* (ed. L. Von Stechow), Vol. 1711 of *Methods in Molecular Biology*, pp. 243–259, Springer New York, New York, NY http://link.springer.com/10.1007/978-1-4939-7493-1_12 (Accessed March 16, 2026).

Ciceri G, Baggiolini A, Cho HS, Kshirsagar M, Benito-Kwiecinski S, Walsh RM, Aromolaran KA, Gonzalez-Hernandez AJ, Munguba H, Koo SY, et al. 2024. An epigenetic barrier sets the timing of human neuronal maturation. *Nature* **626**: 881–890.

Crowley C, Yang Y, Qiu Y, Hu B, Abnousi A, Lipiński J, Plewczyński D, Wu D, Won H, Ren B, et al. 2021. FIREcaller: Detecting frequently interacting regions from Hi-C data. *Comput Struct Biotechnol J* **19**: 355–362.

Dale RK, Pedersen BS, Quinlan AR. 2011. Pybedtools: a flexible Python library for manipulating genomic datasets and annotations. *Bioinformatics* **27**: 3423–3424.

Darmanis S, Sloan SA, Zhang Y, Enge M, Caneda C, Shuer LM, Hayden Gephart MG, Barres BA, Quake SR. 2015. A survey of human brain transcriptome diversity at the single cell level. *Proc Natl Acad Sci* **112**: 7285–7290.

Dong P, Hoffman GE, Apontes P, Bendl J, Rahman S, Fernando MB, Zeng B, Vicari JM, Zhang W, Girdhar K, et al. 2022. Population-level variation in enhancer expression identifies disease mechanisms in the human brain. *Nat Genet* **54**: 1493–1503.

Dong P, Song L, Bendl J, Misir R, Shao Z, Edelstien J, Davis DA, Haroutunian V, Scott WK, Acker S, et al. 2024. A multi-regional human brain atlas of chromatin accessibility and gene expression facilitates promoter-isoform resolution genetic fine-mapping. *Nat Commun* **15**: 10113.

Fishilevich S, Nudel R, Rappaport N, Hadar R, Plaschkes I, Iny Stein T, Rosen N, Kohn A, Twik M, Safran M, et al. 2017. GeneHancer: genome-wide integration of enhancers and target genes in GeneCards. *Database J Biol Databases Curation* **2017**: bax028.

Flyamer IM, Illingworth RS, Bickmore WA. 2020. Coolpup.py: versatile pile-up analysis of Hi-C data. *Bioinformatics* **36**: 2980–2985.

Frankish A, Diekhans M, Jungreis I, Lagarde J, Loveland JE, Mudge JM, Sisu C, Wright JC, Armstrong J, Barnes I, et al. 2021. GENCODE 2021. *Nucleic Acids Res* **49**: D916–D923.

Grove J, Ripke S, Als TD, Mattheisen M, Walters RK, Won H, Pallesen J, Agerbo E, Andreassen OA, Anney R, et al. 2019. Identification of common genetic risk variants for autism spectrum disorder. *Nat Genet* **51**: 431–444.

Heffel MG, Zhou J, Zhang Y, Lee D-S, Hou K, Pastor-Alonso O, Abuhanna KD, Galasso J, Kern C, Tai C-Y, et al. 2024. Temporally distinct 3D multi-omic dynamics in the developing human brain. *Nature* **635**: 481–489.

Hodge RD, Bakken TE, Miller JA, Smith KA, Barkan ER, Graybuck LT, Close JL, Long B, Johansen N, Penn O, et al. 2019. Conserved cell types with divergent features in human versus mouse cortex. *Nature* **573**: 61–68.

Holmqvist S, Lehtonen Š, Chumarina M, Puttonen KA, Azevedo C, Lebedeva O, Ruponen M, Oksanen M, Djelloul M, Collin A, et al. 2016. Creation of a library of induced pluripotent stem cells from Parkinsonian patients. *Npj Park Dis* **2**: 16009.

Hu B, Won H, Mah W, Park RB, Kassim B, Spiess K, Kozlenkov A, Crowley CA, Pochareddy S, Li Y, et al. 2021. Neuronal and glial 3D chromatin architecture informs the cellular etiology of brain disorders. *Nat Commun* **12**: 3968.

Imakaev M, Fudenberg G, McCord RP, Naumova N, Goloborodko A, Lajoie BR, Dekker J, Mirny LA. 2012. Iterative correction of Hi-C data reveals hallmarks of chromosome organization. *Nat Methods* **9**: 999–1003.

Jeffries AM, Yu T, Ziegenfuss JS, Tolles AK, Baer CE, Sotelo CB, Kim Y, Weng Z, Lodato MA. 2025. Single-cell transcriptomic and genomic changes in the ageing human brain. *Nature* **646**: 657–666.

Kozlenkov A, Li J, Apontes P, Hurd YL, Byne WM, Koonin EV, Wegner M, Mukamel EA, Dracheva S. 2018. A unique role for DNA (hydroxy)methylation in epigenetic regulation of human inhibitory neurons. *Sci Adv* **4**: eaau6190.

Li S. 2016. Regulatory genomic profiling in pure iPSC-derived human cortical neurons. *GEO*.

Lieberman-Aiden E, van Berkum NL, Williams L, Imakaev M, Ragoczy T, Telling A, Amit I, Lajoie BR, Sabo PJ, Dorschner MO, et al. 2009. Comprehensive mapping of long-range interactions reveals folding principles of the human genome. *Science* **326**: 289–293.

Love MI, Huber W, Anders S. 2014. Moderated estimation of fold change and dispersion for RNA-seq data with DESeq2. *Genome Biol* **15**: 550.

Lu L, Liu X, Huang W-K, Giusti-Rodríguez P, Cui J, Zhang S, Xu W, Wen Z, Ma S, Rosen JD, et al. 2020. Robust Hi-C maps of enhancer-promoter interactions reveal the function of non-coding genome in neural development and diseases. *Mol Cell* **79**: 521-534.e15.

McKenzie AT, Wang M, Hauberg ME, Fullard JF, Kozlenkov A, Keenan A, Hurd YL, Dracheva S, Casaccia P, Roussos P, et al. 2018. Brain Cell Type Specific Gene Expression and Co-expression Network Architectures. *Sci Rep* **8**: 8868.

Mesecar ME, Duffy MF, Acri DJ, Ding J, Langston RG, Shah SI, Nalls MA, Reed X, Scholz SW, Whitaker DT, et al. 2023. Region-Specific Transcriptional Signatures of Brain Aging in the Absence of Neuropathology at the Single-cell Level. http://biorxiv.org/lookup/doi/10.1101/2023.07.31.551097 (Accessed March 16, 2026).

Nekrasov ED, Vigont VA, Klyushnikov SA, Lebedeva OS, Vassina EM, Bogomazova AN, Chestkov IV, Semashko TA, Kiseleva E, Suldina LA, et al. 2016. Manifestation of Huntington's disease pathology in human induced pluripotent stem cell-derived neurons. *Mol Neurodegener* **11**: 27.

O'Connell KS, Koromina M, van der Veen T, Boltz T, David FS, Yang JMK, Lin K-H, Wang X, Coleman JRI, Mitchell BL, et al. 2025. Genomics yields biological and phenotypic insights into bipolar disorder. *Nature* **639**: 968–975.

Open2C, Abdennur N, Abraham S, Fudenberg G, Flyamer IM, Galitsyna AA, Goloborodko A, Imakaev M, Oksuz BA, Venev SV. 2022. Cooltools: enabling high-resolution Hi-C analysis in Python. 2022.10.31.514564. https://www.biorxiv.org/content/10.1101/2022.10.31.514564v1 (Accessed February 24, 2026).

Open2C, Abdennur N, Fudenberg G, Flyamer IM, Galitsyna AA, Goloborodko A, Imakaev M, Venev SV. 2024. Pairtools: From sequencing data to chromosome contacts. *PLoS Comput Biol* **20**: e1012164.

Pardiñas AF, Holmans P, Pocklington AJ, Escott-Price V, Ripke S, Carrera N, Legge SE, Bishop S, Cameron D, Hamshere ML, et al. 2018. Common schizophrenia alleles are enriched in mutation-intolerant genes and in regions under strong background selection. *Nat Genet* **50**: 381–389.

Patel H, Espinosa-Carrasco J, Wang C, Ewels P, bot nf-core, Silva TC, Peltzer A, Langer B, Guinchard S, Garcia MU, et al. 2024. nf-core/chipseq: nf-core/chipseq v2.1.0 - Platinum Willow Sparrow. https://zenodo.org/records/13899404 (Accessed February 24, 2026).

Patro R, Duggal G, Love MI, Irizarry RA, Kingsford C. 2017. Salmon provides fast and bias-aware quantification of transcript expression. *Nat Methods* **14**: 417–419.

Pedregosa F, Varoquaux G, Gramfort A, Michel V, Thirion B, Grisel O, Blondel M, Müller A, Nothman J, Louppe G, et al. 2018. Scikit-learn: Machine Learning in Python. http://arxiv.org/abs/1201.0490 (Accessed February 24, 2026).

Pletenev IA, Bazarevich M, Zagirova DR, Kononkova AD, Cherkasov AV, Efimova OI, Tiukacheva EA, Morozov KV, Ulianov KA, Komkov D, et al. 2024. Extensive long-range polycomb interactions and weak compartmentalization are hallmarks of human neuronal 3D genome. *Nucleic Acids Res* **52**: 6234–6252.

Rahman S, Dong P, Apontes P, Fernando MB, Kosoy R, Townsley KG, Girdhar K, Bendl J, Shao Z, Misir R, et al. 2023. Lineage specific 3D genome structure in the adult human brain and neurodevelopmental changes in the chromatin interactome. *Nucleic Acids Res* **51**: 11142–11161.

Ritchie ME, Phipson B, Wu D, Hu Y, Law CW, Shi W, Smyth GK. 2015. limma powers differential expression analyses for RNA-sequencing and microarray studies. *Nucleic Acids Res* **43**: e47.

Rizzardi LF, Hickey PF, DiBlasi VR, Tryggvadóttir R, Callahan CM, Idrizi A, Hansen KD, Feinberg AP. 2019. Neuronal brain region-specific DNA methylation and chromatin accessibility are associated with neuropsychiatric trait heritability. *Nat Neurosci* **22**: 307–316.

Soneson C, Love MI, Robinson MD. 2016. Differential analyses for RNA-seq: transcript-level estimates improve gene-level inferences. *F1000Research* **4**: 1521.

Sutton GJ, Poppe D, Simmons RK, Walsh K, Nawaz U, Lister R, Gagnon-Bartsch JA, Voineagu I. 2022. Comprehensive evaluation of deconvolution methods for human brain gene expression. *Nat Commun* **13**: 1358.

Tian W, Zhou J, Bartlett A, Zeng Q, Liu H, Castanon RG, Kenworthy M, Altshul J, Valadon C, Aldridge A, et al. 2023. Single-cell DNA methylation and 3D genome architecture in the human brain. *Science* **382**: eadf5357.

Uhlén M, Fagerberg L, Hallström BM, Lindskog C, Oksvold P, Mardinoglu A, Sivertsson Å, Kampf C, Sjöstedt E, Asplund A, et al. 2015. Tissue-based map of the human proteome. *Science* **347**: 1260419.

Virtanen P, Gommers R, Oliphant TE, Haberland M, Reddy T, Cournapeau D, Burovski E, Peterson P, Weckesser W, Bright J, et al. 2020. SciPy 1.0: fundamental algorithms for scientific computing in Python. *Nat Methods* **17**: 261–272.

Wang X, Park J, Susztak K, Zhang NR, Li M. 2019. Bulk tissue cell type deconvolution with multi-subject single-cell expression reference. *Nat Commun* **10**: 380.

Wightman DP, Jansen IE, Savage JE, Shadrin AA, Bahrami S, Holland D, Rongve A, Børte S, Winsvold BS, Drange OK, et al. 2021. A genome-wide association study with 1,126,563 individuals identifies new risk loci for Alzheimer's disease. *Nat Genet* **53**: 1276–1282.

Won H, de la Torre-Ubieta L, Stein JL, Parikshak NN, Huang J, Opland CK, Gandal M, Sutton GJ, Hormozdiari F, Lu D, et al. 2016. Chromosome conformation elucidates regulatory relationships in developing human brain. *Nature* **538**: 523–527.

Wu T, Hu E, Xu S, Chen M, Guo P, Dai Z, Feng T, Zhou L, Tang W, Zhan L, et al. 2021a. clusterProfiler 4.0: A universal enrichment tool for interpreting omics data. *Innov Camb Mass* **2**: 100141.

Wu W, Kargbo-Hill SE, Nathan WJ, Paiano J, Callen E, Wang D, Shinoda K, van Wietmarschen N, Colón-Mercado JM, Zong D, et al. 2021b. Neuronal enhancers are hotspots for DNA single-strand break repair. *Nature* **593**: 440–444.

Yu G, Wang L-G, Han Y, He Q-Y. 2012. clusterProfiler: an R package for comparing biological themes among gene clusters. *Omics J Integr Biol* **16**: 284–287.

Zaghi M, Banfi F, Massimino L, Volpin M, Bellini E, Brusco S, Merelli I, Barone C, Bruni M, Bossini L, et al. 2023. Balanced SET levels favor the correct enhancer repertoire during cell fate acquisition. *Nat Commun* **14**: 3212.

Zhang Y, Sloan SA, Clarke LE, Caneda C, Plaza CA, Blumenthal PD, Vogel H, Steinberg GK, Edwards MSB, Li G, et al. 2016. Purification and Characterization of Progenitor and Mature Human Astrocytes Reveals Transcriptional and Functional Differences with Mouse. *Neuron* **89**: 37–53. untington's disease pathology in human induced pluripotent stem cell-derived neurons. *Mol. Neurodegener.* **11**, 27 (2016).
